# It runs in the family: New extinct species from charismatic clades are more likely to be published in high-impact journals than those from non-charismatic clades

**DOI:** 10.64898/2026.10.06.756798

**Authors:** Bryan M. Gee

**Affiliations:** Burke Museum of Natural History & Culture, University of Washington, Seattle, WA, United States of America; University of Texas Libraries, University of Texas at Austin, Austin, TX, United States of America

## Abstract

The cultural landscape of paleontology is dominated by dinosaurs and a few other charismatic clades that are more likely to be depicted in media and to be recognized by the public. There is a prevailing belief among researchers that these biases extend to scholarly publishing – that certain clades enjoy a ‘taxonomic advantage’ that renders them more likely to be published in high-impact journals like *Nature* and *Science*. Given the importance of publishing in high-impact journals for career advancement, quantitative testing of this hypothesis can inform on potential biases in this area. I examined articles describing new species that were published between 2000 and mid-2026 in one of thirteen journals (*n* = 4,248 articles, 8,331 new species), ranging from high-profile, multidisciplinary titles like *Nature* to lower-profile, specialist titles like the *Journal of Paleontology*. Comparisons of the taxonomic composition of new species between journals revealed a positive correlation between a journal’s prestige (approximated by its impact factor) and the relative proportion of new species that belong to charismatic clades. Avemetatarsalians and mammaliamorphs represent more than 50% of new species in *Nature* and *Science* but less than 15% of new species in the control group of paleontological journals. These results provide support for the perception that there is a bias between extinct taxa as it relates to publication in high-impact journals. Given the potential for researchers to accrue cumulative advantage through publishing in prestigious journals, bias in favor of charismatic clades creates the potential for opportunities and knowledge generation to become increasingly skewed and consolidated towards these clades, at the expense of less charismatic clades and the people studying them. This threatens the discipline’s ability to continue to construct a holistic understanding of the full history of life that can be disseminated to the public. Further attention should be directed towards examining potential biases in paleontological publications as the primary currency in academia.

## Introduction

> *“Since Victorian times there has been considerable cachet associated with naming a new dinosaur species, and that is still true today. For example, a new species of dinosaur is more likely to grace the pages of* Nature *or* Science*, and to attract wide press interest, than a new species of almost any group other than fossil hominids.” – Benton (2010:1484)*

Some organisms are more popular than other organisms, among researchers and the general public alike. Despite representing minute fractions of the diversity of life and of the fossil record, dinosaurs (and a few other clades) have long-occupied a disproportionately large space in the cultural landscape of prehistoric life, as evidenced by their disproportionate representation in everything from movies to museum exhibits (e.g., Thomson, 2005; Sax, 2018; Rieppel, 2020; Chambers & McCahey, 2024; Manias, 2025; Boll, 2026). Benton’s remarks on the propensity of new dinosaurs to appear in prominent scholarly journals underscores a widespread belief among paleontologists that this disparity extends to academic research – that some (though hardly all) dinosaurs (and a few other clades) carry a particular cachet that confers a ‘taxonomic advantage’ in that research on charismatic clades is more likely to be published in such journals.

In general, charismatic taxa seem to garner disparate research attention, regardless of where related studies are published. For example, over the past decade, there more than 20 articles have been published with a major focus on spinosaurid ecology (mostly viz. *Spinosaurus*; e.g., Gimsa et al., 2016; Hendrickx et al., 2016; Vullo et al., 2016; Hone & Holtz, 2017, 2021; Aureliano et al., 2018; Hassler et al., 2018; Henderson, 2018; Amiot et al., 2020; Ibrahim et al., 2020a, 2020b; Schade et al., 2020; Arden et al., 2021; Beevor et al., 2021; Gimsa & Gimsa, 2021; Fabbri et al., 2022; Sereno et al., 2022, 2026; D’Amore et al., 2024; Smart & Sakamoto, 2024; Yun, 2024; Cau et al., 2026; Pittet, 2026; see also the book by Hone & Witton, 2026). Similarly, more than 20 articles have been published on tyrannosaurid ontogeny in the past decade (e.g., Persons et al., 2019; Peterson & Daus, 2019; Voris et al., 2019, 2022, 2025a; Carr, 2020; Woodward et al., 2020, 2026; Funston et al., 2021; Holtz, 2021; Rowe & Snively, 2021; Therrien et al., 2021, 2023; Yun et al., 2022, 2025; Coppock et al., 2024; Griffin et al., 2025; Zanno & Napoli, 2025; Longrich et al., 2026; Raun et al., 2026). Additionally, several of these studies were published in high-profile journals (e.g., Ibrahim et al. [2020a] and Zanno & Napoli [2025] in *Nature*; Griffin et al. [2025] and Sereno et al. [2026] in *Science*; Woodward et al. [2020] in *Science Advances*) and received extensive press coverage. Most other clades, either in their entirety or with respect to a specific attribute, are not as extensively studied or as high-profile in dissemination.

Disparity in public and scholarly interest is well-documented for extant taxa. Most “charismatic species” are large mammals (e.g., Ducarme et al., 2013; Colléony et al., 2017; Albert et al., 2018), even though such species are proportionately few in number and are not clearly more important in an objective ecological or evolutionary sense. This disparity, often called ‘taxonomic bias’ or ‘taxonomic chauvinism’ (e.g., Bonnet et al., 2002; Pawar, 2003), is widely documented between clades (e.g., Trimble & Van Aarde, 2010; Donaldson et al., 2016; Prokop et al., 2022; Tensen & Teske, 2025) and within clades (e.g., Fleming & Bateman, 2016; da Silva et al., 2020; dos Santos et al., 2020; Chen et al., 2023; Pollet et al., 2026). These biases are often discussed because they are a significant impediment to conservation of less charismatic species. However, negative impacts from disparate study are not restricted to conservation biology but can extend into other aspects of organismal biology, down to the knowledge of organisms’ mere existence, resulting in flaws including omission of taxa from studies, non-random/directional skewedness in a sample, and homogenization of a clade based on a few well-studied exemplars (e.g., Menge et al., 2009; Daru et al., 2018; Hughes et al., 2021; Šmíd, 2022; Simoncini et al., 2025; Anderson et al., 2026).

Conservation biologists have long suggested that taxonomic bias in conservation research is closely related to biases in societal interests (e.g., Seddon et al., 2005; Wilson et al., 2007; Ressurreição et al., 2012; Troudet et al., 2017; Davies et al., 2018; Jarić et al., 2019; Guedes et al., 2026). Given the high-profile nature of paleontology, it is reasonable to infer that the same correlation exists for the study of extinct organisms. Although paleontology writ large is not an applied science, there are still implications for the discipline that result from taxonomic biases in which extinct clades are studied and how they are studied in the same way that other anthropological biases (e.g., collection and sampling) are known to skew our interpretation of deep time (e.g., Davis & Pyenson, 2007; Carrasco, 2013; Irmis et al., 2013; Whitaker & Kimmig, 2020; Battista et al., 2023; Nanglu & Cullen, 2023; Dean & Thompson, 2025).

All clades merit research attention in order to produce a more holistic and objectively comprehensive understanding of the history of life. Potential taxonomic biases inhibit our ability to achieve such an understanding and can be situated within the broader taxonomic impediment in which descriptive and taxonomic work is increasingly undervalued by research institutions, funders, and other scholarly stakeholders. It is likely that this work and other types of research are not equally valued (or devalued) across all clades. Disparate valuation in turn has implications for how paleontological research is conducted and who is conducting it. For example, Gayford et al. (2024) remarked on potential pressure for researchers to overestimate the size of large extinct organisms in order to increase the appeal of manuscripts for consideration by higher-profile journals like *Nature* and *Science*, to obtain funding, and to attract attention from the public. As discussed in that study, a number of undoubtedly large taxa that were originally estimated to be exceptionally large sizes and that were first described in high-profile journals, such the dicynodont *Lisowicia bojani* Sulej & Niedźwiedzki, 2019 (*Science*) and the cetacean *Perucetus colossus* Bianucci, Lambert, Urbina, Merella, Collareta, Bennion, Salas-Gismondi, Benites-Palomino, Post, de Muizon, Bosio, Di Celma, Malinverno, Pierantoni, Villa, Amson, 2023 (*Nature*), have had the original estimates challenged (Romano & Manucci, 2021; Motani & Pyenson, 2024; Paul & Larramendi, 2025). This reflects a broader growing tendency for researchers to use superlatives and descriptors that (sometimes needlessly or baselessly) convey a sense of exceptionalism (e.g., Vinkers et al., 2015; Scott & Jones, 2017; Mammola, 2020) in order to “sell” their work. Raja & Dunne (2022) have discussed how pressure to publish in prestigious journals can lead researchers to overlook ethical concerns related to fossils’ provenance and collection if they have attributes likely to make them attractive to such venues.

Pressure to publish in high-profile journals can be a strong motivator to direct research attention to specific clades and to present research in a specific fashion given the weight accorded to such publications. These publications can be career-defining in the current ecosystem in which academics are assessed via publishing metrics such as publication and citation count, indices (e.g., h-index), and journal attributes like impact factor (JIF) and quadrant (e.g., Walker et al., 2010; Quan et al., 2017; McKiernan et al., 2019; Niles et al., 2020; Rice et al., 2020) that are used as proxies for research quality, regardless of whether they are suitable as such (e.g., Seglen, 1997a, 1997b; Saha et al., 2003; Cheek et al., 2006; The PLoS Medicine Editors, 2006; Dempsey, 2009; Alberts, 2013; Brembs et al., 2013; Eyre-Walker & Stoletzki, 2013; Putirka et al., 2013; Bornmann & Haunschild, 2018; Aksnes et al., 2019; Brito & Navarro-Rodriguez, 2019; Chapman et al., 2019; Thelwall et al., 2023). Prestigious journals like *Nature* and *Science* maintain their standing by being highly selective in order to publish research deemed to be of the greatest import – *Nature* describes itself as “publishing the finest peer-reviewed research in all fields of science and technology,” while *Science* describes itself as seeking “to publish those papers that are most influential in their fields or across fields” (American Association for the Advancement of Science, n.d.b; Nature Portfolio, n.d.b).

These journals (and other high-profile titles) also cite relevance to, or recognition from, the broader public, in their scope of publishing, returning to the point that societal biases can lead to research and publishing biases. A dinosaur expert will likely be able to make a more compelling argument for public interest in their work than a dinoflagellate worker, even if the former is not more “significant” than the latter in an objective evolutionary or ecological sense, leading dinosaur research to appear more often in prestigious journals. Such potential biases are more than an epistemologically interesting observation. The combination of pervasive use of metrics like JIF for research assessment and the scarcity economy created by highly (and increasingly) selective publishers risks creating a self-reinforcing loop in which only certain charismatic study systems are routinely considered for publication in high-impact journals. In turn, given that even one publication in a leading journal can confer significant advantages in professional areas such as funding and job applications and in journal submissions, specialists of more charismatic clades can gain a cumulative advantage (i.e. the ‘Matthew Effect’; Merton, 1968, 1988; Petersen & Penner, 2014; Ebadi & Schiffauerova, 2016; Bol et al., 2018; Katchanov et al., 2023) over specialists of other clades. For example, articles published in higher-JIF journals receive more citations than equivalent articles in lower-JIF journals (e.g., Larivière & Gingras, 2010) for reasons unrelated to their intrinsic quality, such as an assumption that JIF equates to quality or an attempt to signal the quality of one’s own work through citations of articles in high-JIF journals (e.g., Brembs et al., 2013; Drivas & Kremmydas, 2020). Such disparities carry no obvious benefits but are detrimental to academia and the public alike in hindering our ability to produce a holistic, comprehensive understanding of the full Tree of Life.

In recent years, paleontologists have increasingly examined how paleontological research is and has been conducted, largely within a framework of ethics and colonialism (e.g., Cisneros et al., 2022; Monarez et al., 2022; Raja et al., 2022; Chacon-Baca et al., 2023; Kempf et al., 2023; Macário et al., 2025; Valenzuela-Toro et al., 2025). Less attention has been directed towards trends in publishing (bibliometrics) within the discipline (e.g., Palanivel & Baskaran, 2018; Raja & Dunne, 2022; Wang et al., 2023; 2024, 2025a, 2025b; Smith et al., 2024; Amao et al., 2025; Rull, 2025; da Silva et al., 2026; Price, 2026), and most of these studies have focused on who is conducting the research, rather than on what is being studied. However, given the taxon-centered focus of most paleontological work, it is important to examine how biases might present in scholarly publishing based on which taxonomic systems a given researcher focuses on. Benton’s (2010) remark on the cachet associated with dinosaurs reflects a widely-held belief among paleontologists that some clades enjoy a significant competitive advantage in publishing. However, to the best of my knowledge, this has never been directly quantitatively tested. This study aims to address this gap by quantifying taxonomic representation among ‘naming papers,’ publications that describe new species, in high-profile journals like *Nature* and *Science* in order to test for biases between extinct taxa.

## Methods

### Publication sampling

#### Research scope of examined publications

This study focuses specifically on naming papers for several reasons. Firstly, the volume of paleontological literature has greatly expanded as the discipline has expanded from one rooted primarily in descriptive anatomy and taxonomy to a much broader methodological toolkit (e.g., Pandolfi et al., 2020; Dillon et al., 2023; Abdelhady et al., 2024; Smith et al., 2025). Attempting to sample every paleontological paper, even within a relatively narrow timeframe, would therefore be an incredibly labor-intensive undertaking due to the volume of publications.

Secondly, as the discipline has grown, it has increasingly adopted practices and concepts from other disciplines, such as ecological studies of extant taxa. An increasing number of studies combine neontological and paleontological data (e.g., evo-devo, phylogenetics, morphometrics), which complicates efforts to determine whether a study is in-scope or not as many studies lie somewhere between the endpoints of ‘strictly paleontological’ and ‘not at all paleontological.’ Similarly, attempting to identify the focal taxa or taxon of a study is not always straightforward. Many studies may present data for multiple taxa, some very disparately related (e.g., descriptions of large assemblages), and this may include both extant and extinct taxa (e.g., biomechanical or histological work). Especially when taxa sampled in a single study belong to multiple higher clades, this can greatly complicate efforts to taxonomically categorize the study beyond a relatively uninformative overarching clade (e.g., Vertebrata).

Finally, the discoverability, or lack thereof, of relevant literature also poses a barrier. Although there are numerous widely-used bibliometric tools (e.g., Dimensions, Google Scholar, OpenAlex, Scopus, Web of Science), these come with various limitations that are well-documented (e.g., Falagas et al., 2008; Mongeon & Paul-Hus, 2016; Vera-Baceta et al., 2019; Maddi et al., 2025; Simard et al., 2025), including: subscription fees (e.g., Dimensions, Scopus, Web of Science); incomplete topical coverage (all platforms); gaps in non-English language publications (most common for Scopus and Web of Science); and indexing of only metadata versus full-text for non-OA articles (all). The last of these is particular salient given that focal search terms may not appear in public metadata, only in the full text, and many articles have no publicly available full-text. For example, Raja & Dunne’s (2022) analysis of the journals that publish the most paleontology research was based on searching Web of Science for ‘pal(a)eontology’ or ‘pal(a)eobiology’, but many paleontological articles may not have used either of those terms in publicly available metadata fields (e.g., keywords, title) or even in the entire article, especially if they were published in paleontology-specific journals. Additionally, paleontological terms may be used in a study that is not paleontological in nature (e.g., ‘fossil’ for fossil fuel studies); even ‘pal(a)eontology’ can be invoked in work that is entirely neontological or not even in a natural science context (e.g., ethnography).

Naming papers are an accessible subset of paleontological studies for several reasons. Firstly, the naming of extinct taxa routinely garners tremendous public interest. Even if there are many technical components in the naming paper, the naming component is readily comprehensible to a lay audience. Likely for this reason, naming papers appear fairly regularly in high-profile journals compared to other research that may lack the same public appeal or accessibility (e.g., purely phylogenetic work). Naming papers also appear in a wide range of other types of journals, making it possible to compare differences in taxonomic representation between them.

Secondly, because a nomenclatural act is involved, at least some of the perceived value of the paper is based on the higher taxonomy of the studied material. For example, although not all clades of dinosaurs are equally popular, a new dinosaur of any placement is likely to be to be of greater interest than a new dinoflagellate. The focal taxa or taxon is also more apparent in these papers than other types of studies.

Thirdly, because of standards for designating a new taxon (e.g., a new species should be accompanied by a formal designation such as ‘sp. nov.’ or ‘n. sp.’; ICZN, 1999), there are standardized search terms that can yield a lower rate of false positives than might occur when querying large databases for other types of paleontological work (e.g., general anatomical descriptions, phylogenetic analyses); outside of the references, designator terms should essentially never appear unless a novel taxon is being proposed. Even if other methods are used in a naming paper, the act of designating a novel taxon is unambiguous such that the encapsulating article is undoubtedly in-scope. Identifying in-scope studies can be more complicated for multi-method studies when examining other types of paleontological work where there may not be a singular focus (e.g., a study entirely dedicated to paleohistology versus one that presents only brief histological data as part of a multi-method study). The predicted use of standardized terms also partially ameliorates challenges with general discoverability of relevant literature in large indexers.

Finally, because the naming of a new taxon does not necessitate advanced analytical methods, there is a temporally longer and numerically larger sample of naming papers than for many other types of studies that require technical or conceptual advancements that have only become common within the past few years.

#### Scope of taxonomic acts

A wide range of taxonomic acts are published every year, not all of which involve the creation of novel names. While all taxonomic acts are important, some are more interesting or comprehensible, especially to the public. For example, the naming of a new genus and species from never-before-published material is intrinsically more novel and potentially of greater interest than reassigning an existing species to an existing genus or designating an existing taxon as a synonym of another (though there are exceptions when high-profile taxa are involved).

This study is restricted to new species for several reasons. Firstly, hundreds to thousands of new species are named each year, creating the same problem of scale as with attempting to sample all paleontological studies. Secondly, new species are typically higher-profile than many other naming acts (e.g., naming a new unranked clade) because of the role of the species as a basic/fundamental operational unit for scientists and the relative familiarity of this rank to the public. Thirdly, not all Linnaean ranks are consistently used across clades, particularly below the species level; for example, vertebrate paleontologists virtually never designate subgenera or subspecies, whereas this is more common in invertebrate paleontology. Fourthly, although the standardization of designations at Linnean ranks (e.g., ‘sp. nov.’, ‘gen. nov.’) is relatively high, this is less true for unranked clades, which are an increasingly common means of refining classifications when constrained between two Linnean ranks. Finally, although there are often recommended abbreviations for Linnean ranks (e.g., ‘sp. nov.’ or ‘n. sp.’ are the most common and recommended for species; ICZN, 1999; Braby, 2024; Ruedas et al., 2025), per ICZN Article 16.1, authors only need to make an explicit intention through some kind of expression that clearly delineates the intention. This not only means that other abbreviations (e.g., ‘sp. n.’, ‘sp. nov.’) are acceptable but also that unabbreviated terms such as “new family”, “new genus”, “new species”, and “new genus and species” are acceptable. The number of variations for any single rank complicates searching (e.g., most search engines limit the number of terms in a Boolean search), so constraining the focus to one rank was deemed the most efficient.

#### Journal scope of examined publications

Sampling all new extinct species across every known journal would be extremely labor-intensive; for example, Wang et al. (2023) reported over 2,600 new vertebrates between 2017 and 2021 alone, and vertebrates make up a relatively small fraction of novel extinct species. A second limitation as it pertains to this study’s aim of examining patterns between journals based on profile is that not all journals have a JIF, which is a widely-used proxy for the relative profile of a journal. This study therefore samples thirteen journals divided into three groups (Table 1).

**Table 1.** List of journals examined in this study. All journal impact factors (JIF) are for 2025. *The *Papers in Palaeontology* sampling does not include pre-2014 publication under the *Special Papers in Palaeontology* serialization when it emphasized taxon-focused or thematic monographs because the monograph model is more atypical (though a separate examination of monographs could be warranted).

| Journal | Society/Museum | Publisher | Since | JIF |
| --- | --- | --- | --- | --- |
| **High-JIF** |  |  |  |  |
| *Nature* | None | Springer Nature | 1869 | 56.1 |
| *Science* | American Association for the Advancement of Science | American Association for the Advancement of Science | 1880 | 47.3 |
| **Medium-JIF** |  |  |  |  |
| *Current Biology* | None | Cell Press (Elsevier) | 1991 | 7.7 |
| *Gondwana Research* | International Association for Gondwana Research | Elsevier | 1997 | 8.9 |
| *Nature Communications* | None | Springer Nature | 2010 | 18.1 |
| *Nature Ecology & Evolution* | None | Springer Nature | 2017 | 17.1 |
| *Proceedings of the National Academy of Sciences of the United States of America* | National Academy of Sciences of the United States | National Academy of Sciences of the United States | 1915 | 9.5 |
| *Science Advances* | American Association for the Advancement of Science | American Association for the Advancement of Science | 2015 | 13.9 |
| **Control (Low-JIF)** |  |  |  |  |
| *Journal of Paleontology* | Paleontological Society | Cambridge University Press | 1927 | 1.3 |
| *Journal of Systematic Palaeontology* | Natural History Museum London | Taylor & Francis | 2003 | 2.9 |
| *Journal of Vertebrate Paleontology* | Society of Vertebrate Paleontology | Taylor & Francis | 1981 | 1.9 |
| *Palaeontology* | Palaeontological Association | Wiley-Blackwell | 1957 | 3.2 |
| *Papers in Palaeontology* | Palaeontological Association | Wiley-Blackwell | 2014* | 2.4 |

The first group of journals are multidisciplinary (generalist) ones with extremely high JIFs that are known to be common venues for paleontological work in general and naming papers specifically. This includes *Nature* (JIF = 56.1) and *Science* (47.3) as the two highest JIF venues that paleontologists routinely publish in (also reported by Raja & Dunne, 2022).

The second group of journals have moderately high JIFs and may have narrower scopes, although they are still much broader than paleontological journals: *Current Biology* (7.7), *Gondwana Research* [8.9], *Nature Ecology & Evolution* (17.1), *Nature Communications* (18.1), *Proceedings of the National Academy of Sciences of the United States* (hereafter ‘*PNAS*’; 9.5), and *Science Advances* (13.9). Some high-JIF journals in which paleontological research appears but naming papers are rare to non-existent (e.g., *Biological Reviews* [10.9], *Earth-Science Reviews* [11.3], *PLOS Biology* [6.9], *Systematic Biology* [7.4]) are omitted.

The final group of relatively low-JIF, paleontology-specific journals acts as the ‘control group’ against which the other two groups are tested: *Journal of Vertebrate Paleontology* (for vertebrates only; 1.9), *Journal of Systematic Paleontology* (2.9), *Journal of Paleontology* (1.3), *Palaeontology* (3.2), and *Papers in Palaeontology* (2.4). These journals were selected for several reasons. Firstly, all of these are paleontological journals, rather than broad-scope biology or earth science journals that accept paleontological contributions (e.g., *Journal of Anatomy; Proceedings B of the Royal Society of London*; *Zoological Journal of the Linnean Society*) or completely generalist journals (e.g., *PeerJ*, *PLOS ONE*, *Royal Society Open Science*, *Scientific Reports*).

Secondly, with the exception of the *Journal of Vertebrate Paleontology* (vertebrates only) and the *Journal of Systematic Palaeontology* (must have a systematic scope), none of the control-group journals have any taxonomic, geographic, institutional, temporal, or topical restrictions on scope, and all accept, and have routinely published, taxonomic work. This is in contrast to other paleontological journals that have narrower taxonomic (e.g., *Journal of Foraminiferal Research*; *Marine Micropaleontology*; *Palaeoentomology*), temporal (e.g., *Cretaceous Research*), or topical focuses (e.g., *Palaeogeography, Palaeoclimatology, Palaeoecology*; *Paleobiology*); *Gondwana Research*, despite the name, does not have a geographic restriction. The *Journal of Vertebrate Paleontology* is included because it is used as a benchmark for vertebrates specifically, which I hypothesize are typically more charismatic and high-profile than invertebrates or non-metazoans based on anecdotal experience and the conservation biology literature (e.g., Donaldson et al., 2016). This journal is also a well-known and widely-used venue for naming new species. The *Journal of Systematic Palaeontology* is included since naming papers are a component of systematic work, so the fact that the journal has a topical focus facilitates, rather than confounds, the aims of this study.

Thirdly, all of these journals are affiliated with a scholarly entity (society or museum) and are well-known among the paleontological community. They tend to publish relatively large numbers of articles compared to other paleontological journals (e.g., Raja & Dunne, 2022). They have traditionally been free to publish in, making them more accessible than gold OA journals (e.g., MDPI’s *Fossil Studies*), and do not require society or institutional affiliation (e.g., some museum publications like *Bulletin of the American Museum of Natural History*). However, these attributes do not eliminate all biases (e.g., all are published by western societies and require English-language publication). All of these journals are also published under hybrid OA models, which are less accessible than diamond OA titles (e.g., *Acta Palaeontologica Polonica*; *Geodiversitas*; *Open Palaeontology*; *Palaeodiversity*; *Palaeontologia Electronica*; *Revista Brasileira de Paleontologia*; *Vertebrate Anatomy Morphology Palaeontology*).

Diamond OA journals are not sampled because most of them have very low JIFs, which can make them less desirable in the current ecosystem in which JIF is widely-used (e.g., Walker et al., 2010; McKiernan et al., 2019; Demeter & Istratii, 2020; Niles et al., 2020; Kumari & A, 2025). Additionally, across all disciplines, they are far more common outside of North America and western Europe (the two regions where the major commercial publishers are based and where the majority of scholarship is generated; e.g., Bosman et al., 2021).

Two other ‘types’ of journals are excluded: lower-profile, non-paleontological society or association journals (semi-arbitrarily defined as journals with JIF < 4.0; e.g., *Biology Letters*, *Proceedings B of the Royal Society of London*, *The Anatomical Record*, *Zoological Journal of the Linnean Society*) and high-volume gold OA journals, often called open-access mega-journals (OAMJ; e.g., *PeerJ*, *PLOS ONE*, *Scientific Reports*, journals in the MPDI and Frontiers Media portfolios). The former are excluded because they are not considered high-profile for the purposes of this study and because they are not paleontology-specific. The latter are excluded for those reasons and because the ‘soundness only’ model of some OAMJs (i.e. that the journal only seeks to publish work deemed sound, rather than restricting it to work that is deemed significant or sufficiently novel; e.g., Wakeling et al., 2016; Erfanmanesh & Teixeira da Silva, 2019) could introduce a confound into an analysis examining how journals might use taxonomic charisma as an editorial/reviewing criterion.

#### Search strategy

Centralized indexers like Dimensions, Google Scholar, OpenAlex, Scopus, and Web of Science are commonly used in bibliometric or scientometric analyses. However, this study opts not to use these for several reasons, most of which have been previously mentioned, such as problems with coverage and inclusion (e.g., Mongeon & Paul-Hus, 2016; Vera-Baceta et al., 2019; Asubiaro et al., 2024) and their inability to search full-text versions of paywalled articles, even though key search terms may only appear there (e.g., Lin, 2009). Additionally, analyses done with proprietary tools like Scopus and Web of Science are limited in reproducibility because they are subscription-based, and many researchers lack access to these closed-source, high-cost tools – annual subscription fees for institutional use of these platforms can exceed $200,000 USD (e.g., Brundy & Thornton, 2024). Due to costs and to closed-source designs, they run counter to the aspirations of the open science movement. Google Scholar is a freely available (but closed-source) solution that is mainly impeded by a near total absence of search filters, including some that would be essential for this study (e.g., filter by journal or publisher). OpenAlex is a newer open-source solution with extensive filtering capabilities that sometimes outperforms commercial solutions in terms of coverage (e.g., Céspedes et al., 2025; Culbert et al., 2025; Maddi et al., 2025; Simard et al., 2025). However, it shares the full-text restriction of other platforms, leaving it susceptible to limited discoverability.

For hybrid-OA journals, which include high-JIF titles like *Nature* and *Science*, the most reliable way to ensure comprehensive full-text searching is through the publisher’s own interface. Therefore, an identical search query was made in each journal’s web browser search, which captures the full-text of non-OA works and older works that do not have HTML representations. It should be acknowledged that these search interfaces are also closed-source systems, so silent changes to content (e.g., digitization of older publications) and database structure (e.g., metadata tags) may lead to different results from a query made at a different time. Boolean searches that include numerous taxonomic abbreviations (e.g., ‘sp. nov.’, ‘n. sp.’) were constructed; this includes queries for novel ichnospecies and oospecies as well, although these are a minority in all sampled journals. Several non-abbreviated terms (e.g., ‘new species’) were included to mitigate the inability to identify every conceivable designating phrase. For non-paleontology journals, the search terms were combined with a requirement that the article also contain at least one of the terms ‘fossil,’ ‘paleo*,’ or ‘palaeo*,’ where the ‘*’ denotes a wildcard (e.g., ‘paleo*’ captures ‘paleontology,’ ‘paleoanthropology,’ ‘paleoecology’, etc.); it is possible that a naming paper did not use any of these expected terms, but this would be expected to be extremely rare because the search covers all text fields, including the references, where at least one of these terms is highly likely to appear, even if they do not appear in the main-text. Search results were restricted to 2000 to 2026, inclusive. Data were collected between March and September 2026, with a final check on all journals performed on September 4, 2026 to ensure a consistent cutoff date. The search queries are provided in the supporting appendices (Gee, 2026c).

#### Article inclusion

Only standard research articles were included. Other types of articles, such as editorials, opinions, reviews, comments, replies, notes, and errata/corrections were excluded; these classifications are based on formal article categories in the journal, not on a personal set of criteria. Articles where a replacement name is proposed (either genus or species in this case), regardless of where the original name was published, are excluded because these are an atypical type of article and do not represent a fully distinct work. I have endeavored to use the replacement name in the analysis when one could be efficiently identified (e.g., when the replacement name was published in the same journal and the same year as the original name).

Taxa that are presently considered junior synonyms of others are retained since they were considered valid at the time of original publication. If a naming publication was retracted (e.g., Christiansen & Mazak, 2009; Xing et al., 2020), the publication and its novel name(s) are retained in the dataset since the retraction has no direct bearing on the perceived value of the work at the time of original publication.

### Data collection

#### Manual collection of article and novel species metadata

Given the focus of this study, the data that were manually collected from each naming paper are narrow in scope: journal name, article title, publication year, the DOI, the binomial of the new species, the country or equivalent geographic unit (e.g., ocean), and the erathem and system per the International Commission on Stratigraphy (ICS). Cenozoic records also have the series recorded. For articles describing multiple new species, each species is recorded as a separate entry with identical article metadata. Species had to be formally named with a binomial to be recorded; the practice of “open nomenclature” (e.g., *Genus* sp. nov., *Genus* sp. A; see Zeng et al., 2026, for a recent example) does not formally establish a name, and these were not recorded.

#### Programmatic collection of article metadata

Additional article metadata were retrieved through a script for two secondary purposes. The first is a cursory examination of the frequency of repeat authors in high-JIF journals. Each article DOI was queried through the OpenAlex API to extract author names and ORCIDs; the number of times a unique ORCID occurred was then used to estimate repeat authorship, as this provides a more unambiguous unique count than name matching alone (e.g., Haak et al., 2012).

The second is the automated generation of a reference list for all naming papers in the dataset, regardless of whether they are cited in the main text, in order to ensure proper citation credit for primary data generators (e.g., Smith et al., 2024; Jones et al., 2025). Each article DOI was queried through the Crossref API, with standard citation metadata extracted (e.g., author list, volume). Year of print publication is used over year of online print when an article has multiple media, so the listed year of a naming paper might differ from that listed in the dataset if the electronic version and the print version were not published in the same year. Only minor cleaning steps were performed (e.g., title-casing author names, replacing HTML tags for italics with italics formatting). Due to the very large number of references, individual references were not vetted, so formatting inconsistencies such as titles in all-caps or lack of italicization of genera and species are due to how journals submitted metadata to Crossref when minting the DOI; these should not impact indexers’ ability to identify the article.

### Data processing

Data were manually entered in a spreadsheet using Google Sheets, then exported to CSV format for processing using a series of Python scripts (Gee, 2026a) that make use of the following modules *csv*, *datetime*, *glob*, *json*, *matplotlib* (Hunter & Droettboom, 2026), *numpy* (Oliphant, 2006), *os*, *pandas* (McKinney, 2010; The pandas development team, 2026), *pyalex* (De Bruin, 2026), *pypandoc-binary* (Tegner, 2026), *re*, *requests* (Reitz, 2025), *rich* (McGugan, 2026), *scipy* (Virtanen et al., 2020), and *time*. Columns were cleaned for excess whitespace, and the genus was extracted from the binomial name as a new column. For genera that had already been identified as having replacement names, the name is updated in this genus column in the scripts.

#### Retrieval of taxonomic rank information

Each genus was then queried through the Paleobiology Database (PBDB) API to retrieve higher taxonomic ranks on September 8, 2026. While I am well-aware of the limitations of the PBDB regarding completeness and current taxonomy, these limitations are intrinsic to any large database, and the PBDB was considered to be the best available resource for semi-standardized taxonomic metadata across the entire Tree of Life and across standard Linnaean ranks. Although some rank information is provided in naming papers, this is rarely done for all possible ranks, and various higher clades may go into dis-use or be reclassified over time. Thus, manual recording of the original taxonomy provided in the naming papers would still necessitate extensive augmentation and standardization within some framework. The relationships of many taxa remain without consensus such that any large-scale framework will be criticized in some areas. Because this analysis is focused on high-level clades (e.g., are dinosaurs more commonly described in *Nature* than arthropods), I believe the potential impacts of instability at lower ranks where there may be greater taxonomic ambiguity or debate (e.g., family and subfamily) are minimal (e.g., Wagner et al., 2007). For example, the placement of turtles and the relationship of insects to crustaceans have no impact on the analyses.

Taxonomic rank information was programmatically retrieved through the PBDB API for the majority of genera. However, 1,012 species belonging to 861 unique genera did not have any rank information retrieved for them; this represents 12.14% of all species and 13.21% of all genera in the dataset. All unique genera were further manually examined for manual augmentation. A small proportion of these (39 species [0.46%], 32 genera [0.49%]) were missing data due to API connection failures or typographic errors in data entry and should have been retrieved. Another small portion are listed in the PBDB, but at the subgenus rank, not the genus rank (13 species [0.15%], 12 genera [0.18%]); one is listed only as a junior synonym at the genus level. All remaining deficiencies (965 species [11.58%], 823 genera [12.63%] are considered “true” deficiencies; note that some were added very shortly after data collection (not by myself). For missing genera, rank information was manually added by querying progressively higher ranks that were listed for a new species in its naming paper until a taxon was found in the PBDB (e.g., if a genus was not in the PBDB, but its originally listed family was, the full PBDB taxonomy down through the family level was populated for that species). If a genus that had to be manually entered was listed for multiple articles, the taxonomy from the most recent publication was used.

#### Taxonomic ranks cleaning processes

Given the scale of the dataset (>8,300 new species), the frequent lack of consensus about the placement of various taxa (at all ranks), and the emphasis on higher ranks, I did not systematically review all species to see whether their placement had changed; it is inevitable that at least some are still debated today. It is rare for taxa to have been originally identified to a different high-level rank (e.g., superfamily or order) than they are presently classified. Some examples include *Oculudentavis khaungraae* Xing, O’Connor, Schmitz, Chiappe, McKellar, Yi, & Li, 2020 [retracted]), originally described as a bird and now considered to be a squamate (Bolet et al., 2020, 2021; Li et al., 2020, 2021) and ‘*Galeaplumosus abilus*’ Hou, Aldridge, Siveter, Siveter, Williams, Zalasiewicz, & Ma, 2011, originally described as a hemichordate before it was recognized to be part of the same organism as the ctenophore *Xianguangia sinica* Chen & Erdtmann, 1991, by Ou et al. (2017). Given the unusual circumstances for both species, the taxonomic ranks listed in the dataset reflect the original classification, not the current one, since the original interpretation is the context in which the species was considered for publication; for *Oculudentavis* Xing, O’Connor, Schmitz, Chiappe, McKellar, Yi, & Li, 2020 [retracted]), this information was only changed for the type species, *O. khaungraae*, since the second named species has always been classified as a squamate.

When a genus of contested higher placement was not found in the PBDB, the manually added taxonomy is based on the original description unless there is unambiguous consensus around revised placement (as in the above examples). For, example *Afrocascudo saharaensis* Brito, Dutheil, Gueriau, Keith, Carnevale, Britto, Meunier, Khalloufi, King, de Amorim, & Costa, 2024, was originally identified as the oldest known catfish, but subsequently suggested to be a lepisosteiform by Britz et al. (2024). Because I do not have comprehensive knowledge of every clade, there may be other instances like these that were not adjusted, but I would expect that they are exceedingly rare, at least at the fairly coarse taxonomic levels of analysis performed here.

Beyond these taxa, a small number of changes were made to insert ranks for single genera (noted in the dataset). During the categorization of species (see next section), some overt errors were noted (e.g., Squamata not being listed under Lepidosauria), and ranks were added manually here. The two other instances in which missing ranks were manually added for an entire higher-level taxon were the listing of Tritylodontidae under Mammaliamorpha and the listing of Foraminifera under Retaria. Theropoda, listed directly under Dinosauria in the PBDB, was placed under Saurischia; this placement only affects a few secondary analyses.

As previously mentioned, I endeavored to use replacement names where appropriate; some replacement names were easily detected because they are published in the same journal as the original name; replacement name papers were frequently retrieved in journal searches. However, because of different avenues for formally replacing a name (e.g., many journals do not publish replacement name articles or may do so only as a corrigendum), there could be a small number of remaining homonyms that cannot be identified without manually performing a literature search for every genus. The PBDB sometimes lists the replacement name and records the relationship to the original name; sometimes lists the replacement name but without the relationship (the taxonomy between the two may or may not be essentially identical); and sometimes does not include the replacement name. Unlike when a name remains valid but the taxonomy has drastically changed, the taxonomy for all genera with replacement names was updated to the current PBDB taxonomy. This avoids issues where, for example, a genus name originally created for a vertebrate holotype that is still classified as a vertebrate is found to be a junior homonym of a previously named invertebrate since homonymy is unrelated to taxonomic placement of a genus. Species names found to be homonyms were also edited in the dataset, but the genus was unchanged for all identified instances, so the taxonomic rank information remained unchanged.

Some types of fossils have other systems of classification (e.g., ‘morphospecies’). These were adjusted to conform to the most conservative Linnaean taxonomy for the inferred source. Most ichnotaxa entered in the PBDB list ‘Ichnofossils’ as the highest taxonomic rank (instead of ‘Life’); these were edited to use the taxonomy of the inferred tracemaker (*n* = 22).

Finally, all taxa for which data were retrieved from the PBDB but two or fewer ranks were listed were manually examined, as this usually represents a lack of nesting of a taxon under a parent taxon (e.g., many of these were missing the typical ‘Life’ rank); 73 genera comprising 87 species taxa were edited, the majority of which were for a lack of rank information other than genus when additional ranks could be confidently added based on the original naming paper.

#### Temporal and geographic data cleaning processes

As part of the quality control measures, any entry that did not have a chronostratigraphic system or geographic provenance originally recorded during data collection was re-examined to ensure that information had not been missed. For entries where this information had been collected, the listed time bins (or lack thereof) were not reviewed to see if this metadata had changed since the temporal categorization used in the analyses is relatively coarse; it is more common for fossil-bearing units to be re-dated to a different series (e.g., Induan to Olenekian), which is not analyzed here, than to a different system (e.g., Jurassic to Cretaceous), unless the locality is close to a boundary. Likewise, originally identified geographic provenance was also not reviewed since this was considered extremely unlikely to change; if there was uncertainty about the provenance in the initial data entry, this field was left blank.

#### Categorization of genera

Because the dataset includes taxa from across the entire Tree of Life, the most common clades vary significantly by journal. Furthermore, it is not possible to effectively visualize all possible lower ranks (e.g., families), which number in the hundreds to thousands. Therefore, I categorized all taxa based on their rank hierarchies into one of 20 categories (one of which is a catch-all ‘Other’; Table 2). These categories attempt to capture groupings that are both charismatic and well-represented, with a preference for more charismatic but species-poor groupings over less charismatic but species-rich groupings, while keeping the ‘Other’ bin to less than 10% of the total sample. It should be noted that ‘charisma’ is not quantified here as is sometimes done with extant taxa based on either measures like body mass (e.g., Berti et al., 2020; Prokop et al., 2022) that can only be estimated at best for extinct taxa or public surveys (e.g., Ducarme et al., 2013). Instead, the operational definition follows general observations for extant taxa in which taxa like birds (including dinosaurs here) and mammals are more charismatic than other vertebrates like amphibians and fish, and vertebrates are more charismatic than non-vertebrates; anecdotally, in both a research and a broader societal context, these generally hold true for extinct clades as well (e.g., theropods are more popular than temnospondyls). Undoubtedly, there is further nuance within this (e.g., not all mammals are equally charismatic, some taxa in relatively uncharismatic clades are extremely charismatic).

**Table 2.**
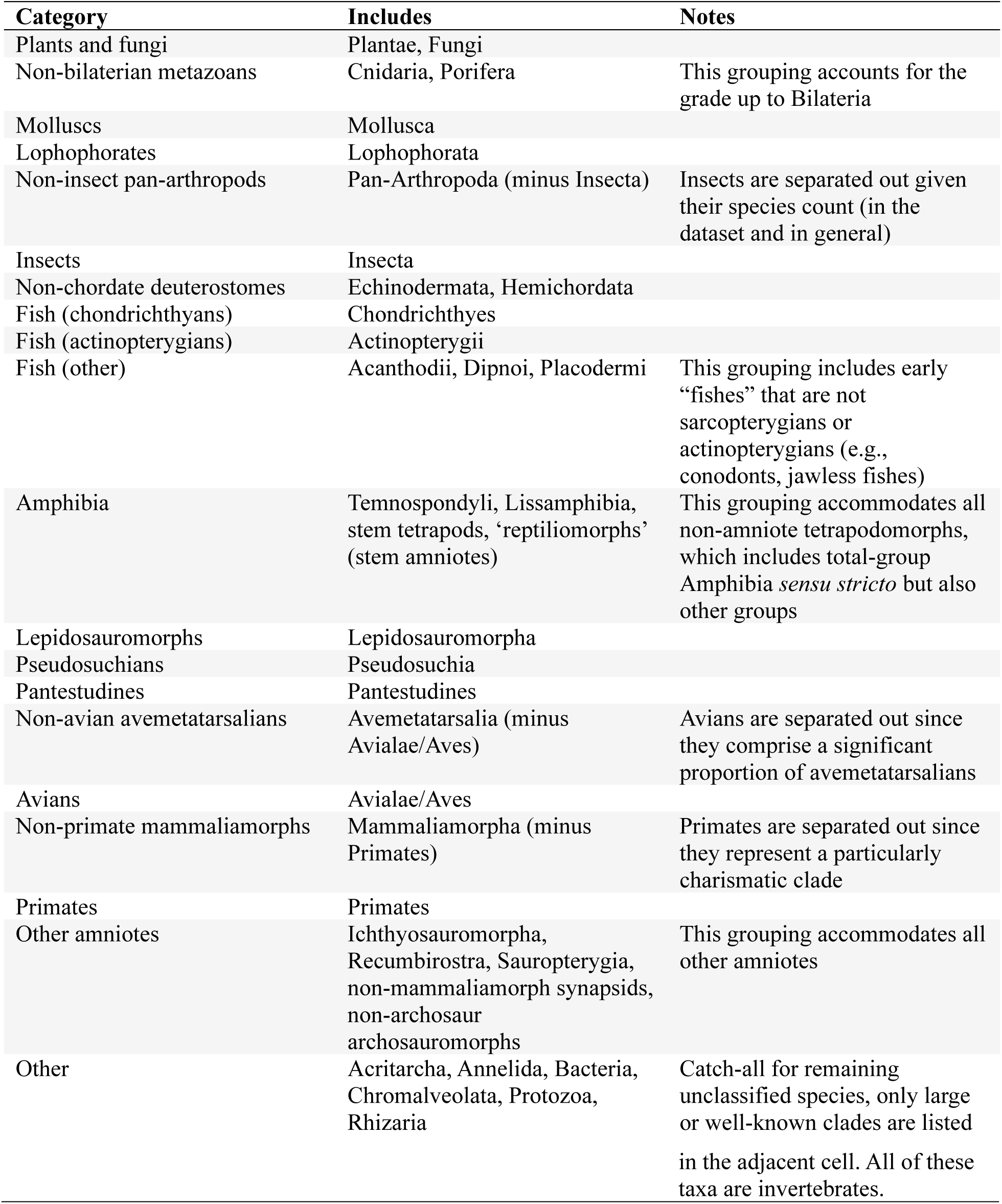
List of 20 categories into which all species were grouped for analysis.

This approach explains why, for example, Pantestudines, which includes 95 species in this dataset, is a distinct category, whereas Echinodermata, which includes more than 400 species, is not a separate category. For the same reason, the catch-all ‘Other’ includes some fairly well-known non-vertebrate clades (e.g., Annelida); their inclusion in this category should not be misconstrued as anything more than a decision borne out of logistical constraints. These groupings do not always represent monophyletic clades (e.g., non-chordate deuterostomes; plants and fungi), and not all groupings that do represent clades are at the same Linnean rank (e.g., primates vs. pseudosuchians).

## Results

### Full dataset

The full dataset represents 8,331 unique species, distributed among 6,515 unique genera and published across 4,248 unique articles (a total of 6,027 articles were examined for the study). The overwhelming majority of these are from the control group (Table 3).

**Table 3.** Summary of counts for articles retrieved and retained and counts of genera and species analyzed. *One additional paper was manually added after a quality control check on replacement names returned the replacement name paper but not the original description (this is noted in the data file). The total number of assessed articles for this journal is thus 1,491. All other counts for this journal include the manually added article.

| Journal | Articles retrieved | Articles retained | Genera | Species |
| --- | --- | --- | --- | --- |
| *Nature* | 280 | 186 | 199 | 205 |
| *Science* | 122 | 56 | 65 | 70 |
| *Current Biology* | 277 | 82 | 88 | 90 |
| *Gondwana Research* | 145 | 64 | 91 | 102 |
| *Nature Communications* | 146 | 49 | 71 | 74 |
| *Nature Ecology & Evolution* | 126 | 24 | 28 | 28 |
| *Proceedings of the National Academy of Sciences* | 252 | 70 | 84 | 87 |
| *Science Advances* | 46 | 15 | 17 | 18 |
| *Journal of Vertebrate Paleontology* | 1,490* | 1,032 | 1,134 | 1,129 |
| *Journal of Paleontology* | 1,298 | 1,291 | 2,385 | 3,167 |
| *Palaeontology* | 848 | 584 | 971 | 1,171 |
| *Papers in Palaeontology* | 354 | 284 | 600 | 727 |
| *Journal of Systematic Palaeontology* | 642 | 511 | 1,072 | 1,353 |

#### Taxonomic categories

Some brief summary metrics are listed below for the dataset in its entirety (all percentages are out of the 8,331 total). Figure 1 visualizes the proportion of new species that each of the 20 categories represents, with non-insect pan-arthropods, molluscs, and non-primate mammaliamorphs being the most common and pseudosuchians, turtles, and primates being the least common. Table 4 summarizes the most common category as well as the most common age-category and age-geography-category combinations per journal.

**Figure 1.**
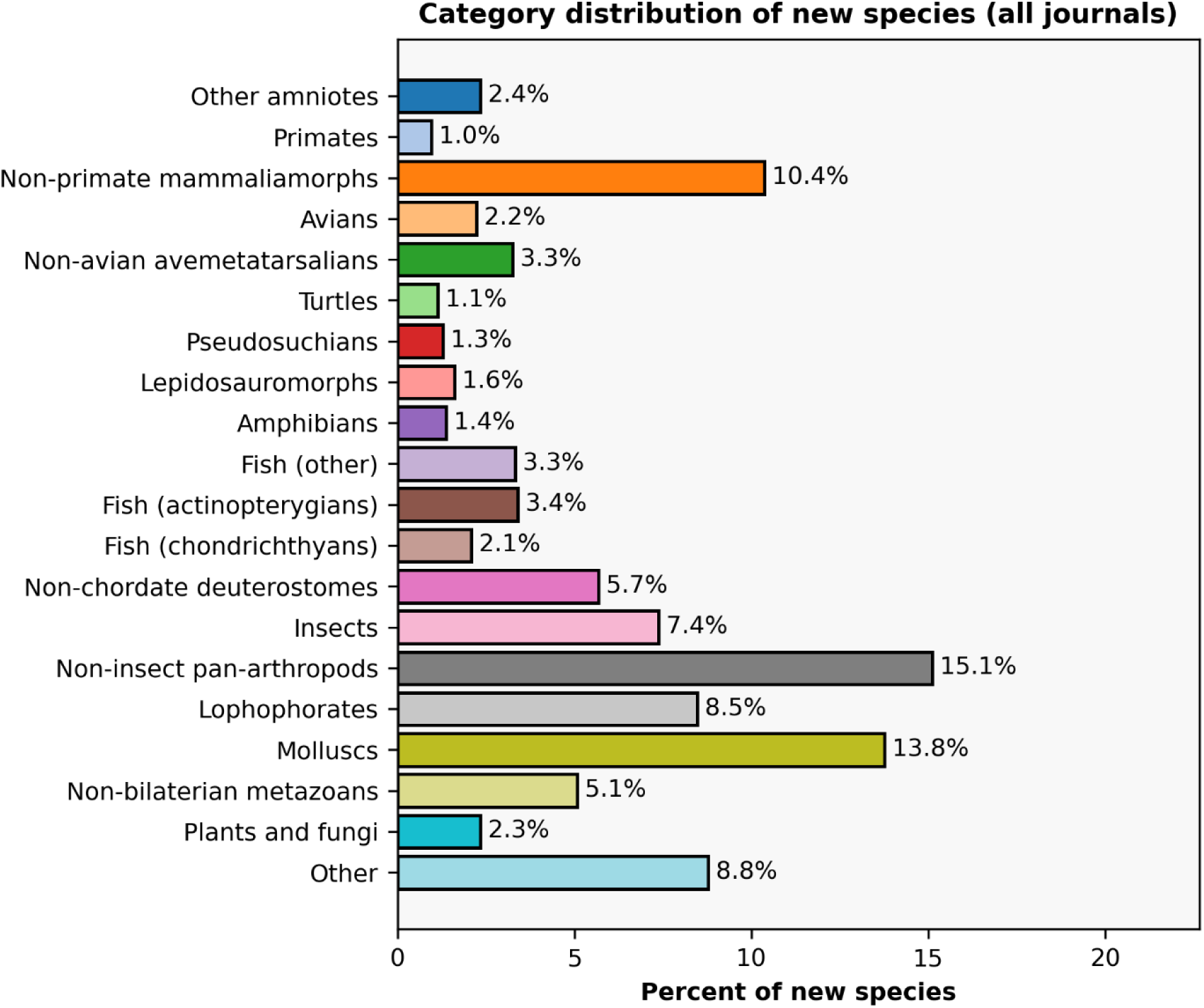
Distribution of new species by taxonomic categorization. Categories are roughly arranged in order of phylogenetic divergence, with invertebrates at the bottom, fish and non-amniotes in the middle, and amniotes at the top. This color scheme is applied consistently across all figures showing this taxonomic breakdown (e.g., insects are always a light pink bar).

**Table 4.**
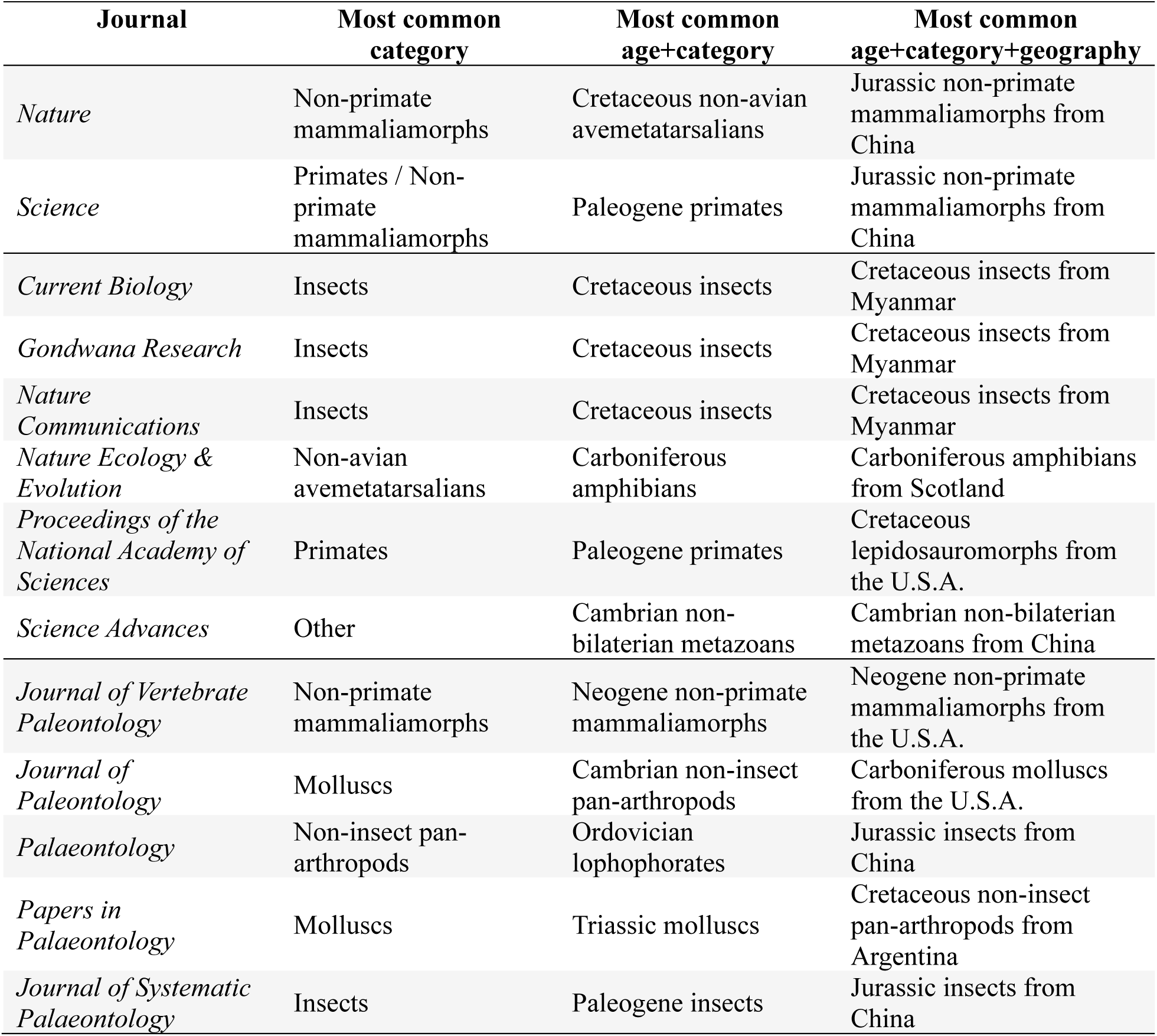
Summary of the most common taxonomic category for each journal sampled here. Journals are divided by the three groups used in this study. ‘*Most common category*’ represents the most common of the 21 categories; ‘*age+category*’ and ‘*age+category+geography*’ represent the most common two- or three-variable combination for each journal; these will not always be the same (e.g., *Nature*).

| Journal | Most common category | Most common age+category | Most common age+category+geography |
| --- | --- | --- | --- |
| *Nature* | Non-primate mammaliamorphs | Cretaceous non-avian avemetatarsalians | Jurassic non-primate mammaliamorphs from China |
| *Science* | Primates / Non-primate mammaliamorphs | Paleogene primates | Jurassic non-primate mammaliamorphs from China |
| *Current Biology* | Insects | Cretaceous insects | Cretaceous insects from Myanmar |
| *Gondwana Research* | Insects | Cretaceous insects | Cretaceous insects from Myanmar |
| *Nature Communications* | Insects | Cretaceous insects | Cretaceous insects from Myanmar |
| *Nature Ecology & Evolution* | Non-avian avemetatarsalians | Carboniferous amphibians | Carboniferous amphibians from Scotland |
| *Proceedings of the National Academy of Sciences* | Primates | Paleogene primates | Cretaceous lepidosauromorphs from the U.S.A. |
| *Science Advances* | Other | Cambrian non-bilaterian metazoans | Cambrian non-bilaterian metazoans from China |
| *Journal of Vertebrate Paleontology* | Non-primate mammaliamorphs | Neogene non-primate mammaliamorphs | Neogene non-primate mammaliamorphs from the U.S.A. |
| *Journal of Paleontology* | Molluscs | Cambrian non-insect pan-arthropods | Carboniferous molluscs from the U.S.A. |
| *Palaeontology* | Non-insect pan-arthropods | Ordovician lophophorates | Jurassic insects from China |
| *Papers in Palaeontology* | Molluscs | Triassic molluscs | Cretaceous non-insect pan-arthropods from Argentina |
| *Journal of Systematic Palaeontology* | Insects | Paleogene insects | Jurassic insects from China |

- 8,280 (99.35%) are classified under Eukaryota.
- 7,535 (90.41%) are classified under Animalia.
- 3,832 (45.98%) are classified under Protostomia, while 3,275 (39.30%) are classified under Deuterostomia.
- 2,802 (33.62%) are classified under Chordata; 2,648 (31.77%) are classified under Vertebrata.
- 1,935 (23.22%) are classified under Amniota, of which 412 (4.93%) are classified under Dinosauria (inclusive of avians), and 923 (11.08%) are classified under Mammalia.

#### Spatiotemporal categories

Temporally, the dataset is dominated by taxa from the Cretaceous, Neogene, and Paleogene; collectively, these represent 41.3% of all new species (Fig. 2; Table 5). Spatially, the dataset is dominated by taxa from just five countries, the United States of America, China, Canada, Argentina, and Australia (all >3% of the total dataset; Fig. 3; Table 5); collectively, these account for 47.5% of all new species, even there are 138 unique geographic entities in the dataset (most are countries or territories, although some oceans are included as well).

**Figure 2.**
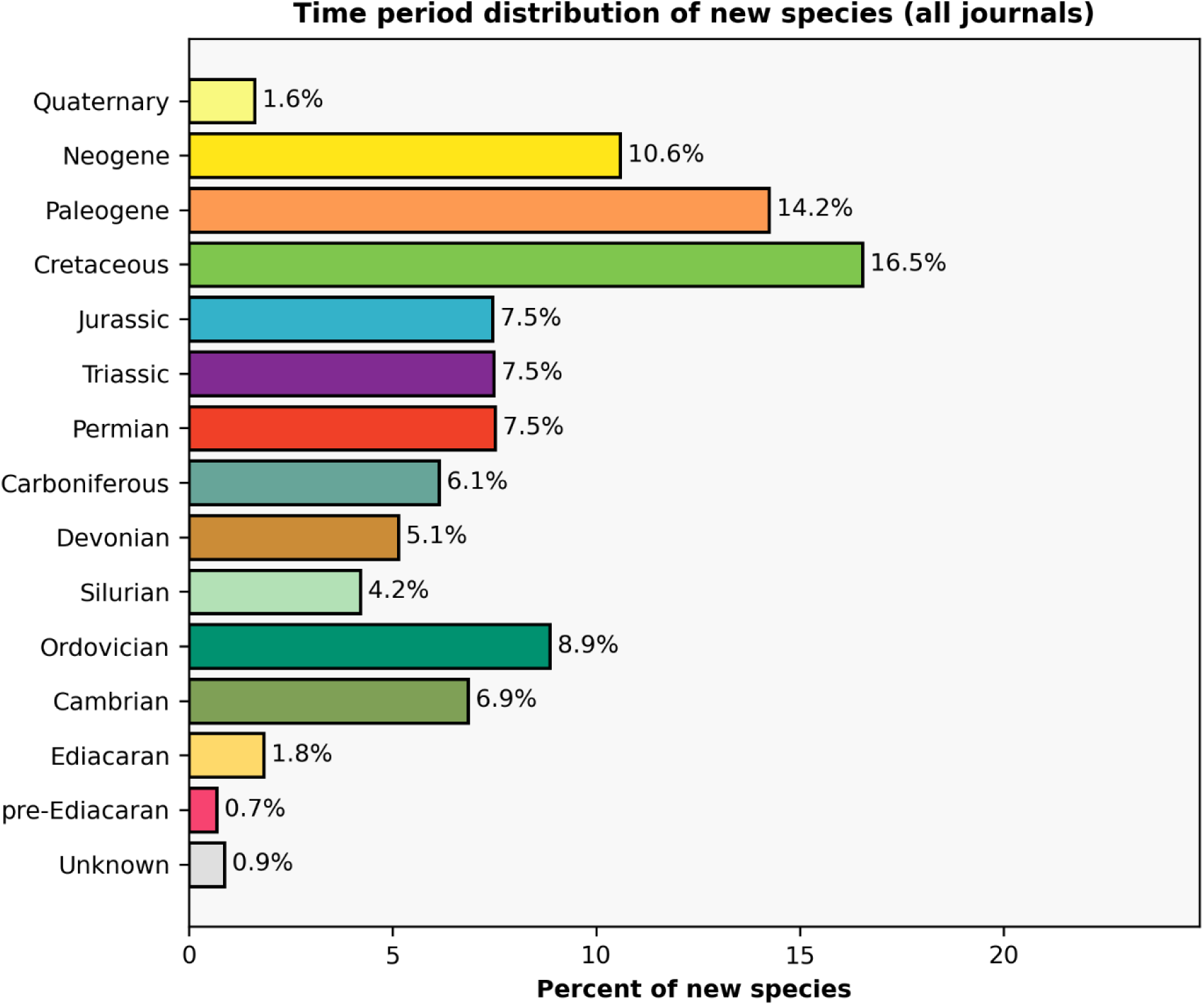
Distribution of new species by temporal bin. Periods are arranged from oldest at the bottom to youngest at the top. ‘Unknown’ entries typically result from a lack of precise placement due to occurrence near a boundary (e.g., near the Oligocene-Miocene boundary) or broader uncertainty around a holotype’s provenance. The color scheme follows the International Commission on Stratigraphy’s Chronostratigraphic Chart (v2026-06).

**Figure 3.**
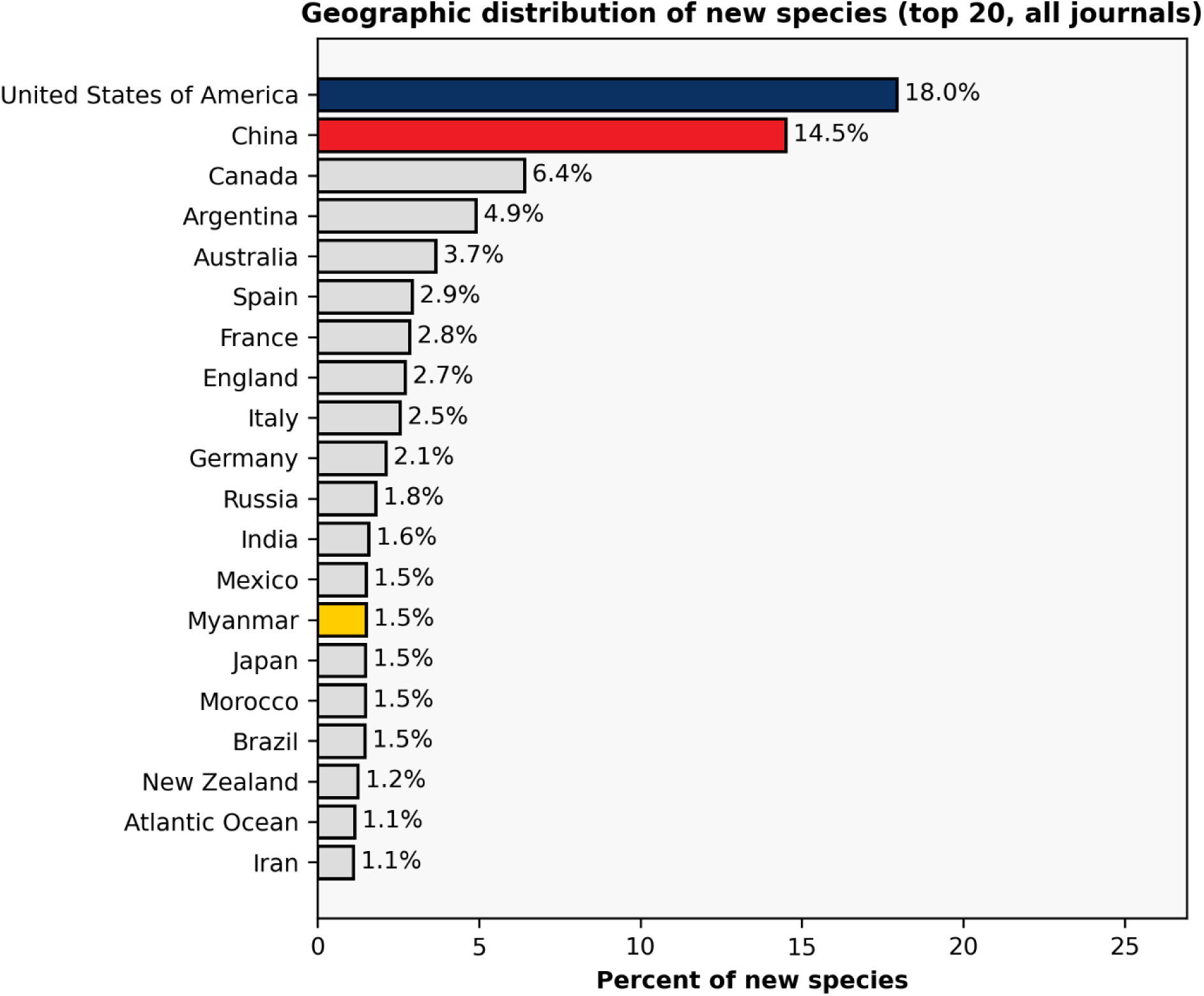
Distribution of new species by geographic provenance of their holotypes (top 20 countries/regions). Only select countries/regions are colored here when they are called out in later parts of the manuscript.

**Table 5.** Summary of the most common temporal and geographic category for each journal sampled here. ‘*Most common age*’ represents the most common age (of known records) of holotypes, ‘most common geography’ represents the most common geographic entity (of known records), and ‘*age+geography*’ represents the most common two-variable combination for each journal; this does not always equal the most common age and the most common geography (e.g., *Gondwana Research*).

| **Journal** | **Most common age** | **Most common geography** | **Most common age+geography** |
| --- | --- | --- | --- |
| *Nature* | Cretaceous | China | Jurassic of China |
| *Science* | Cretaceous | China | Jurassic of China |
| *Current Biology* | Cretaceous | China | Cretaceous of Myanmar |
| *Gondwana Research* | Cretaceous | Australia | Cambrian of Australia |
| *Nature Communications* | Cretaceous | Myanmar | Cretaceous of Myanmar |
| *Nature Ecology & Evolution* | Carboniferous | China/Scotland/United States of America (tie) | Carboniferous of Scotland |
| *Proceedings of the National Academy of Sciences* | Cretaceous | China | Cretaceous of the United States of America |
| *Science Advances* | Cambrian/Cretaceous | China | Cretaceous of Myanmar |
| *Journal of Vertebrate Paleontology* | Neogene | United States of America | Cretaceous of the U.S.A. |
| *Journal of Paleontology* | Ordovician | United States of America | Carboniferous of the U.S.A. |
| *Palaeontology* | Cretaceous | China | Jurassic of China |
| *Papers in Palaeontology* | Triassic | China | Cretaceous of Argentina |
| *Journal of Systematic Palaeontology* | Paleogene | China | Jurassic of China |

### Journal groups

The first sub-section provides summaries of the three groups of journals that were analyzed here. Most of the visualizations combine all journals in a given group together, but journal-specific remarks and some journal-specific visualizations are also presented here; other journal-specific visualizations are included as supplemental figures.

#### *High-JIF journals:* Nature *and* Science

In both journals, a majority of new species belong to one of three categories (two clades): primates, non-primate mammaliamorphs, and non-avian avemetatarsalians (Fig. 4). Non-primate mammaliamorphs are the most common in *Nature* and tied for the most common with primates in *Science* (Fig. 5; Table 4). The remaining distributions differ in a few respects (e.g., *Science* has a distinctly higher proportion of insects but is completely devoid of new species in several categories, such as molluscs and pseudosuchians; Fig. 5). Correspondent with the prevalence of these groups is a temporal skew towards new species from the early Cenozoic (Paleogene) and middle-late Mesozoic (Jurassic-Cretaceous), which collectively represent more than 50% of the holotypes; the Ediacaran and much of the Paleozoic (excepting the Cambrian) are poorly represented (Fig. 6; Table 5; Supp. Fig. 1). For *Nature*, the most common age-category combination is a Cretaceous non-avian avemetatarsalian; for *Science*, it is a Paleogene primate. Finally, both journals are dominated by new species discovered in China (>40% of new species for each journal), which vastly outranks any other country (none exceeds 6.5%; Fig. 7; Supp. Fig. 2).

*Medium-JIF journals:* Current Biology, Gondwana Research, Nature Communications, Nature Ecology & Evolution, PNAS*, and* Science Advances

**Figure 4.**
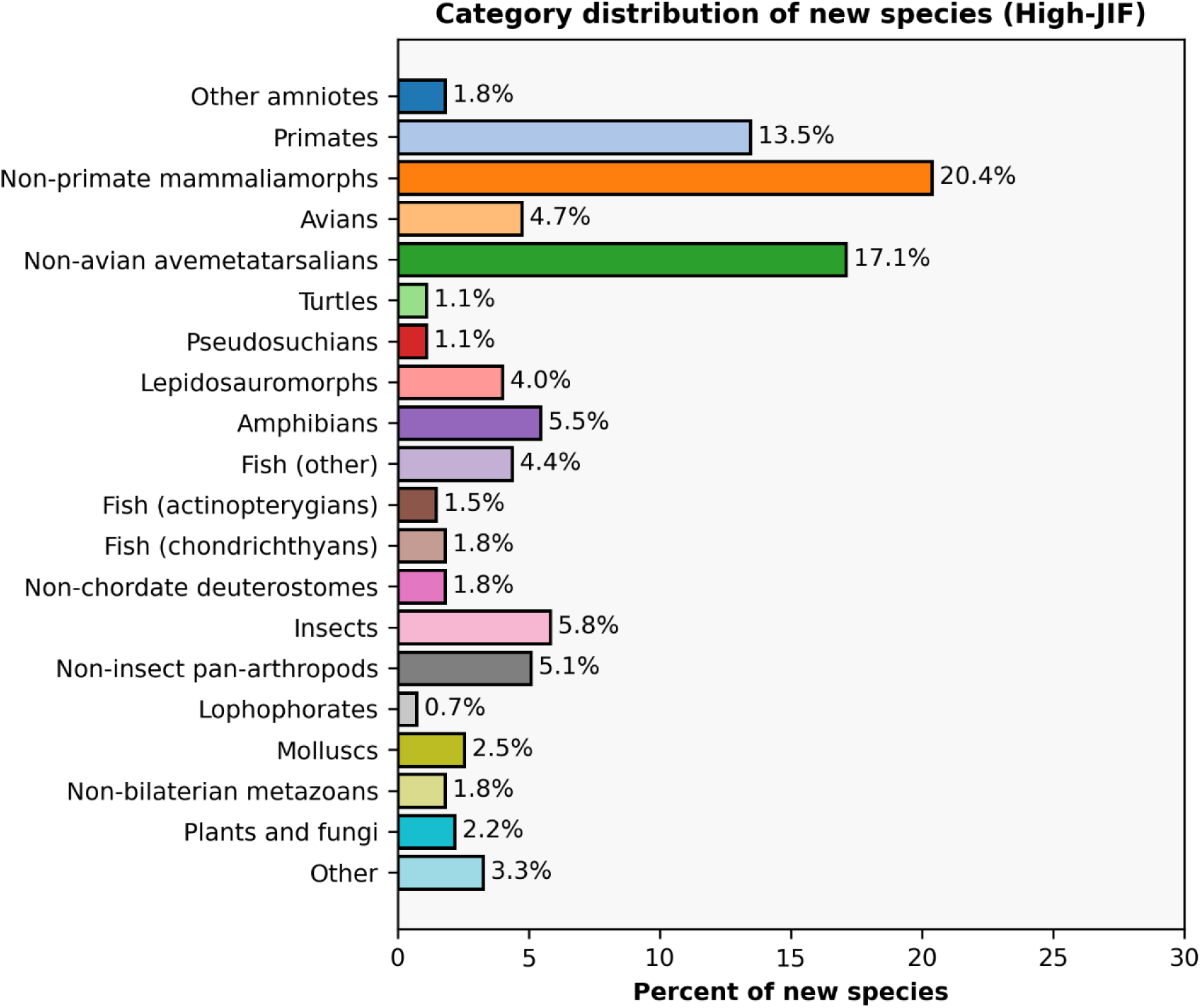
Distribution of taxonomic categories for new species published in *Nature* and *Science*, combined. Categories are roughly arranged in order of phylogenetic divergence, with invertebrates at the bottom, fish and non-amniotes in the middle, and amniotes at the top.

**Figure 5.**
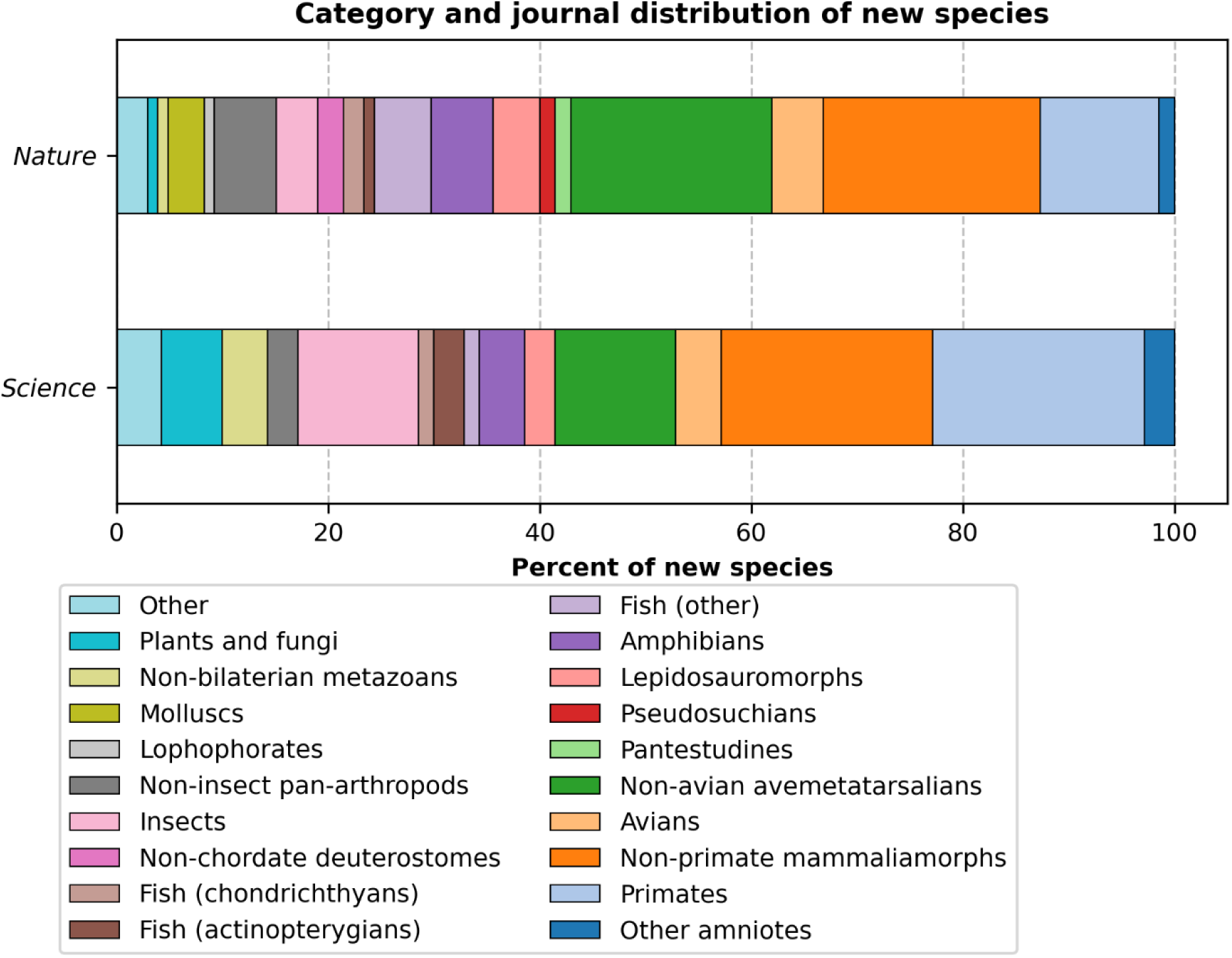
Distribution of taxonomic categories for new species published in *Nature* and *Science*, separated. Categories are roughly arranged in order of phylogenetic divergence, with invertebrates on the left, fish and non-amniotes in the middle, and amniotes on the right.

**Figure 6.**
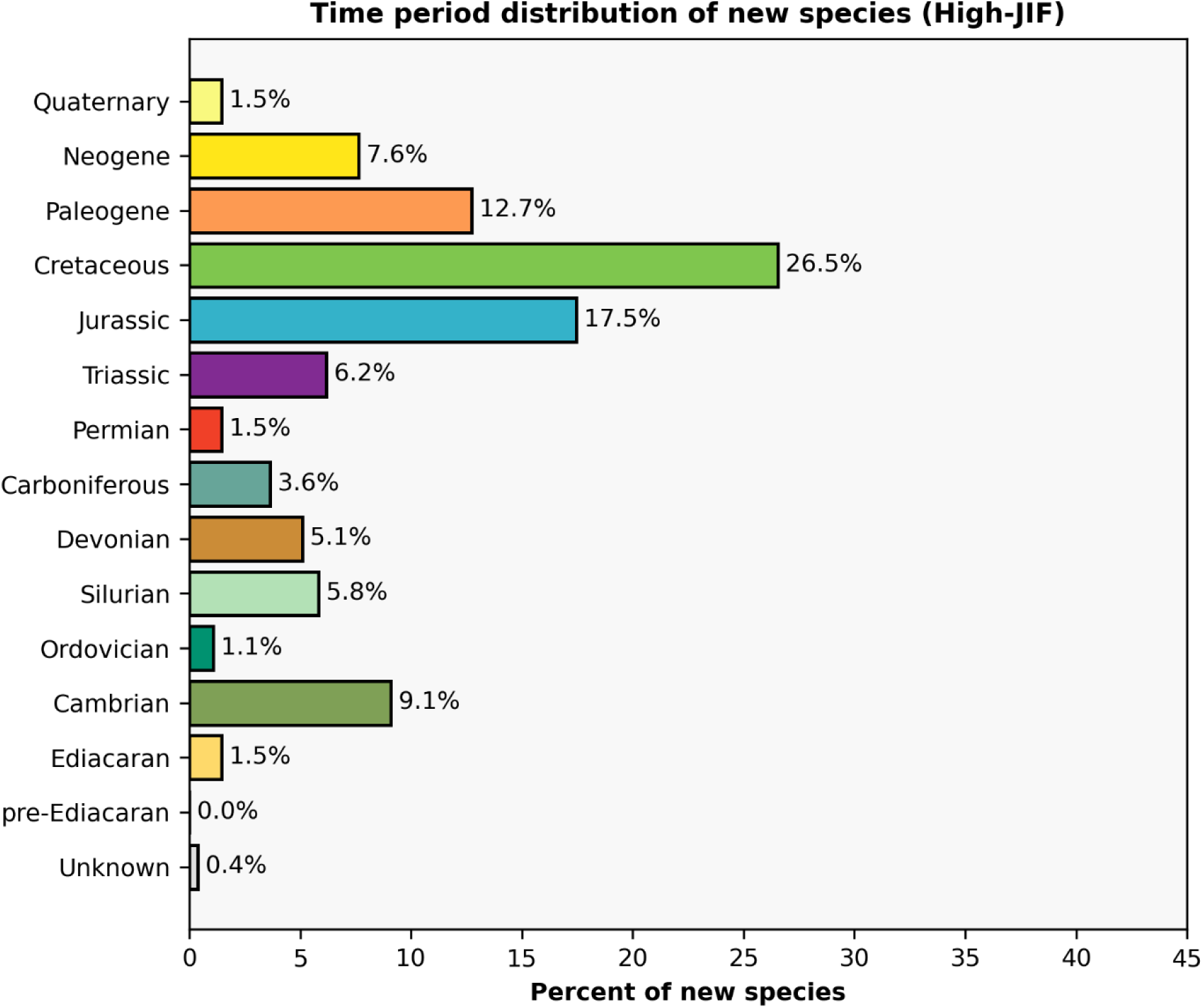
Distribution of ages (where known) of holotypes published in *Nature* and *Science*, combined. Periods are arranged from oldest at the bottom to youngest at the top. ‘Unknown’ entries typically result from a lack of precise placement due to occurrence near a boundary (e.g., near the Oligocene-Miocene boundary) or broader uncertainty around a holotype’s provenance. The color scheme follows the International Commission on Stratigraphy’s Chronostratigraphic Chart (v2026-06).

**Figure 7.**
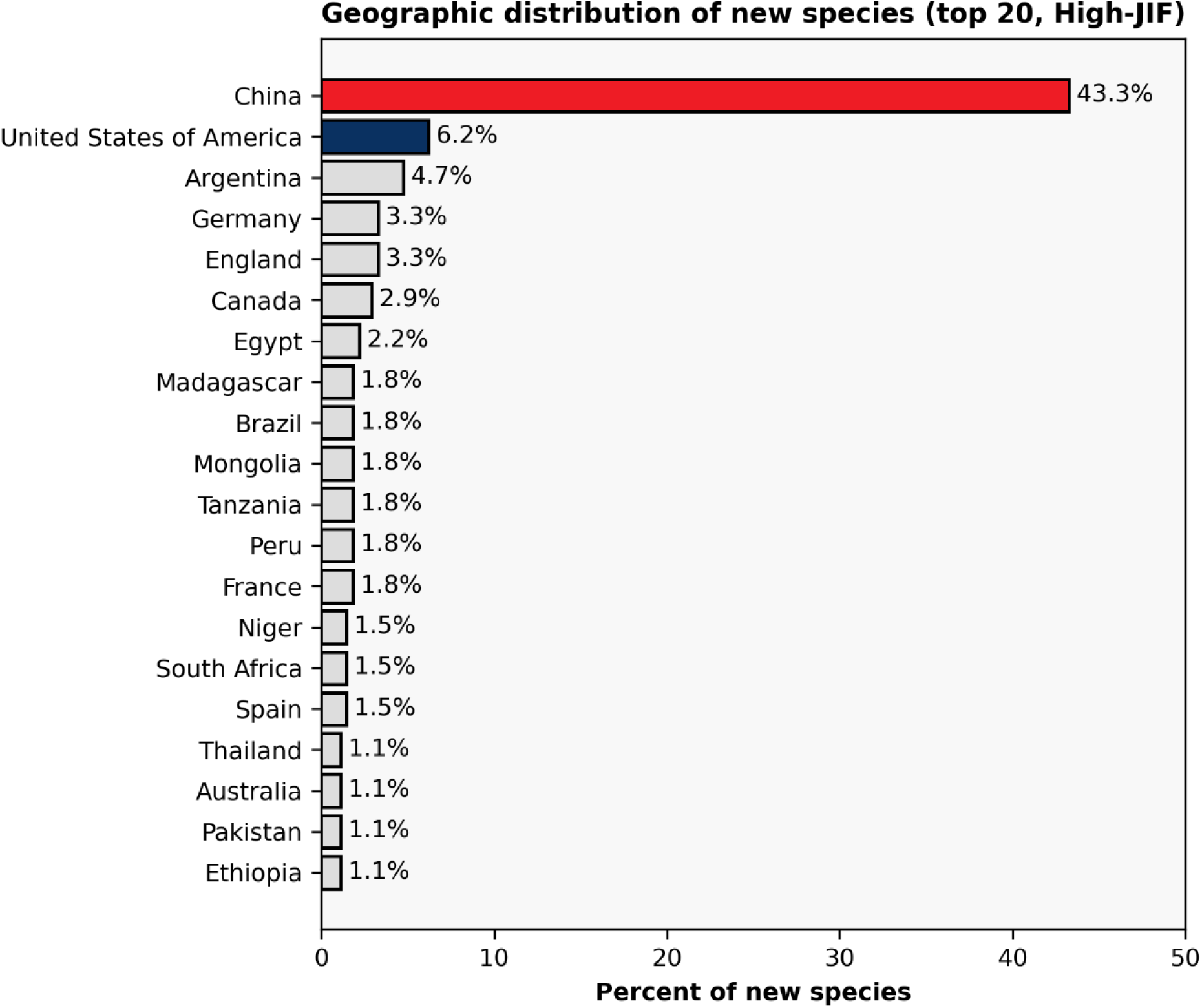
Distribution of the top 20 countries or regions of provenance (where known) of holotypes published in *Nature* and *Science*, combined. Only select countries are colored here when they are called out in later parts of the manuscript.

The medium-JIF journals exhibit some distinct patterns from the high-JIF journals when compared in aggregate (Fig. 8). Although avemetatarsalians (16%) and mammaliamorphs (11.3%) remain relatively well-represented, they are proportionately less represented than in the high-JIF journals (21.8% and 33.9%, respectively; Fig. 4), with pan-arthropods (both inclusive and exclusive of insects) being proportionately much more represented than in the high-JIF journals (10.9% in high-JIF, 35.1% in medium-JIF). Insects in particular are much more represented in the medium-JIF group (23.8% vs. 5.8% in high-JIF group). Other categories with noticeable but less prominent differences include non-bilaterian metazoans (1.8% in high-JIF, 4.8% in medium-JIF) and plants and fungi (2.2% in high-JIF, 4.5% in medium-JIF).

Within the medium-JIF group, there are some prominent differences that reveal the source of the broader inter-group differences noted above (Fig. 9). As a note, *Science Advances* is generally omitted from comparisons here due to its low sample size (Table 3). Firstly, avemetatarsalians and mammaliamorphs are proportionately overrepresented in four of the six journals (as compared to their proportions across the entire dataset; Fig. 2), but in different ways. For example, *Nature Ecology & Evolution* has the highest proportion of avemetatarsalians (35.7%) but a proportion of mammaliamorphs that is more in line with the full sample (10.7% vs. 11.4% across the full dataset). *Current Biology*, *Nature Communications*, and *PNAS* all have avemetatarsalians proportions between ∼18% to 20%. *Gondwana Research* differs from all four in having just 4.9% of its new species classified as avemetatarsalians (none of which are avians).

**Figure 8.**
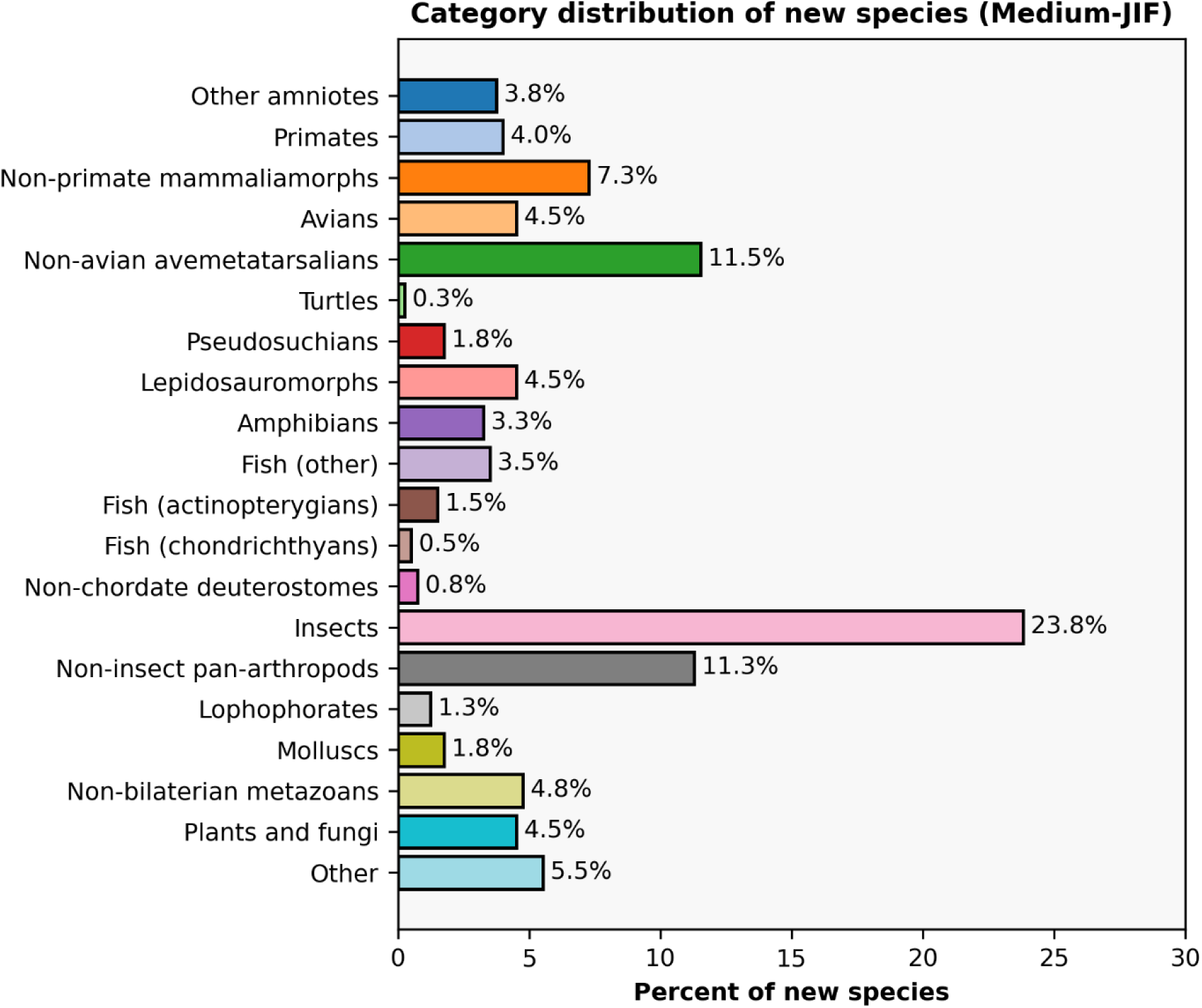
Distribution of taxonomic categories for new species published in Current Biology, Gondwana Research, Nature Communications, Nature Ecology & Evolution, Proceedings of the *National Academy of Sciences*, and *Science Advances*, combined. Categories are roughly arranged in order of phylogenetic divergence, with invertebrates at the bottom, fish and non-amniotes in the middle, and amniotes at the top.

**Figure 9.**
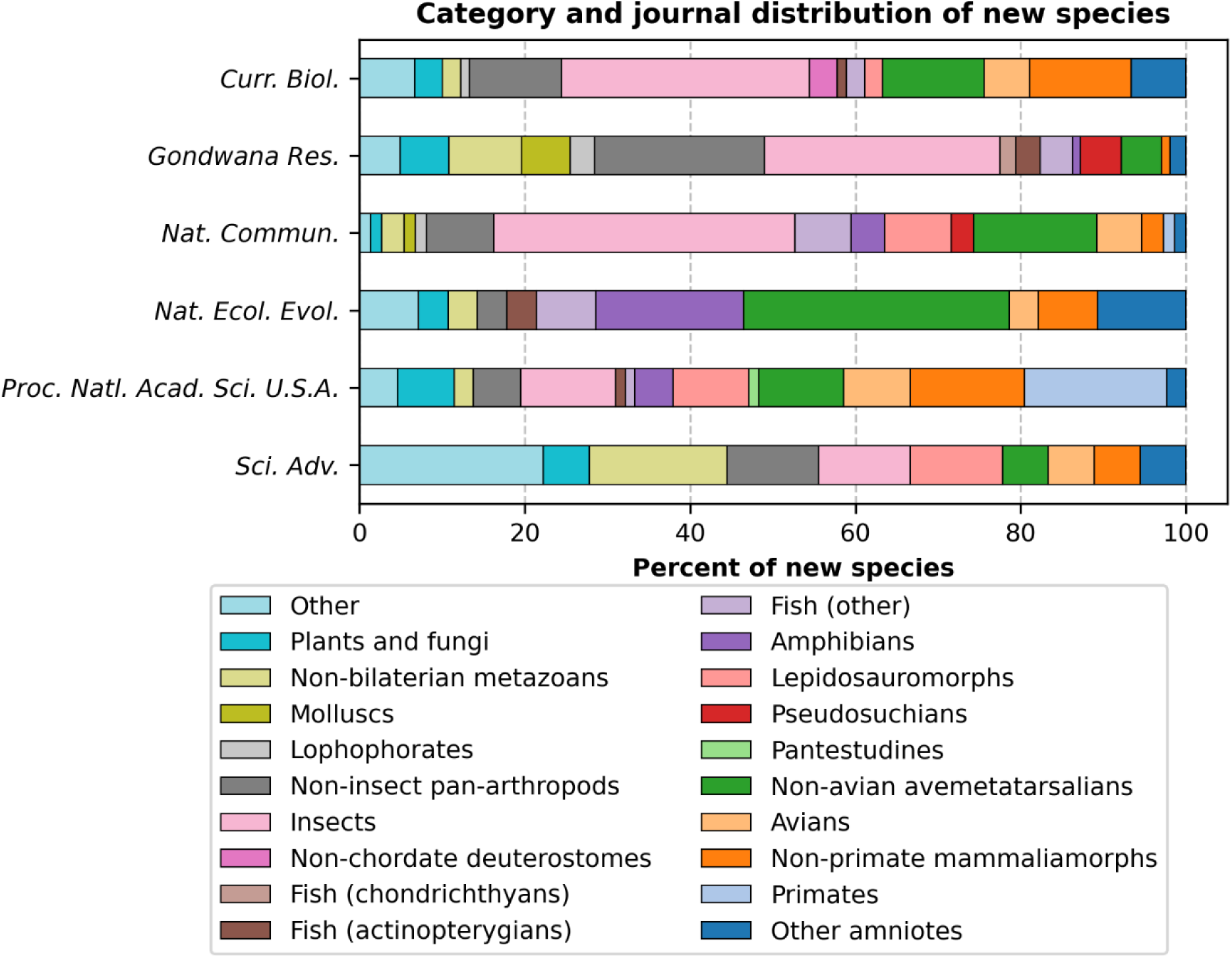
Distribution of taxonomic categories for new species published in Current Biology, Gondwana Research, Nature Communications, Nature Ecology & Evolution, Proceedings of the *National Academy of Sciences*, and *Science Advances*, separated. Categories are roughly arranged in order of phylogenetic divergence, with invertebrates on the left, fish and non-amniotes in the middle, and amniotes on the right.

It also differs markedly in having just 1% of its new species classified as mammaliamorphs (none of which are primates). A general paucity of primates (<1.5% of new species) characterizes all medium-JIF journals except for *PNAS*, in which primates are the most common group of new species, comprising 17.2%. *PNAS* also has a relatively high proportion of non-primate mammaliamorphs (13.8%) that is otherwise only shared with *Current Biology* (12.2%); the other four journals all have less than 7.5% of new species classified as non-primate mammaliamorphs.

The overall abundance of new insects that characterizes the group (Fig. 8) is largely the result of patterns in three journals (*Current Biology*, *Gondwana Research*, *Nature Communications*), each of which have over 28% of their new species classified under this well-known group (Fig. 9). For all three journals, this skew stems from the prevalence of new species published from Cretaceous Myanmar amber specifically (Cretaceous insects and Cretaceous insects from Myanmar are the most common combinations for all three; Figs. 9–10; Table 4; Supp. Fig. 3). Across all clades, Myanmar ranks first in holotype count for *Nature Communications*, second for *Gondwana Research* and *Science Advances*, and ties for second with the United States for *Current Biology*, resulting in the country ranking second for holotype count across the medium-JIF group writ large (Fig. 11; Supp. Fig. 4). Myanmar is tied for 13^th^ across the full dataset (Fig. 3; Supp. Fig. 4) but is not in the top 20 for *Nature* and *Science* combined (it is tied for 8^th^ for *Science*; Fig. 7; Supp. Fig. 2).

**Figure 10.**
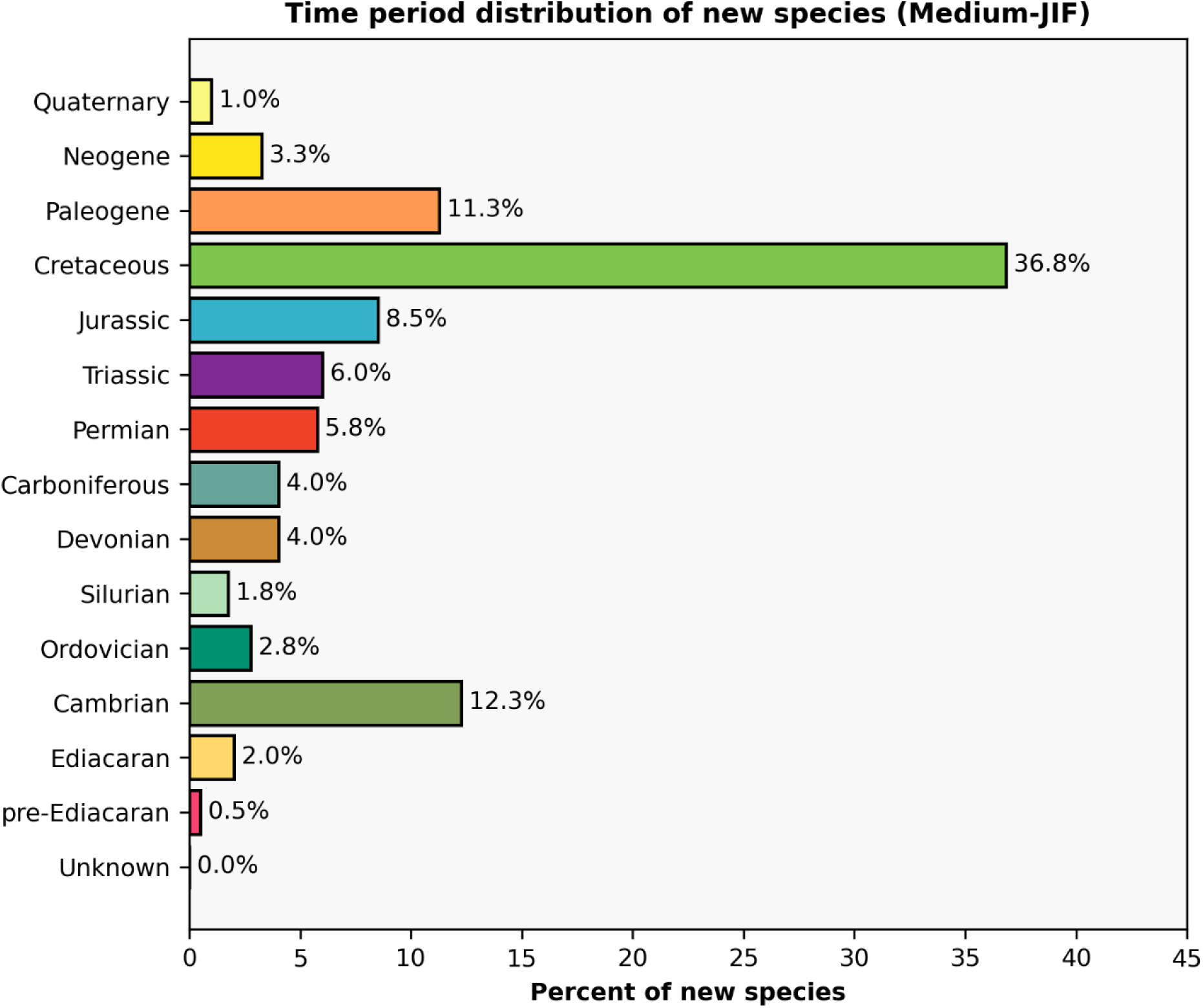
Distribution of ages (where known) of holotypes published in Current Biology, Gondwana Research, Nature Communications, Nature Ecology & Evolution, Proceedings of the National *Academy of Sciences*, and *Science Advances*. Periods are arranged from oldest at the bottom to youngest at the top. ‘Unknown’ entries typically result from a lack of precise placement due to occurrence near a boundary (e.g., near the Oligocene-Miocene boundary) or broader uncertainty around a holotype’s provenance. The color scheme follows the International Commission on Stratigraphy’s Chronostratigraphic Chart (v2026-06).

**Figure 11.**
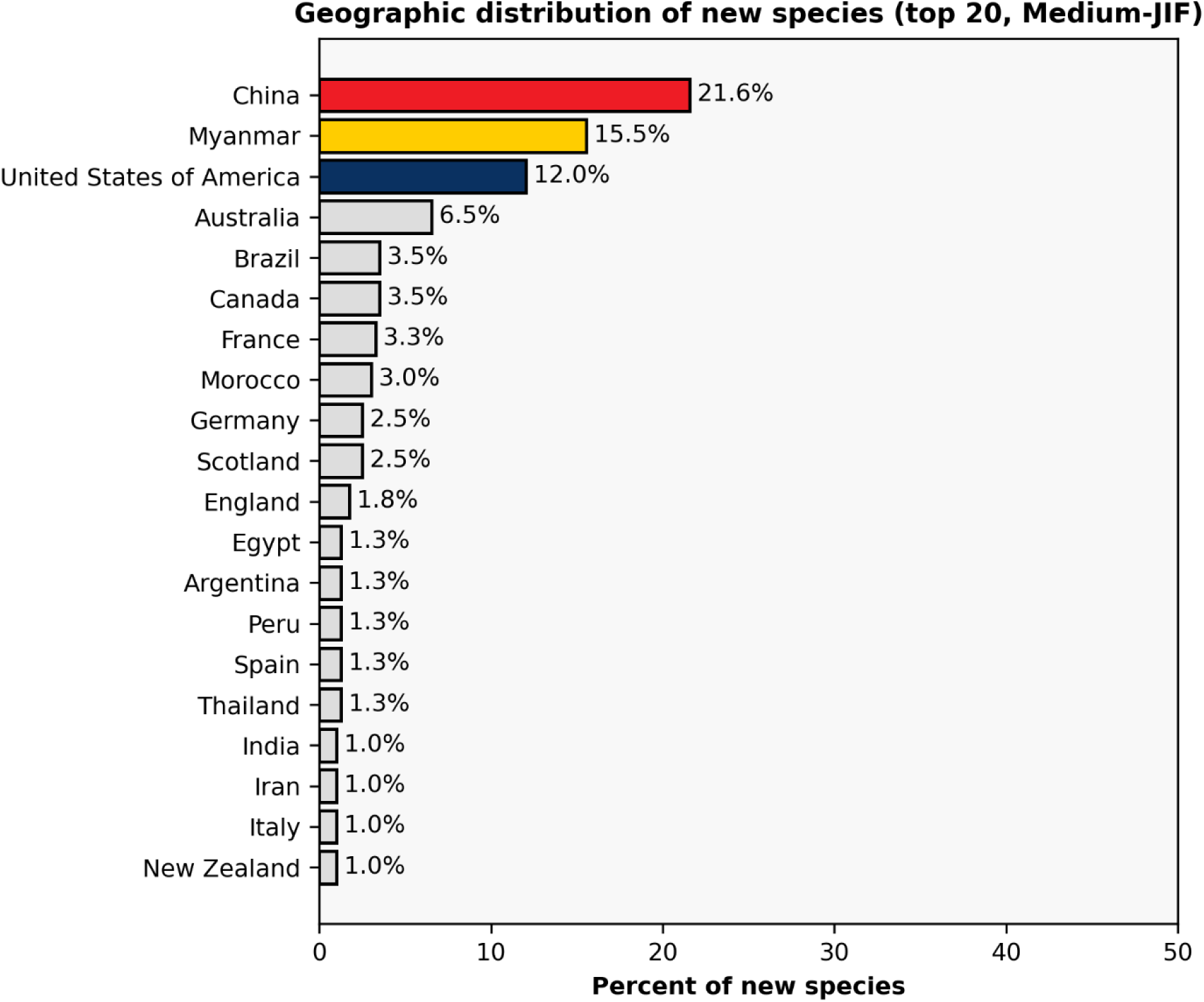
Distribution of the top 20 countries or regions of provenance (where known) of holotypes published in Current Biology, Gondwana Research, Nature Communications, Nature Ecology & Evolution, Proceedings of the National Academy of Sciences, and Science Advances. Only select countries/regions are colored here when they are called out in later parts of the manuscript.

The productivity of these ethically contentious deposits (e.g., Haug et al., 2020; Peretti, 2021; Shi et al., 2021; Zin-Maung-Maung-Thein & Khin Zaw, 2021; Dunne et al., 2022) includes abundant non-insect arthropods (Ross, 2024), hence the moderately high proportions of non-insect pan-arthropods in these three journals as well. For *Current Biology*, *Gondwana Research*, and *Nature Communications*, all pan-arthropods comprise over 40% of all new species, and for *Gondwana Research*, nearly half of all new species are pan-arthropods (Fig. 9). Given the more mixed charisma profile of insects and arthropods, at least some of their appeal for more selective journals probably lies in the exceptional preservation of amber specimens, pointing to another possible factor in editorial decision-making.

Other well-represented clades include lepidosauromorphs, which comprise at least 9% of the new species published in *PNAS* and *Science Advances* (Fig. 9) and non-bilaterian metazoans, which comprise at least 8.5% of new species published in *Gondwana Research* and *Science Advances* (again noting the particularly small sample size of the latter). All six journals have multiple categories without representation.

The temporal and geographic trends (Figs. 10–11) can also be tied to the prevalence of new Myanmar pan-arthropods from the Cretaceous-aged amber deposits in this group. The relative paucity of mammaliamorphs compared to the high-JIF group accounts for lower Cenozoic representation in the medium-JIF group; similar relative paucity of avemetatarsalians accounts for distinctly lower Jurassic representation in the medium-JIF group.

#### Control journals

The control journals predictably show the most distinct patterns in aggregate from the other two groups (Fig. 12). The proportions of mammaliamorphs (inclusive and exclusive of primates) and avemetatarsalians (inclusive and exclusive of avians) are all substantially less than in the high-JIF group (Fig. 4). Other categories with notably less representation in the control group include lepidosauromorphs and amphibians. Somewhat similar to the medium-JIF group, this underrepresentation of vertebrate categories relative to the high-JIF group is compensated for with relative overrepresentation of non-vertebrates. Non-insect pan-arthropods are much more common in the control group than in the high-JIF group (15.7% vs. 5.1%), although interestingly, the proportions of insects are not particularly different (6.6% vs. 5.8%). Lophophorates (9.1% vs. 0.7%), molluscs (14.8% vs. 2.5%), and non-chordate deuterostomes (6.1% vs. 1.8%) also show marked differences. The percent of new species placed in the catch-all ‘Other’ is also much higher in the control group (9.2% vs. 3.3%). Compared to the medium-JIF group (Fig. 8), the charismatic vertebrate categories are still underrepresented in the control group, with the exception of non-primate mammaliamorphs, as are lepidosauromorphs, amphibians, and insects. This is primarily compensated for with higher representation of molluscs, lophophorates, and non-chordate deuterostomes in the control group.

**Figure 12.**
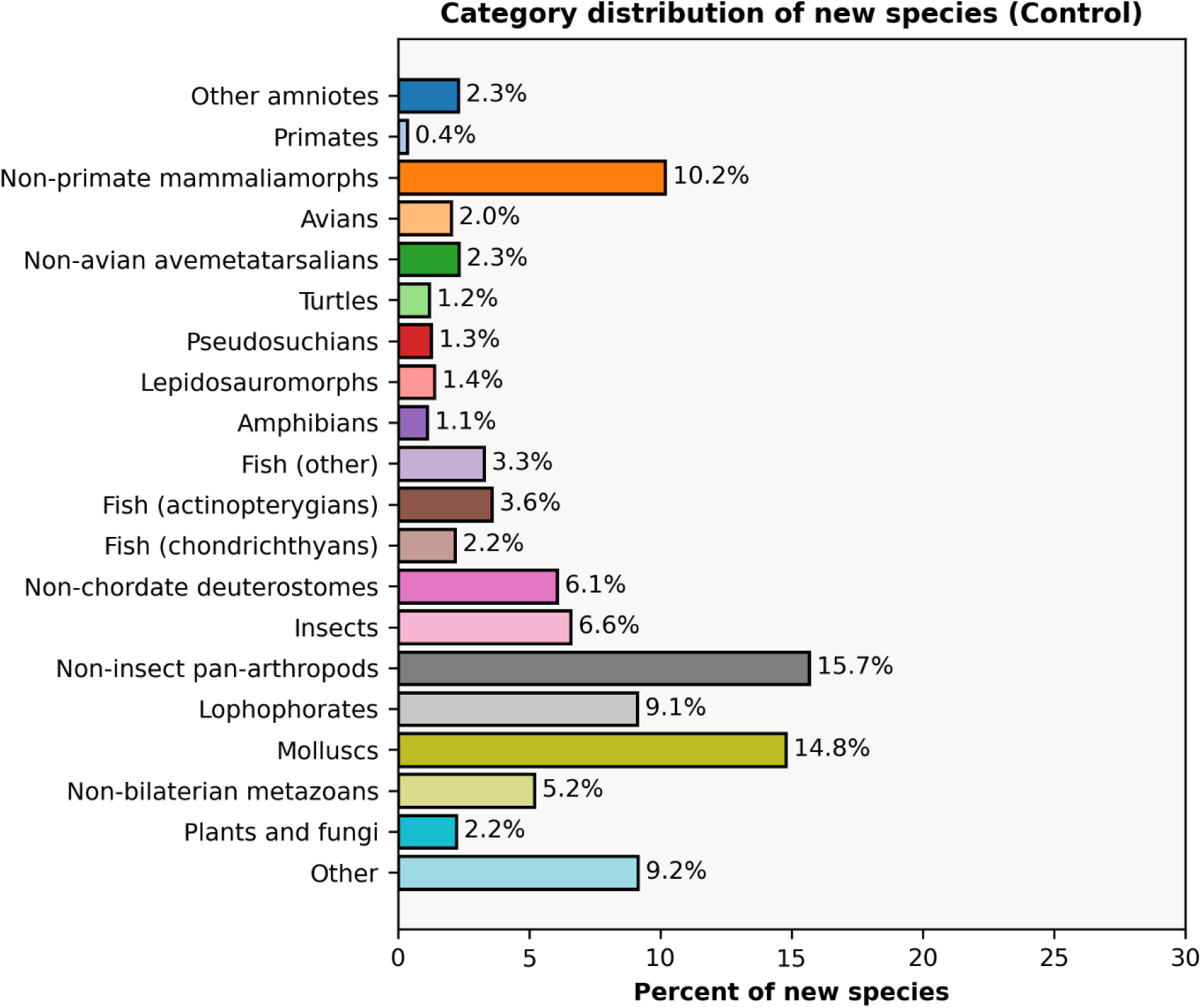
Distribution of taxonomic categories for new species published in the Journal of Paleontology, the Journal of Systematic Palaeontology, the Journal of Vertebrate Paleontology, Palaeontology, and Papers in Palaeontology, combined. Categories are roughly arranged in order of phylogenetic divergence, with invertebrates at the bottom, fish and non-amniotes in the middle, and amniotes at the top.

Within the control group, there are also fairly distinct patterns (and not merely on account of the vertebrate-only focus of the *Journal of Vertebrate Paleontology*). Firstly, the *Journal of Vertebrate Paleontology* is taken as a benchmark for a vertebrate-focused sample since vertebrate naming papers are frequently proportionately overrepresented in high-JIF journals. It has by far the highest proportion of non-primate mammaliamorphs (36.2%), which contrasts with a very low proportion of primates (just 1%), as well as the highest proportion of actinopterygians (12.3%; no other journal has more than 5% of its new species as actinopterygians; Fig. 13). The proportion of avemetatarsalians in this journal is higher than the proportion across the full dataset (Fig. 1) and is slightly higher than that of the medium-JIF journals (Fig. 8), but it is still less than that of the high-JIF journals (Fig. 4). Additionally, the ratio of avians to non-avian avemetatarsalians is more even in this journal (7.3% and 9.4%, respectively) than in the high-JIF group (4.7% and 17.1%, respectively). Most of the vertebrate categories that are scarcely represented in other journals are well-represented here, including chondrichthyans, amphibians, lepidosauromorphs, turtles, and pseudosuchians (Fig. 13).

**Figure 13.**
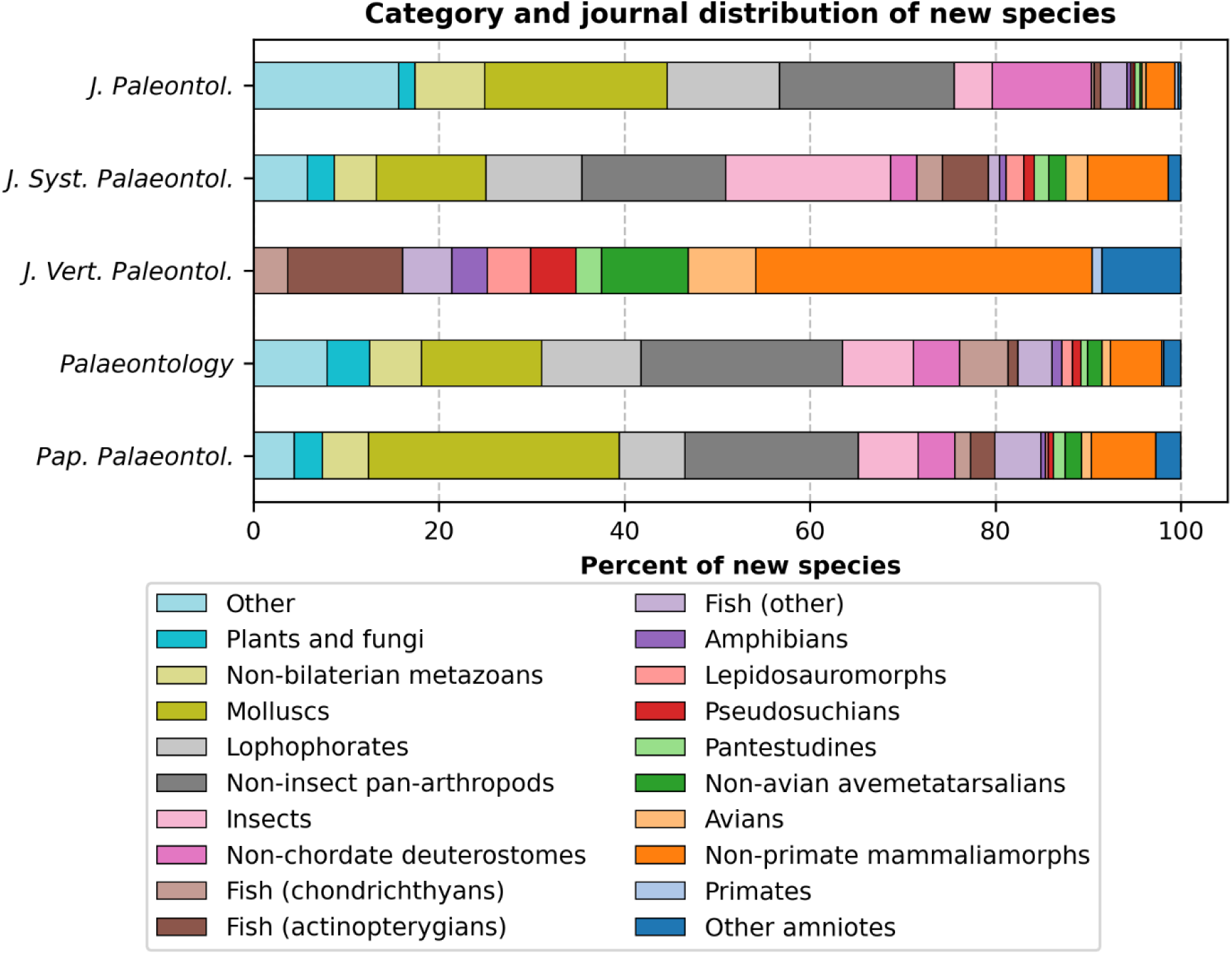
Distribution of taxonomic categories for new species published in the Journal of Paleontology, the Journal of Systematic Palaeontology, the Journal of Vertebrate Paleontology, Palaeontology, and Papers in Palaeontology, separated. Categories are roughly arranged in order of phylogenetic divergence, with invertebrates at the bottom, fish and non-amniotes in the middle, and amniotes at the top. Categories are roughly arranged in order of phylogenetic divergence, with invertebrates on the left, fish and non-amniotes in the middle, and amniotes on the right.

In sharp contrast, the other four control journals are notably underrepresented with respect to most vertebrate clades (Fig. 13). Avemetatarsalians (never above 4.5%) and mammaliamorphs (never above 9%) are lower than the full dataset proportions, and most other vertebrate clades do not comprise more than about 1.5 to 2% of a given journal’s new species. Instead, all four are dominated by invertebrates. The *Journal of Systematic Palaeontology* has a high proportion of insects driven primarily by new species from China, and to a lesser extent, new taxa from Myanmar (note that this journal has prohibited publication of such material since 2020; Barrett & Johanson, 2020; Table 4; Supp. Fig. 6). The journal is also well-represented by non-insect pan-arthropods, lophophorates, and molluscs. *Palaeontology* generally has a similar distribution to the *Journal of Systematic Palaeontology*, but with non-insect pan-arthropods greatly outpacing insects. *Papers in Palaeontology* and the *Journal of Paleontology* have strong representation of non-insect pan-arthropods but also of molluscs, which are about 20% in the latter and over 25% in the former (the most common category in each). The *Journal of Paleontology* also has strong representation by non-chordate deuterostomes, mainly through prolific naming of echinoderms. It also has the highest proportion of ‘Other’ species, which includes less charismatic groups like acritarchs, foraminifera, and radiolarians. Correspondingly, this journal also has the lowest proportion of new vertebrate species of any sampled journal – less than 10%.

Temporally, the control journals show a more even distribution among temporal bins, although the Cretaceous, the Paleogene, and the Neogene still represent over 40% of all holotypes (Fig. 14). The *Journal of Vertebrate Paleontology* has the strongest skew towards these three, with over 66% of all holotypes coming from these bins and a correspondingly low proportion of holotypes from the Paleozoic (Supp. Fig. 5). Conversely, the other journals, with much richer non-vertebrate corpuses, have better representation in the Paleozoic, though only the *Journal of Paleontology* is characterized by a Paleozoic time bin as the most common (Ordovician; Supp. Fig. 5; Table 4).

Spatially, the United States and China are the leading producers of novel species across the control group (Fig. 15). Comparison between journals reveals some patterns that suggest a correlation with the country a sponsoring entity is based in; the two U.S.-based journals (*Journal of Paleontology*, *Journal of Vertebrate Paleontology*) have published more new species from the United States than from any other country, with China being second, whereas the other three control journals (U.K.-based sponsors) are led by new species from China, with European countries ahead of the United States (Supp. Fig. 6).

**Figure 14.**
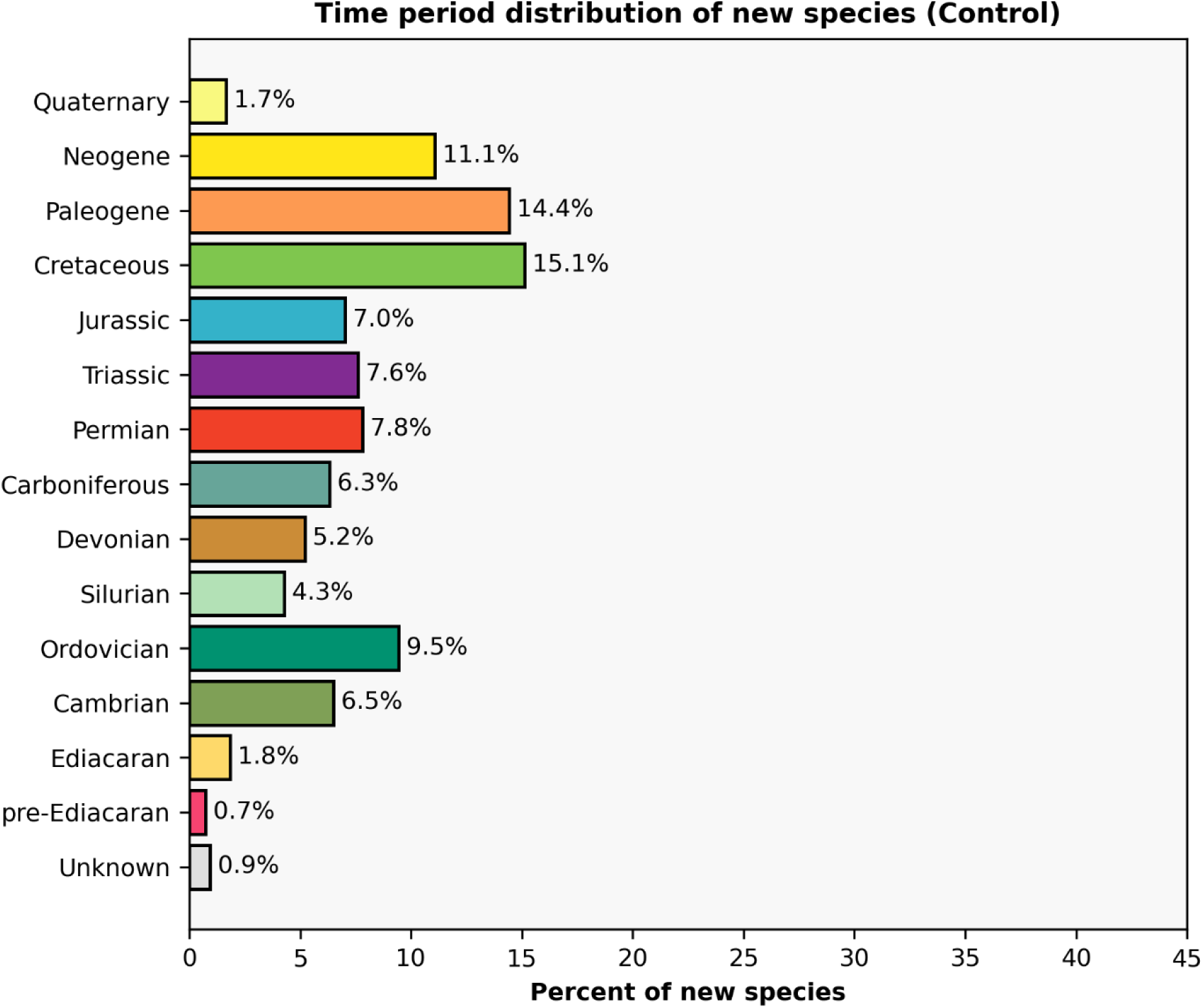
Distribution of ages (where known) of holotypes published in the Journal of Paleontology, the Journal of Systematic Palaeontology, the Journal of Vertebrate Paleontology, Palaeontology, and Papers in Palaeontology. Periods are arranged from oldest at the bottom to youngest at the top. ‘Unknown’ entries typically result from a lack of precise placement due to occurrence near a boundary (e.g., near the Oligocene-Miocene boundary) or broader uncertainty around a holotype’s provenance. The color scheme follows the International Commission on Stratigraphy’s Chronostratigraphic Chart (v2026-06).

**Figure 15.**
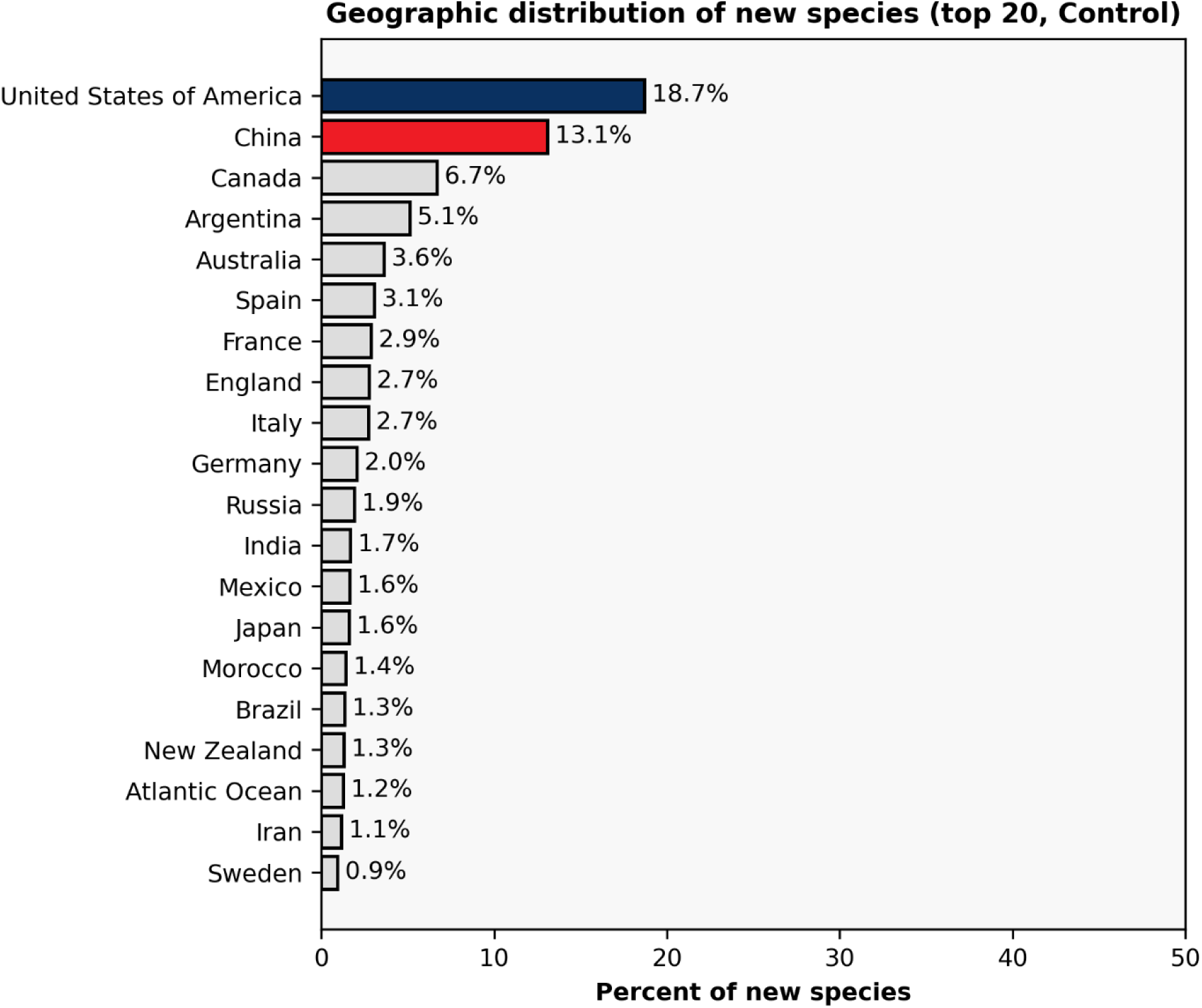
Distribution of the top 20 countries or regions of provenance (where known) of holotypes published in the *Journal of Paleontology*, the *Journal of Systematic Palaeontology*, the *Journal of Vertebrate Paleontology*, *Palaeontology*, and *Papers in Palaeontology*. Only select countries/regions are colored here when they are called out in later parts of the manuscript.

#### Clade-specific patterns

Several of the 20 taxonomic categories are further visualized and briefly discussed in this section, with a focus on contextualizing some of the aforementioned journal-level patterns.

The first case study is avemetatarsalians, which are widely agreed upon as one of the most charismatic clades, primarily through dinosaurs (Fig. 16). Different avemetatarsalian clades exhibit distinct patterns between journals, and readers are likely particularly interested in the breakdown of dinosaurs, which is shown here. Across all journals other than *Gondwana Research* (which has no saurischians in the present sample) and the small-sample-size *Science Advances*, saurischian naming papers are more common than ornithischian naming papers. Furthermore, the high-JIF journals and the medium-JIF journals, especially the three Nature Portfolio titles, *Science*, and *PNAS*, have relatively high portions of non-avian theropods (dark green in Figure 16), which includes many of the most iconic taxa such as tyrannosauroids and spinosaurids. Pterosaurs, a relatively charismatic non-dinosaur clade, are well-represented across most journals. The catch-all ‘Other non-dinosaurs’ category includes taxa that capture the early stages towards dinosaur evolution (e.g., the aphanosaur *Teleocrater*; Nesbitt et al., 2017 [*Nature*]).

**Figure 16.**
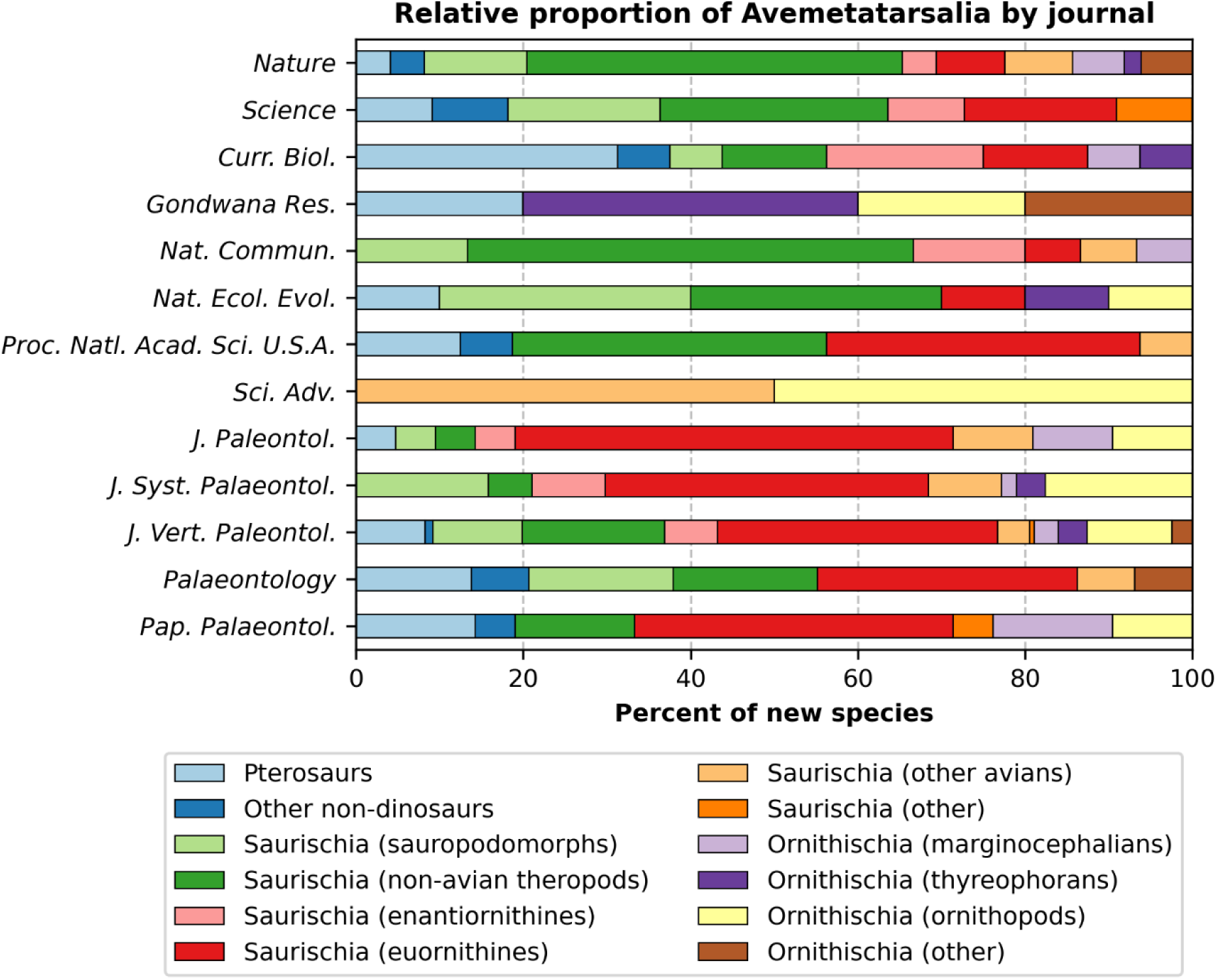
Proportional representation of different avemetatarsalian groups between journals.

The second case study is mammaliamorphs, another charismatic clade. Many taxa on the mammalian stem (non-mammalian mammaliamorphs; darkest blue in Fig. 17) have also been described in high-JIF journals, particularly *Nature*, likely because they contribute insight into the acquisition of mammalian features; eutriconodonts are particularly common in this journal (Fig. 17). Across the high- and medium-JIF journals, primates (dark brown in Fig. 17) or ungulates (mustard in Fig. 17) are the most common group. In the control journals, ungulates are still appreciably common, but a number of other clades like Metatheria, Afrotheria, Rodentia, and the catch-all ‘Other placental mammals’ (e.g., condylarths) are better represented, and primates are relatively rare. Rodents in particular are either the most common or the second most common clade in the control-group journals.

The third case study is pan-arthropods, the most speciose category in this dataset (*n* > 1,800). Pan-arthropods are undoubtedly well-recognized, arguably to the same extent as dinosaurs and mammals, but their charisma is probably best characterized as middling to low among the broader public, with certain groups being generally reviled (e.g., Shipley & Bixler, 2017; Sumner et al., 2018; Mammola et al., 2020; Fukano & Soga, 2021). Especially in high-JIF journals, insects are typically the most common arthropod clade, although they are far less common in the control group where slightly less well-known groups (to the public) like ostracods represent an appreciable minority (Fig. 18). Trilobites, a fairly well-known extinct clade are also much more common in the control group. Other groups common in higher-JIF journals include radiodonts, a fairly well-known group of relatively large arthropods (viz. taxa like *Anomalocaris* Whiteaves, 1892) and arachnids.

**Figure 17.**
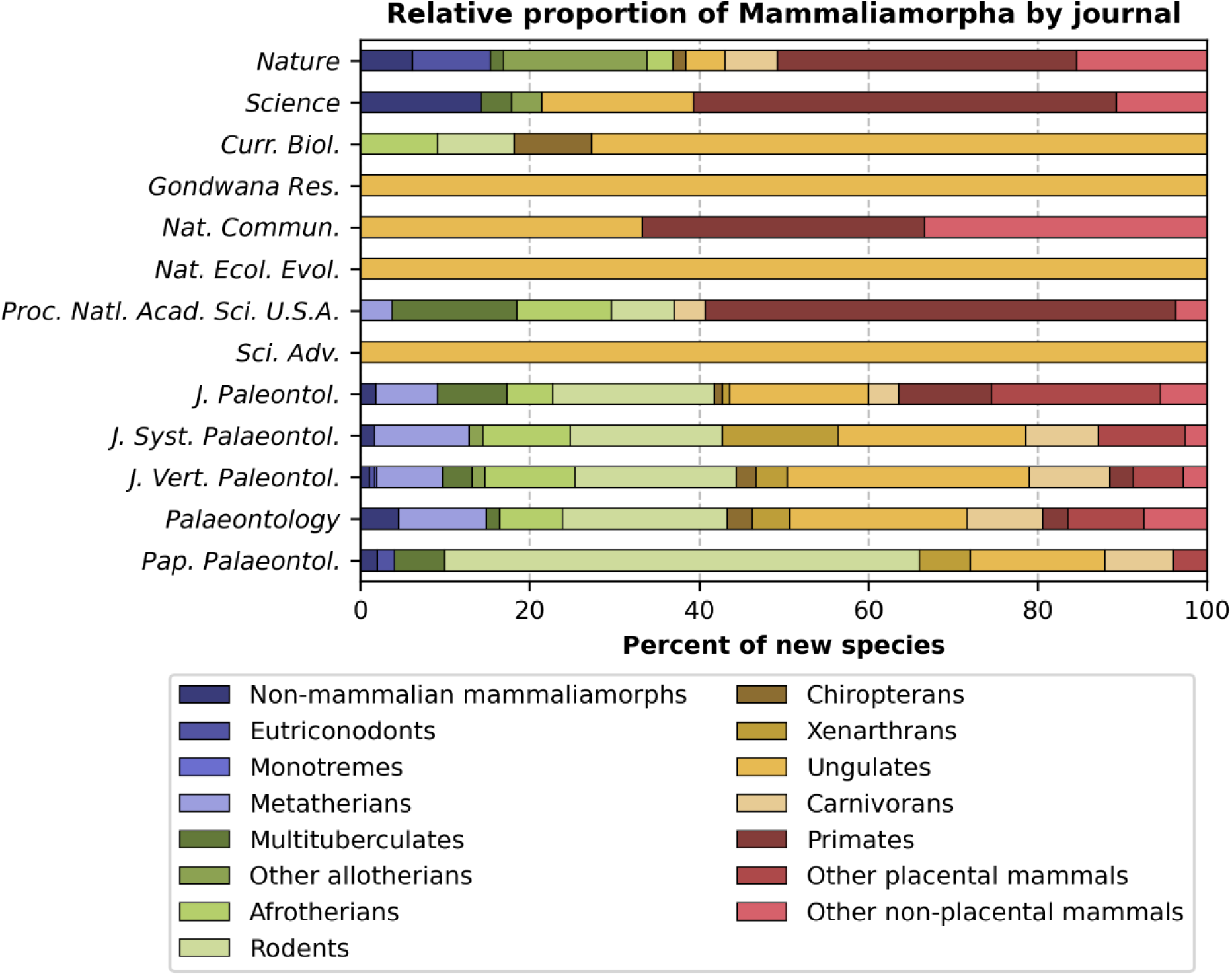
Proportional representation of different mammaliamorph groups between journals.

**Figure 18.**
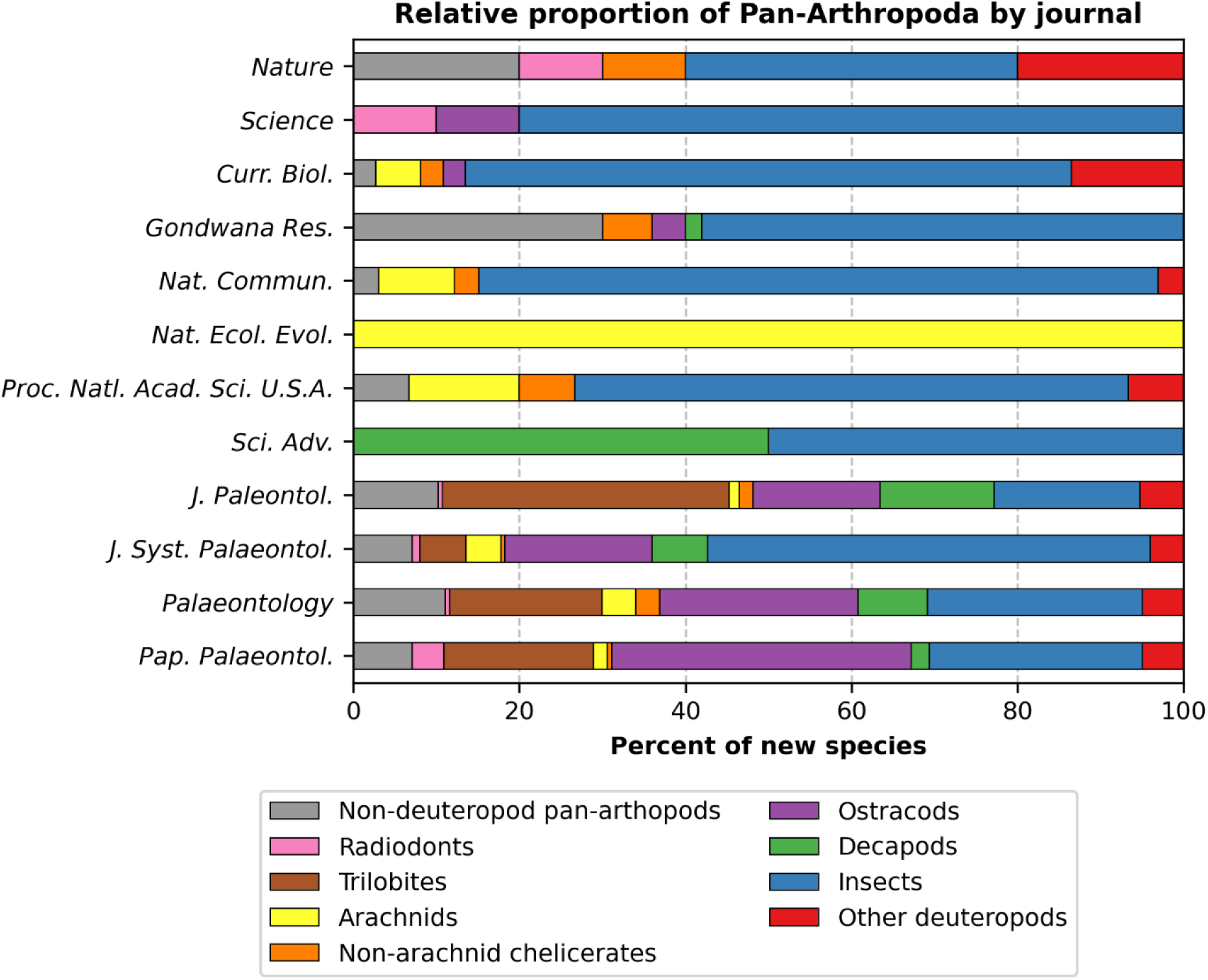
Proportional representation of different pan-arthropod groups between journals. The *Journal of Vertebrate Paleontology* is omitted because it does not publish invertebrate material

Insects are usually the most common pan-arthropod clade in high- and medium-JIF journals (*Current Biology*, *Gondwana Research*, *Nature Communications*), as well as in the *Journal of Systematic Paleontology* (compared to other control journals; Fig. 18). As previously mentioned, this can be largely tied to publication of new species of insects from Myanmar amber (Figs. 8– 11; Table 4). This significant source of new species (globally and for this dataset) warrants some further discussion given the relative lack of charisma of insects.

Across all sampled journals, insects account for 80% of new species from Myanmar, most but not all of which are from Myanmar amber (Fig. 19), although Myanmar is outranked by China in terms of which country was the source for the highest number of new insect species among sampled journals (Fig. 20). New species from the less contentious Baltic amber are also well-represented, albeit in lower-JIF journals (this accounts for “Paleogene insects” being the most common age-category combination for the *Journal of Systematic Palaeontology*; Table 4); the lack of country-level provenance for many holotypes from Baltic amber accounts for the 2.9% categorized as ‘Not defined’ in terms of geographic provenance (Fig. 20).

**Figure 19.**
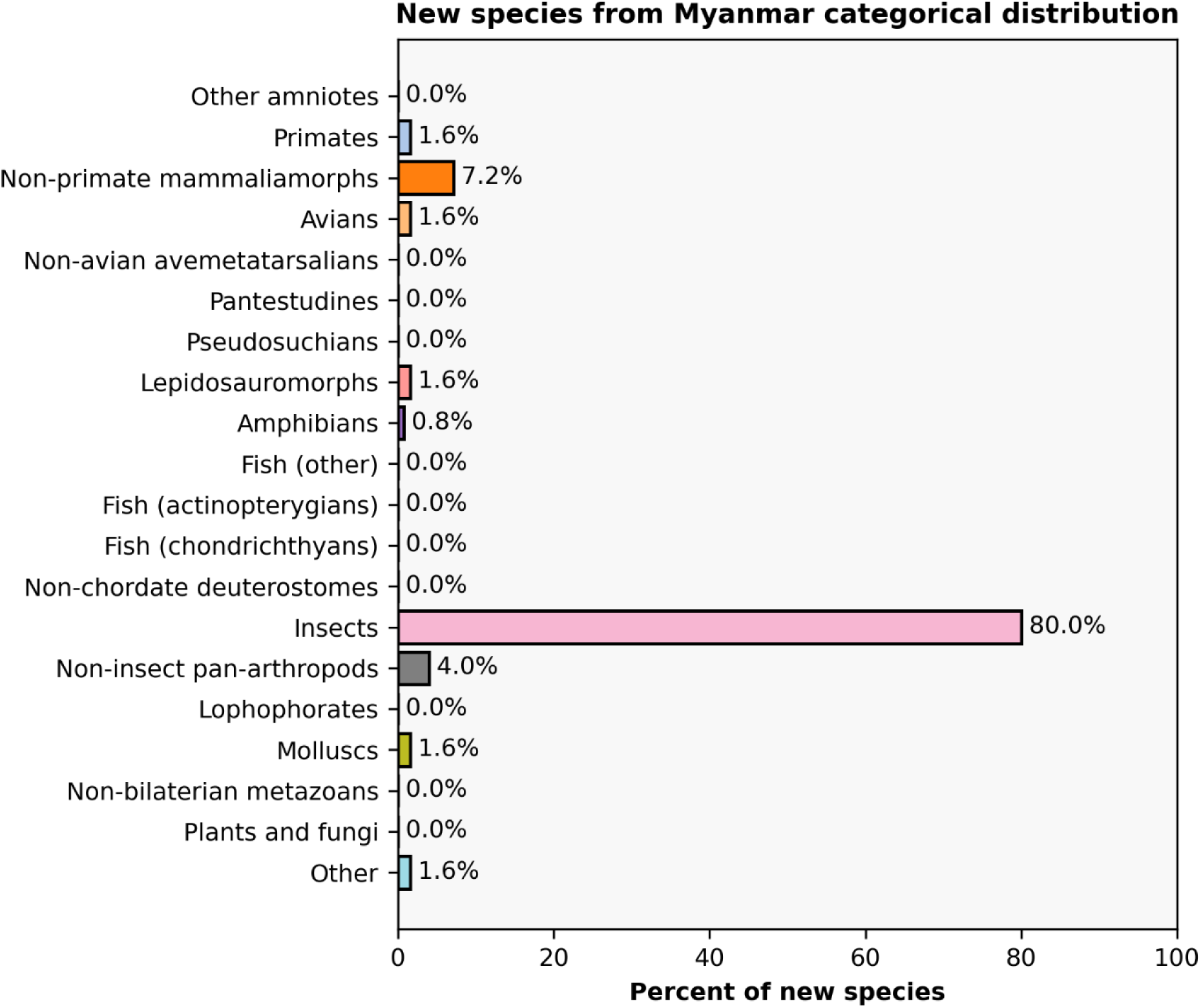
Distribution of taxonomic categories for new species from Myanmar that were published in the journals sampled here. Note that most, but not all, new species come from Myanmar amber.

**Figure 20.**
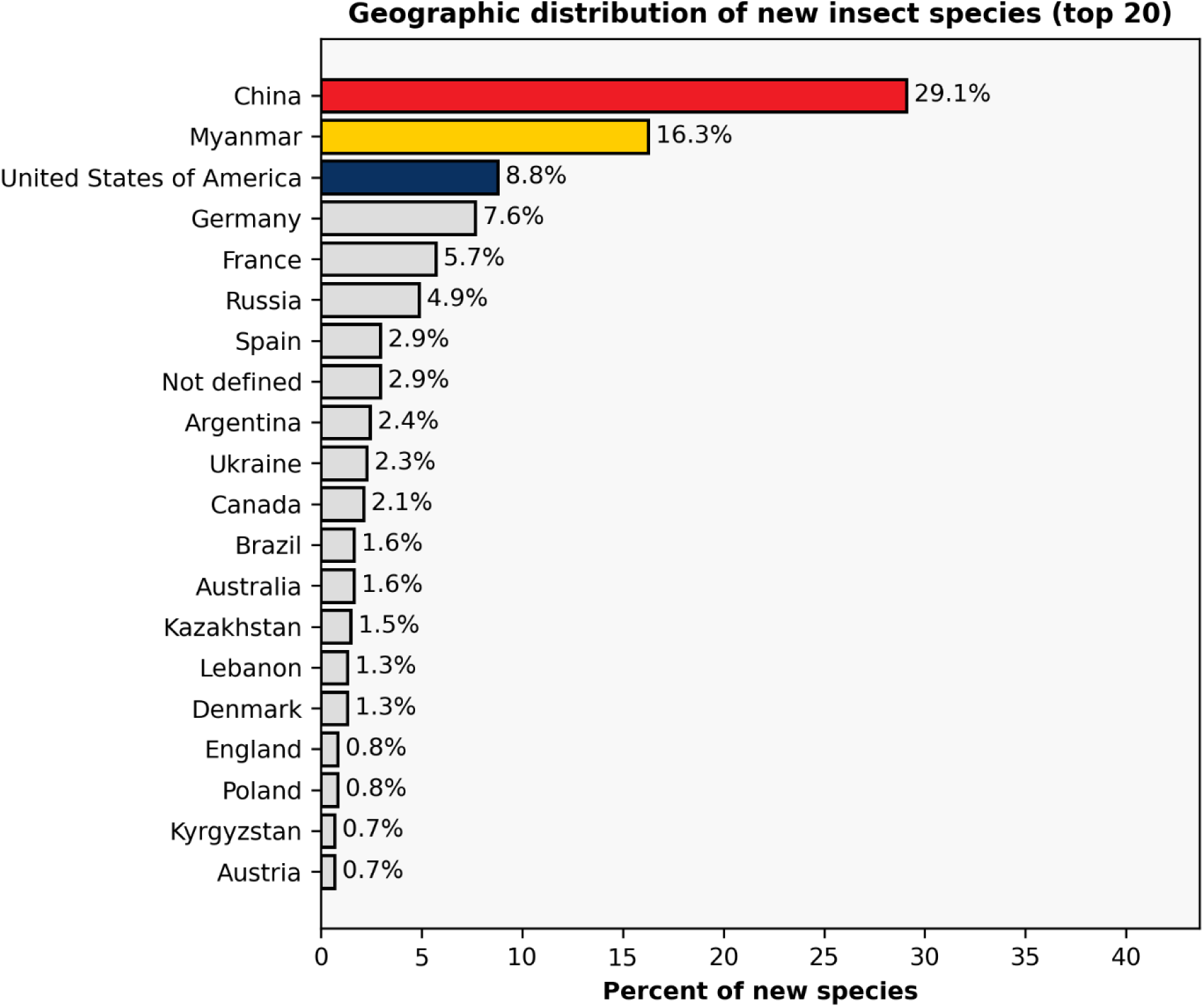
Distribution of the top 20 countries or regions of provenance (where known) of holotypes of insects published in any one of the journals sampled here. Only select countries/regions are colored here when they are called out in later parts of the manuscript.

An important observation given the quarter-century-long sampling of this study is that publishing trends at differing levels have probably fluctuated to some degree over time (e.g., there is no reason to assume the same proportion of new extinct primates that are published in *PNAS* each year is the same). In the above example of new species from Myanmar, there is a clear rise in the late 2010s, peaking in 2018 across the sampled journals, and then declining to levels less than, or around, just 25% of the peak in the most recent years (Fig. 21). Dunne et al. (2022) suggested a possible decline in total publications on Myanmar amber in the early 2020s might be due to increased scrutiny by some groups within the research community. Temporal trends in general are not further investigated here because of the many confounds over the sample interval (e.g., the rise of OA publishing, country-level changes in research funding and support).

Insects also provide a useful entry to brief discussion of some marked skews in visualizations of relative proportions that are introduced by another pattern in naming papers – the relative tendency for non-vertebrate workers to name multiple species in a single paper compared to vertebrate workers (Fig. 22). For example, the large proportion of ‘Other antliophorans’ in *Science* is the result of two factors, a small sample size of new taxa (*n* = 8) and an even smaller number of naming papers (*n = 3*); Ren et al. (2009) described six new species of scorpionflies, which are too rare in other journals to warrant a separate category in the figure. The domination of ‘Other insects’ in *Nature* can also be explained in a similar fashion (also *n* = 8 for species, *n* = 4 for naming papers); Nel et al. (2013) described five new species of holometabolous insects, some of which belong to early diverging clades without extant representation. This is not to say that vertebrate workers never publish multiple new species in a single publication. The combination of small article sample size and a multi-species paper explains why Carboniferous tetrapods from Scotland (Clack et al., 2016; five new species of ‘amphibians’ as defined here) are the most common three-variable combination in *Nature Ecology & Evolution*. However, the dataset analyzed here reflects a much higher tendency for multi-species papers to describe non-vertebrates across all three journal groups (Figs. 23–24). Journals with larger samples show less skewedness towards any single group of insects, with hymenopterans, coleopterans, and dipterans being among the best represented clades (Fig. 22).

**Figure 21.**
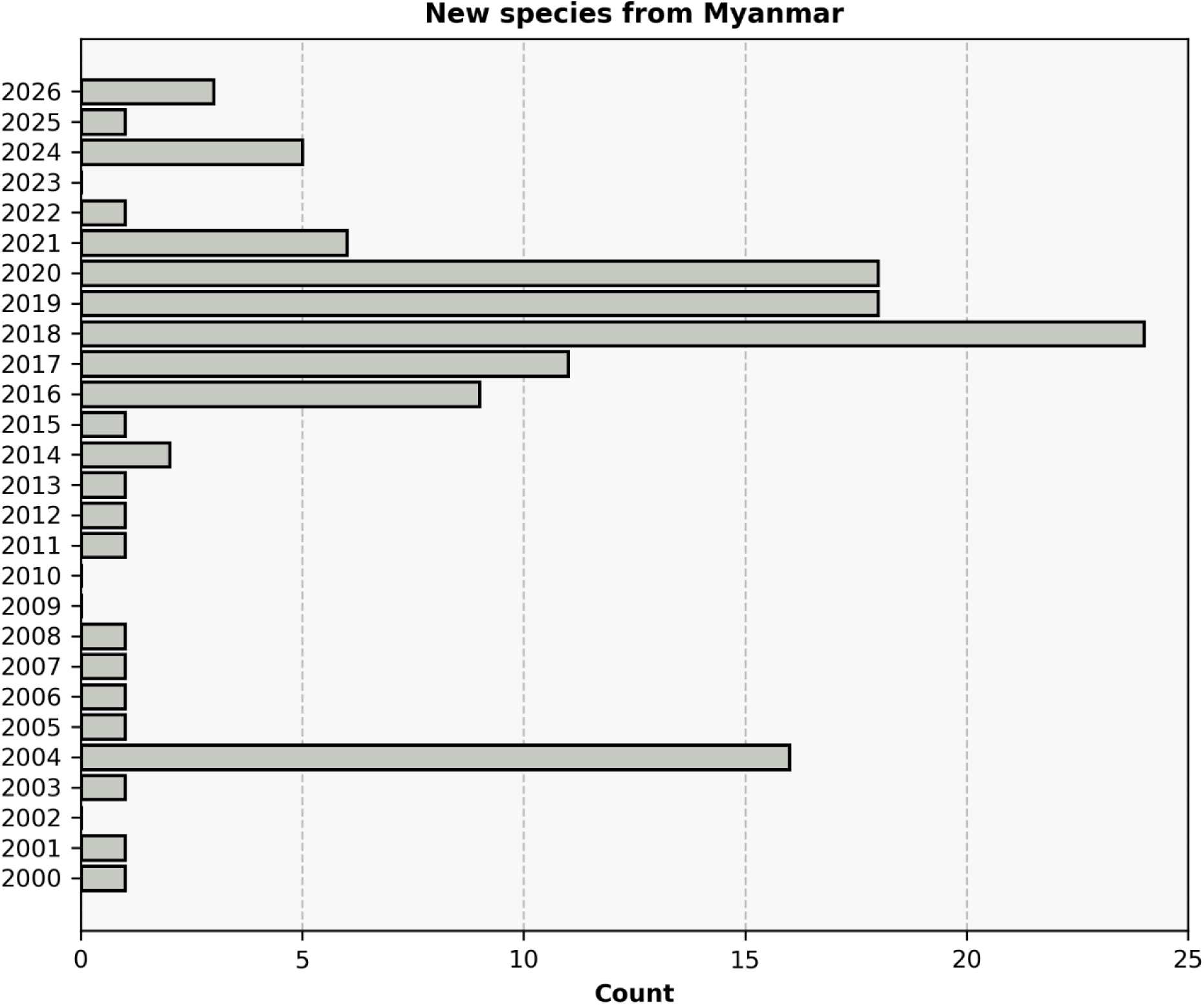
Comparison of the number of new species published from Myanmar, per year, across sampled journals.

**Figure 22.**
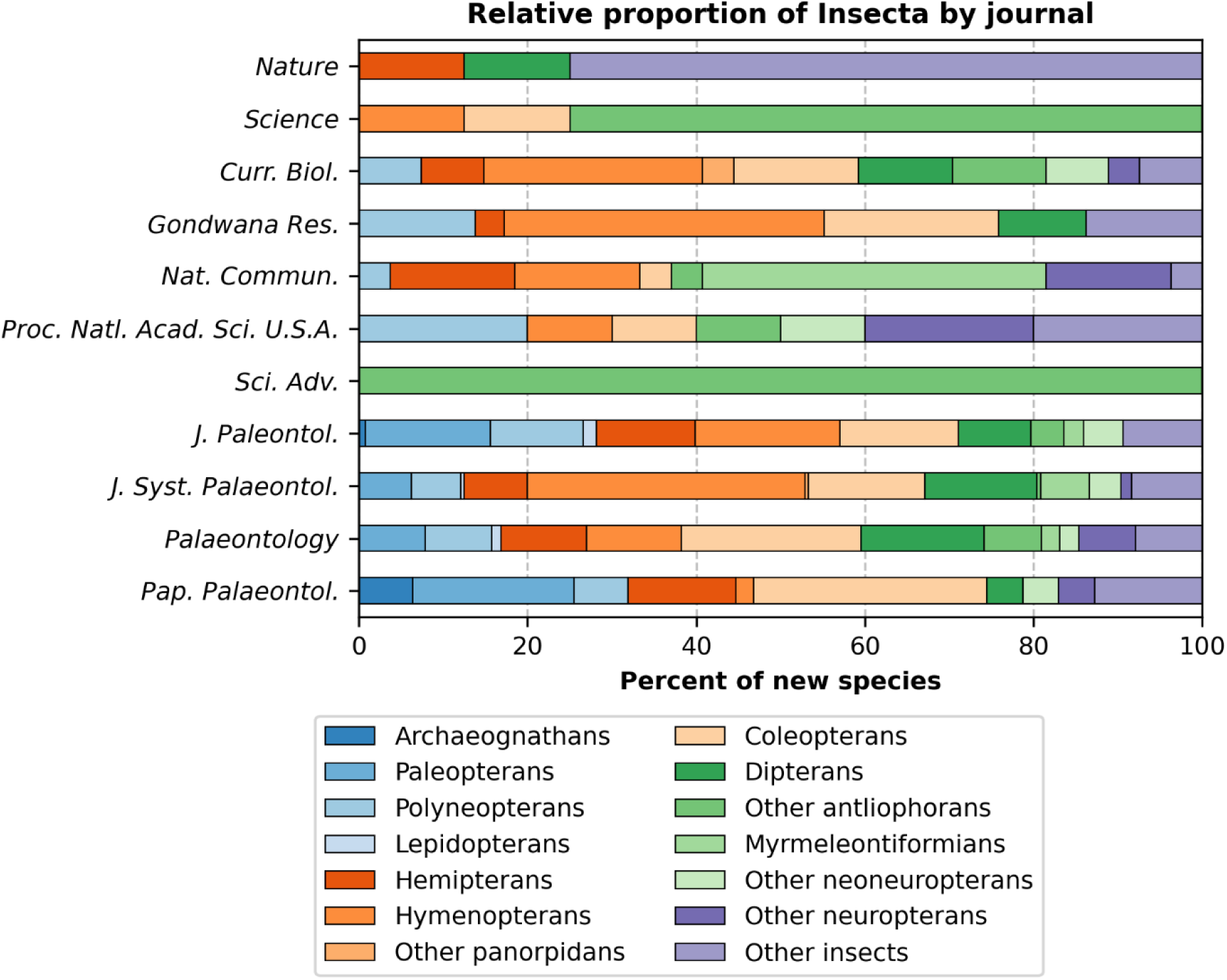
Proportional representation of different insect groups between journals. *Nature Ecology & Evolution* is omitted because no new species published in this venue were classified as insects; the *Journal of Vertebrate Paleontology* is omitted because it does not publish invertebrate material.

**Figure 23.**
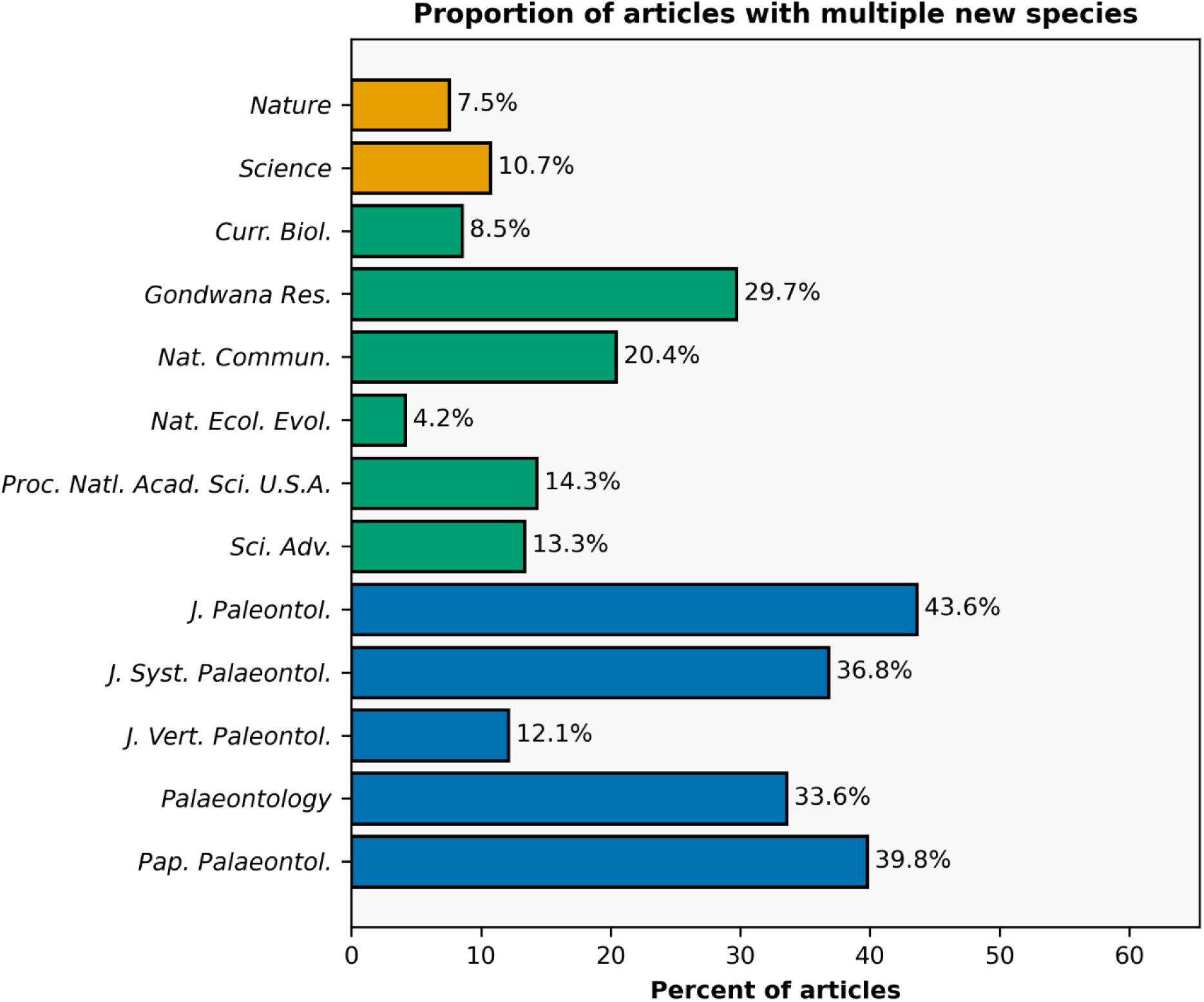
Proportion of naming papers published in each journal that name multiple (>1) new species. Colors correspond to the tripartite grouping of journals used in this study (yellow = high-JIF, green = medium-JIF, blue = low-JIF).

**Figure 24.**
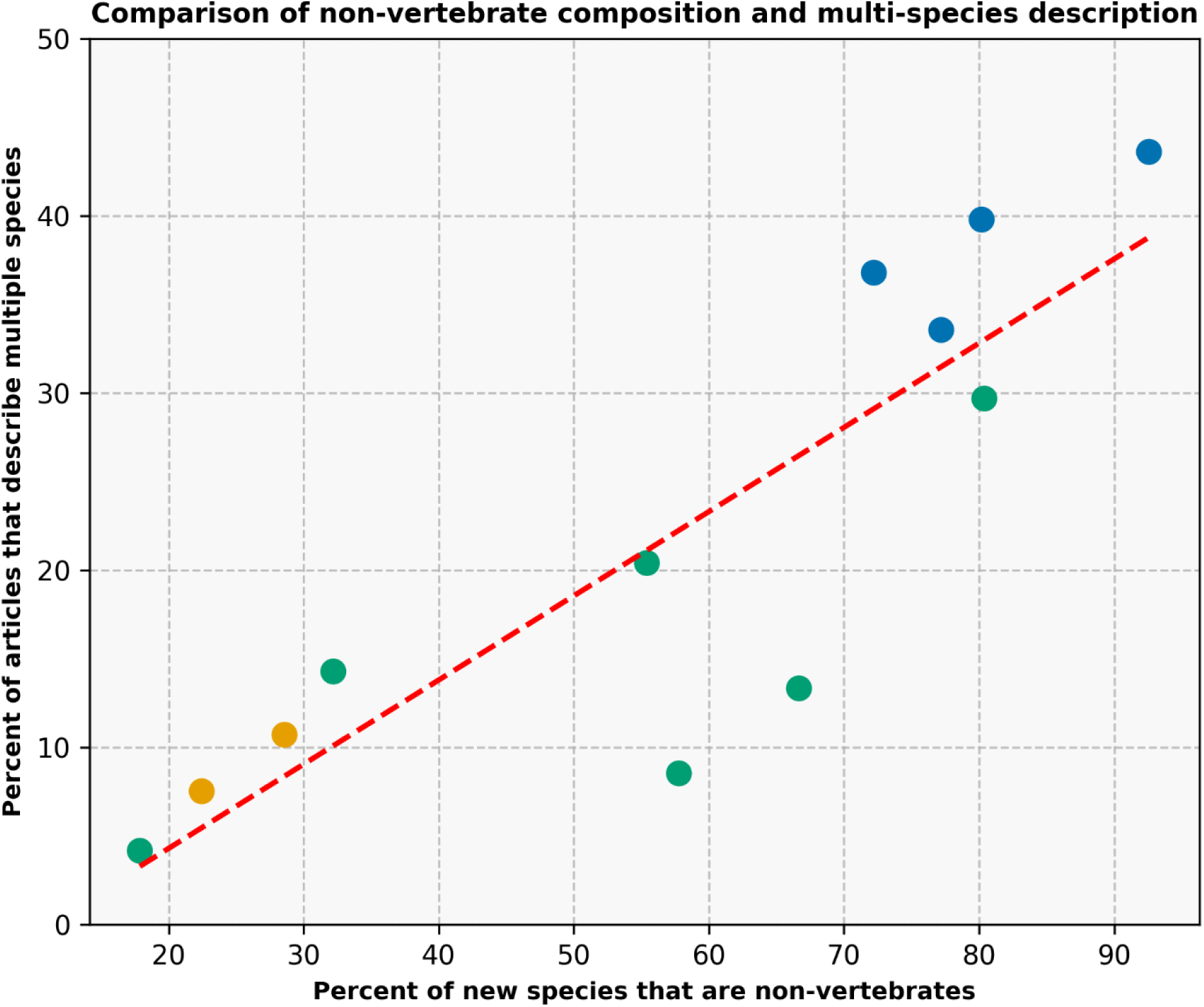
Comparison of the percent of new species that are non-vertebrates in a journal versus the percent of naming papers in that journal that describe multiple new species. The Journal of *Vertebrate Paleontology* is omitted since it does not describe non-vertebrates (includes both invertebrates and non-metazoans). Colors correspond to the tripartite grouping of journals used in this study (yellow = high-JIF, green = medium-JIF, blue = low-JIF).

The trend for naming multiple new species in one paper can be compared across journals (Figs. 23–24). Firstly, medium- and -high-JIF journals show lower rates of multi-species description; *Gondwana Research* may be relatively high due to the large proportion of pan-arthropods, while the *Journal of Vertebrate Paleontology* likely has a much lower proportion than the four other control journals and some of the higher JIF journals due to the vertebrate focus (Fig. 23). A strong correlation is identified between the percent of new species in a journal that are non-vertebrates and the percent of articles in that journal that describe multiple species (*r^2^* = 0.75; Fig. 24).

The fourth case study pertains to amphibians (a polyphyletic grouping of non-amniote tetrapodomorphs as defined here; Fig. 25). This group comprises just 1.4% of new species in the dataset, ranking it 17^th^ by total species count (*n* = 114; Fig. 1) but represents 5.9% of new species in *Nature* and 4.3% of new species in *Science* (Fig. 5). In the smaller sample size of *Nature Ecology & Evolution*, amphibians represent 17.9% of new species (Fig. 9). Closer examination reveals that this overrepresentation is largely due to the moderately charismatic stem tetrapods (non-tetrapod tetrapodomorphs) that capture important transitions in the origin of tetrapods and the colonization of land such as *Tiktaalik* (Daeschler et al., 2006 [*Nature*]), as well as crown tetrapod taxa often lumped under ‘early tetrapods,’ such as *Pederpes* (Clack, 2002 [*Nature*]) and *Parmastega* (Beznosov et al., 2019 [*Nature*]). As previously mentioned, the single assemblage description by Clack et al. (2016) in *Nature Ecology & Evolution* accounted for five new species.

**Figure 25.**
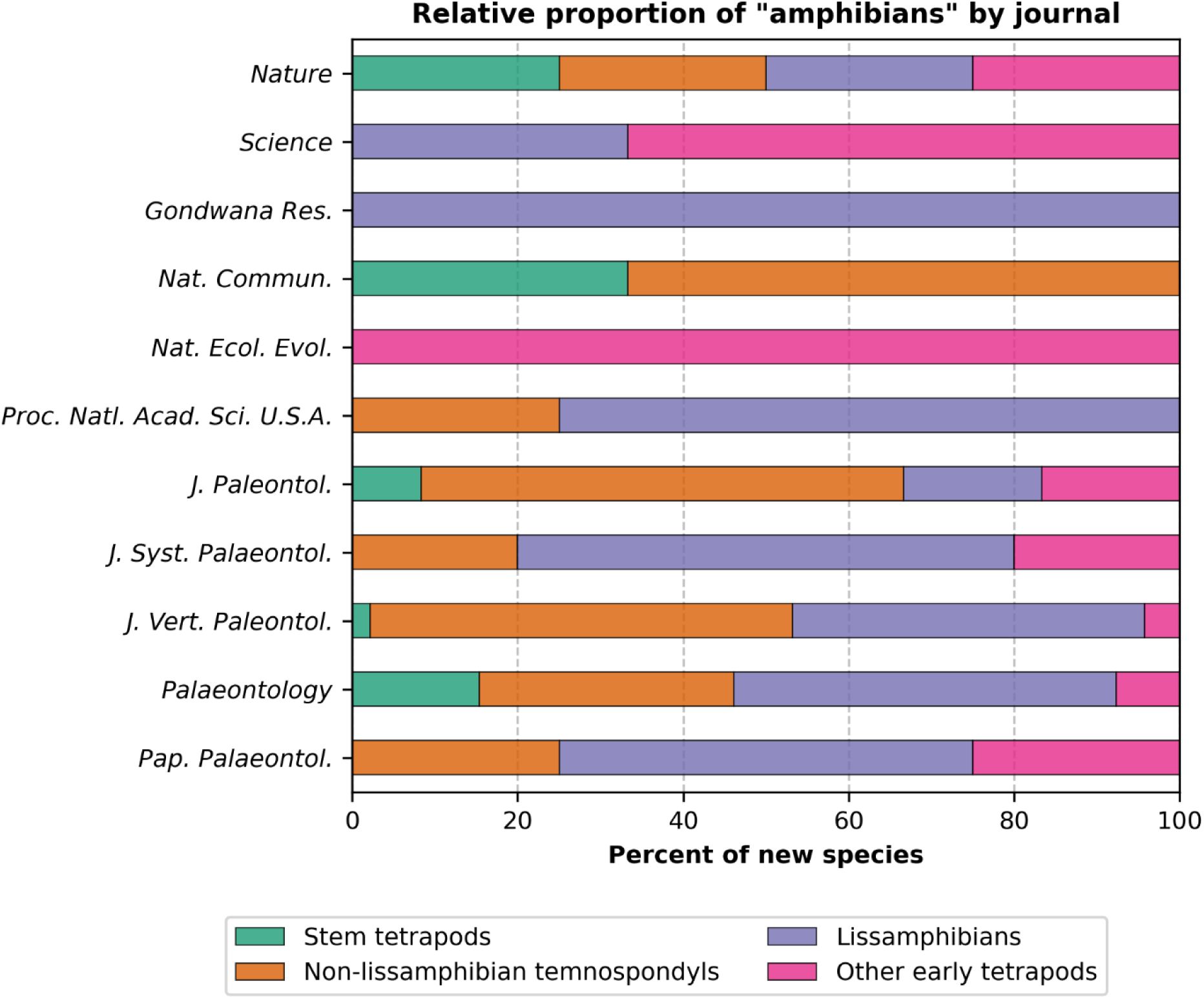
Proportional representation of different amphibian (sensu lato) groups between journals. *Current Biology* and *Science Advances* are omitted because no new species published in these journals were classified as amphibians.

The last case study is the catch-all ‘Other’ category, which constitutes a small proportion in high-JIF journals but a much more sizeable proportion in the low-JIF journals (Figs. 4, 12–13). Here, this category is further subdivided into additional groups (Fig. 26). The low-JIF journals, for which this category is proportionately larger, show a greater representation of clades that are fairly well-known among paleontologists but probably not among the public, such as acritarchs, dinoflagellates, and radiolarians, as well as some that are well-known broadly but not charismatic (e.g., annelids). For the high-JIF journals, each of which has only a handful of species in this category, otherwise unclassified chordates (e.g., tunicates), annelids, chaetognaths, and further uncategorized taxa are the most common. The last of these mainly comprises metazoans of uncertain placement (e.g., the bilaterian *Yilingia spiciformis*, Chen, Zhou, Yuan, & Xiao, 2019) or that belong to a clade with few species in this dataset (e.g., the lobopodian *Diania cactiformis* Liu, Steiner, Dunlop, Keupp, Shu, Ou, Han, Zhang, & Zhang, 2011).

**Figure 26.**
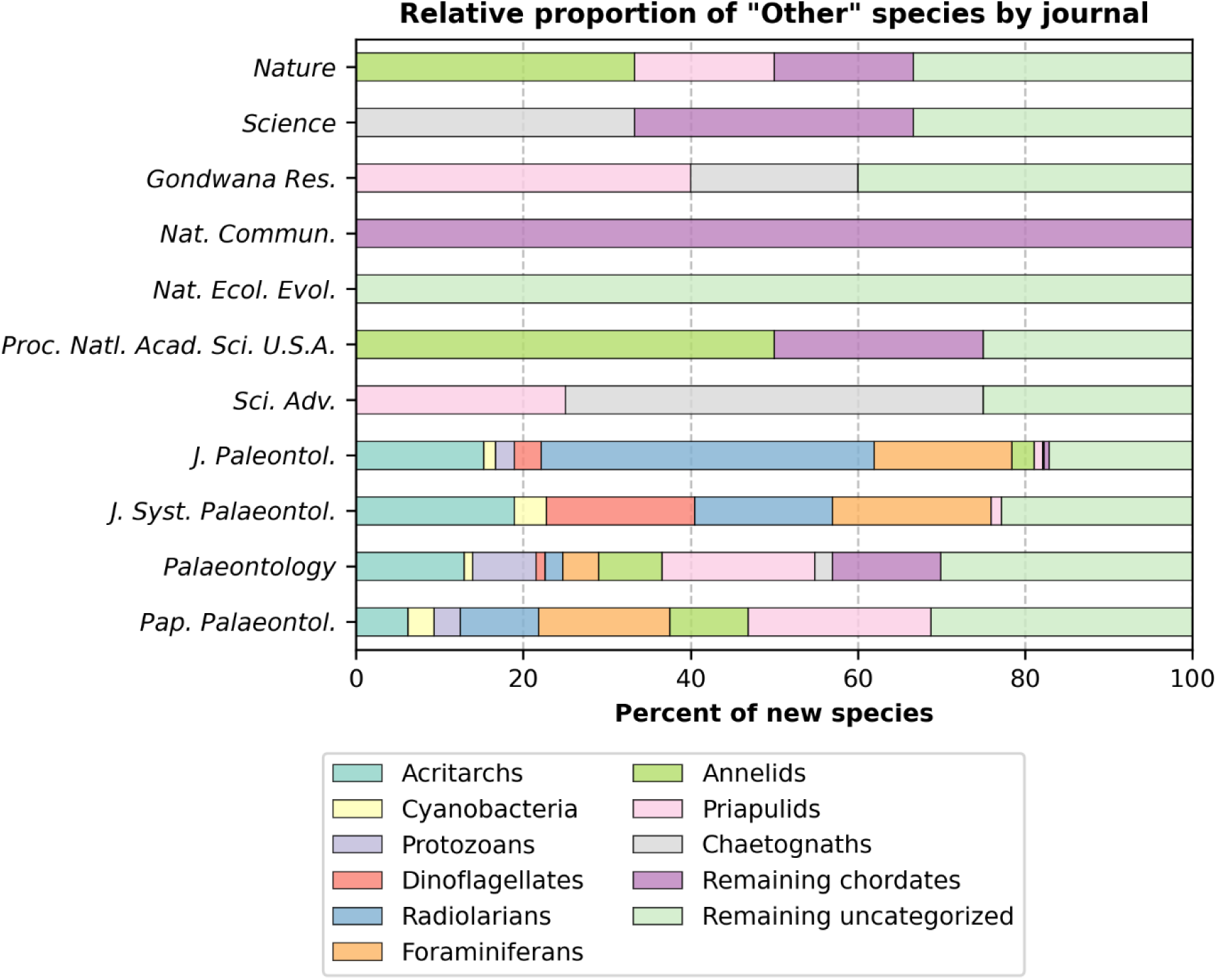
Proportional representation of further discretized groups within the ‘Other’ catch-all category between journals. The *Journal of Vertebrate Paleontology* is omitted because no new species published in this journal were classified as ‘Other.’ All high-JIF journals other than *Nature* (*n* = 6) have fewer than six species categorized as such.

## Discussion

### There are clear disparities in taxonomic representation between journals

The data presented here provide quantitative support for the common anecdotal belief that certain charismatic groups of extinct organisms, particularly the clades including dinosaurs and mammals (Fig. 27), are more likely to be published in prestigious journals. In the sample analyzed here, avemetatarsalians (inclusive of avians) comprise less than 5% of new species in the control-group journals, yet they comprise 15.7% of new species published in *Science* and 23.9% of new species published in *Nature.* Likewise, mammaliamorphs (inclusive of primates) comprise less than 11% of new species in the control-group journals, yet they comprise 40% of new species published in *Science* and 31.7% of new species published in *Nature*. Relatively high proportions of these groups are also found in most of the middle-JIF journals except for *Gondwana Research*. The high proportion of these groups in the *Journal of Vertebrate Paleontology* is undoubtedly in part the result of its vertebrate-only focus and a high ratio of non-primate mammaliamorphs to primates compared to higher-JIF journals (Fig. 27). For *Nature* and *Science*, the combination of all avemetatarsalians and all mammaliamorphs comprise more than 50% of all new species published since 2000.

**Figure 27.**
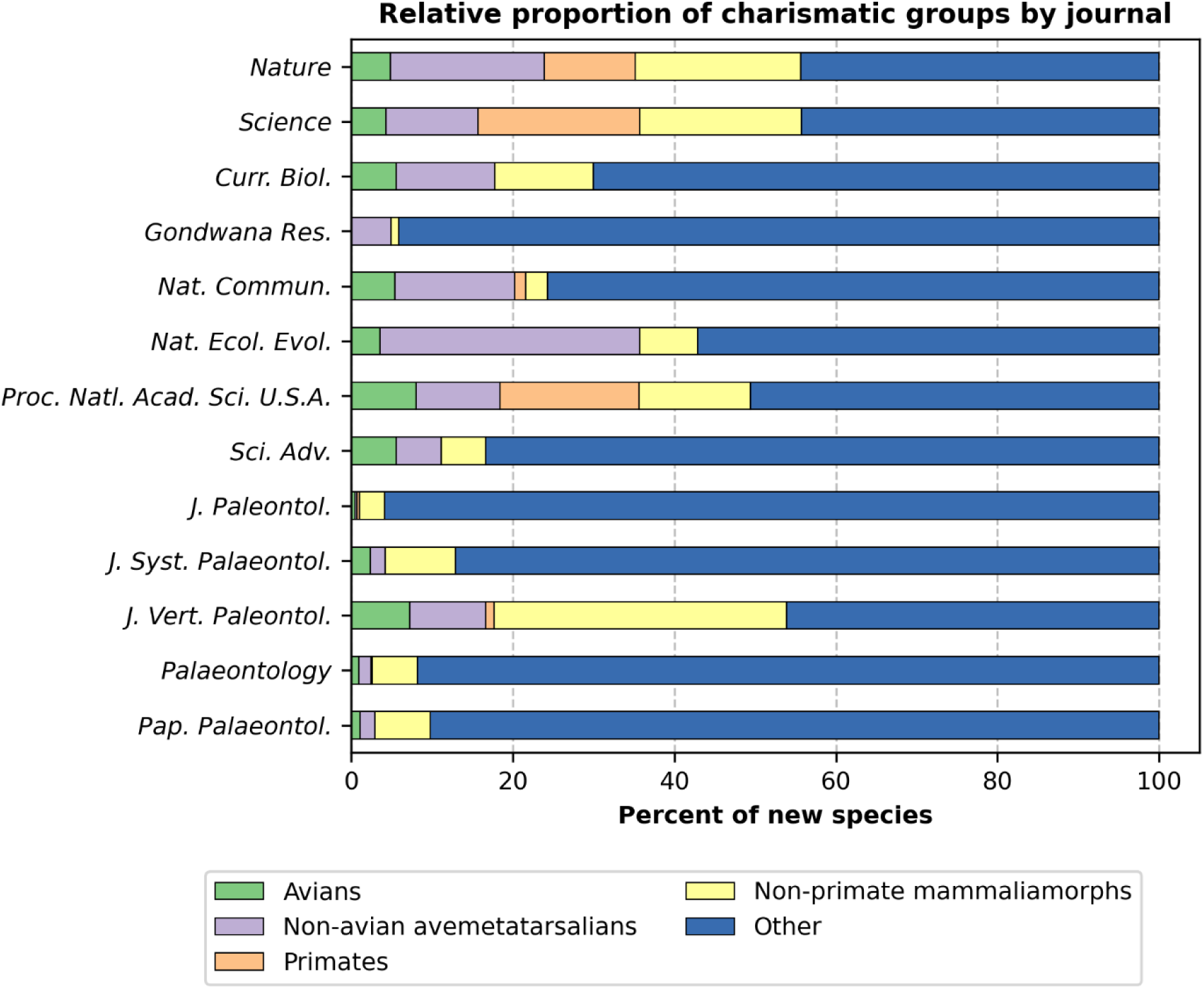
Distribution of charismatic taxonomic categories between journals. Avians, non-avian avemetatarsalians, primates, and non-primate mammaliamorphs are retained as separate categories, while all other taxa are lumped under the ‘Other’ category.

In general, most vertebrate categories (usually with the exception of fish) tend to be overrepresented in high-JIF journals. Figures 28 and 29 visualize the relative over- or underrepresentation of each category by comparing the proportion that each of the 20 categories comprises in the high- and medium-JIF controls when compared to the control journals.

**Figure 28.**
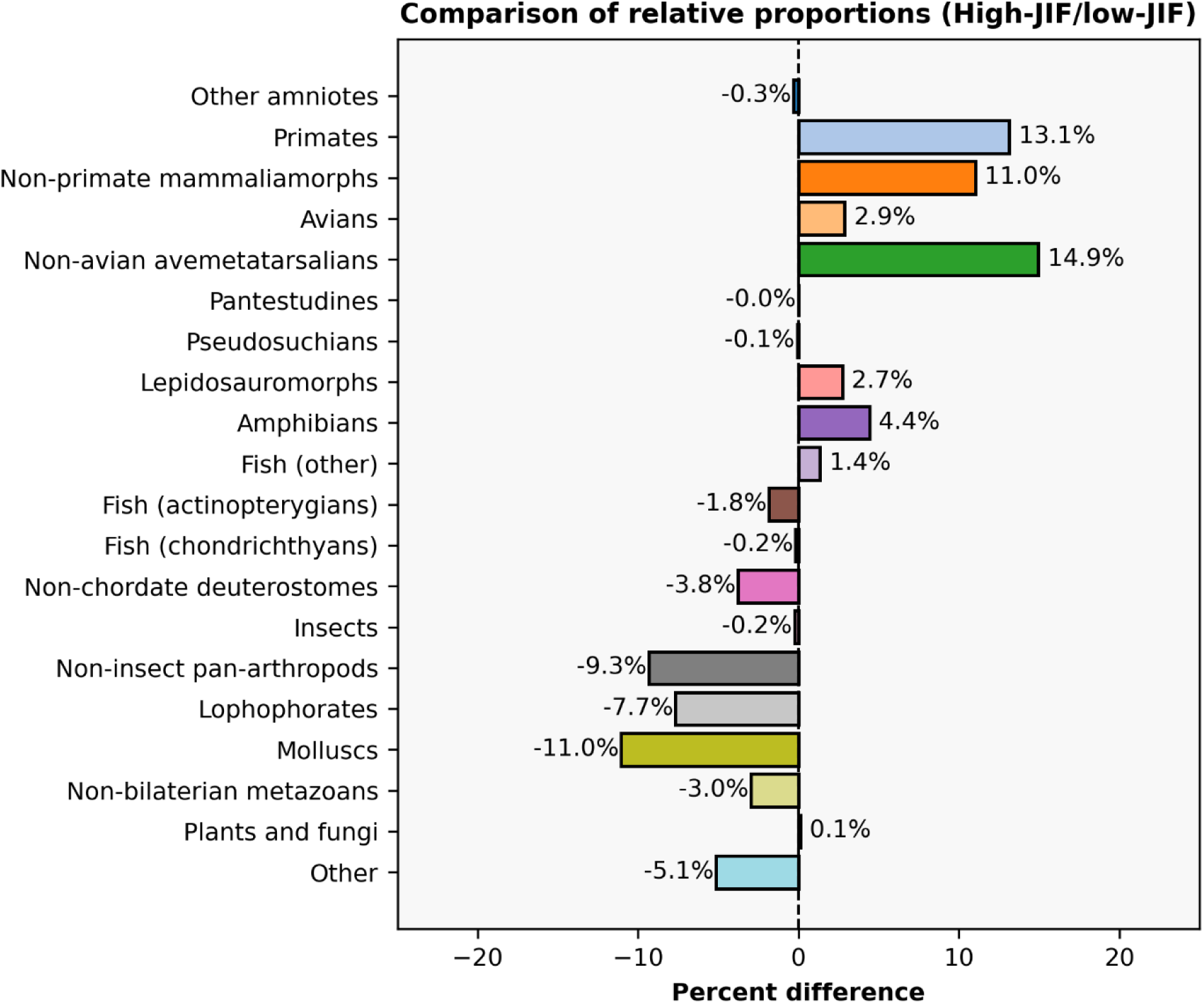
Comparison of proportional representation of taxonomic categories between *Nature*/*Science* and the control-group journals. The values represent the difference between the percent of new species in each group (non-control minus control). Positive values (to the right) indicate that a new species in that category constitute a higher percent of all species published in *Nature*/*Science* than they do in the control journals (i.e. proportional overrepresentation, as with primates); negative values (to the left) indicate the inverse (i.e. proportional underrepresentation, as with molluscs).

**Figure 29.**
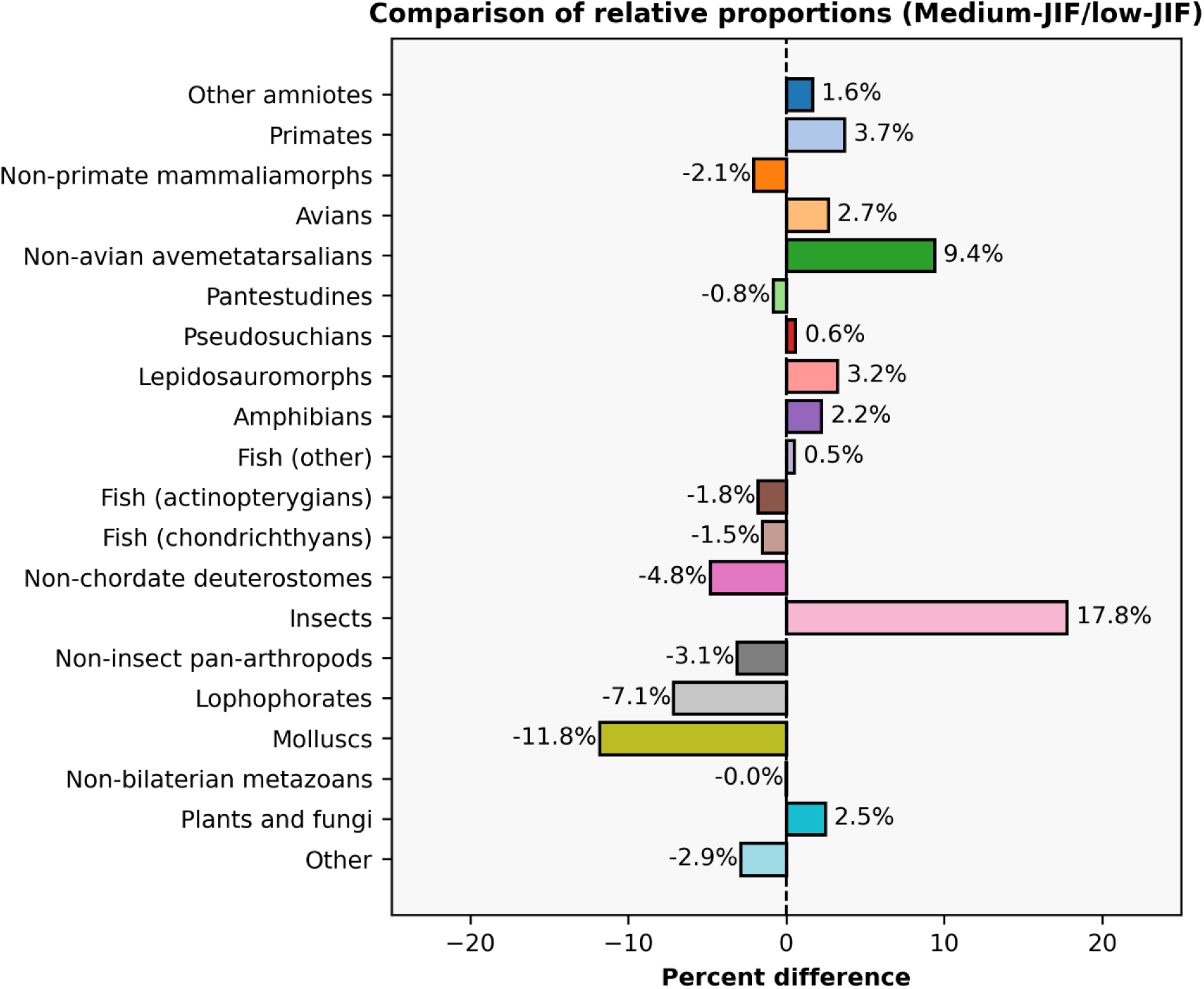
Comparison of proportional representation of taxonomic categories between Current Biology/Gondwana Research/Nature Communications/Nature Ecology & Evolution/Proceedings of the National Academy of Sciences/Science Advances and the control-group journals. The values represent the difference between the percent of new species in each group (non-control minus control). Positive values (to the right) indicate that a new species in that category constitute a higher percent of all species published in the non-control journals than they do in the control journals (i.e. proportional overrepresentation, as with insects); negative values (to the left) indicate the inverse (i.e. proportional underrepresentation, as with molluscs).

Vertebrates’ overrepresentation comes primarily at the expense of invertebrates (rather than non-metazoans), which represent over 60% of new species (Fig. 1). For example, no new species of lophophorates (8.5% of the total sample) or molluscs (13.8% of the total sample) have been published in *Science* in this century (though there have also been no new species of turtles or pseudosuchians in this venue either; Fig. 5). Actinopterygian and chondrichthyan fish are also proportionately underrepresented in *Nature* and *Science*; the ‘Fish (other)’ category, which represents 5.9% of new species in the former (Fig. 5), comprises mainly Paleozoic taxa that capture unusual or noteworthy features in the evolution of gnathostomes and towards the tetrapod lineage (e.g., the placoderm *Qilinyu* Zhu, Ahlberg, Pan, Zhu, Qiao, Zhao, Jia, & Lu, 2016 [*Science*]). These trends generally hold true when examining the medium-JIF journals (Fig. 29), with the exception of non-primate mammaliamorphs (underrepresented, likely due to very strong representation in the *Journal of Vertebrate Paleontology*) and insects (overrepresented).

Finally, comparing JIF against the percent of new species in a journal that belong to either Avemetatarsalia or Mammaliamorpha reveals a positive correlation between the JIF and the percent of new species that belong to a clade designated as ‘charismatic’ here (Fig. 30). The relationship is in part skewed by the *Journal of Vertebrate Paleontology* given its vertebrate focus but relatively low JIF; removing this journal (Fig. 30B), increases the *r^2^* from 0.32 to 0.65.

**Figure 30.**
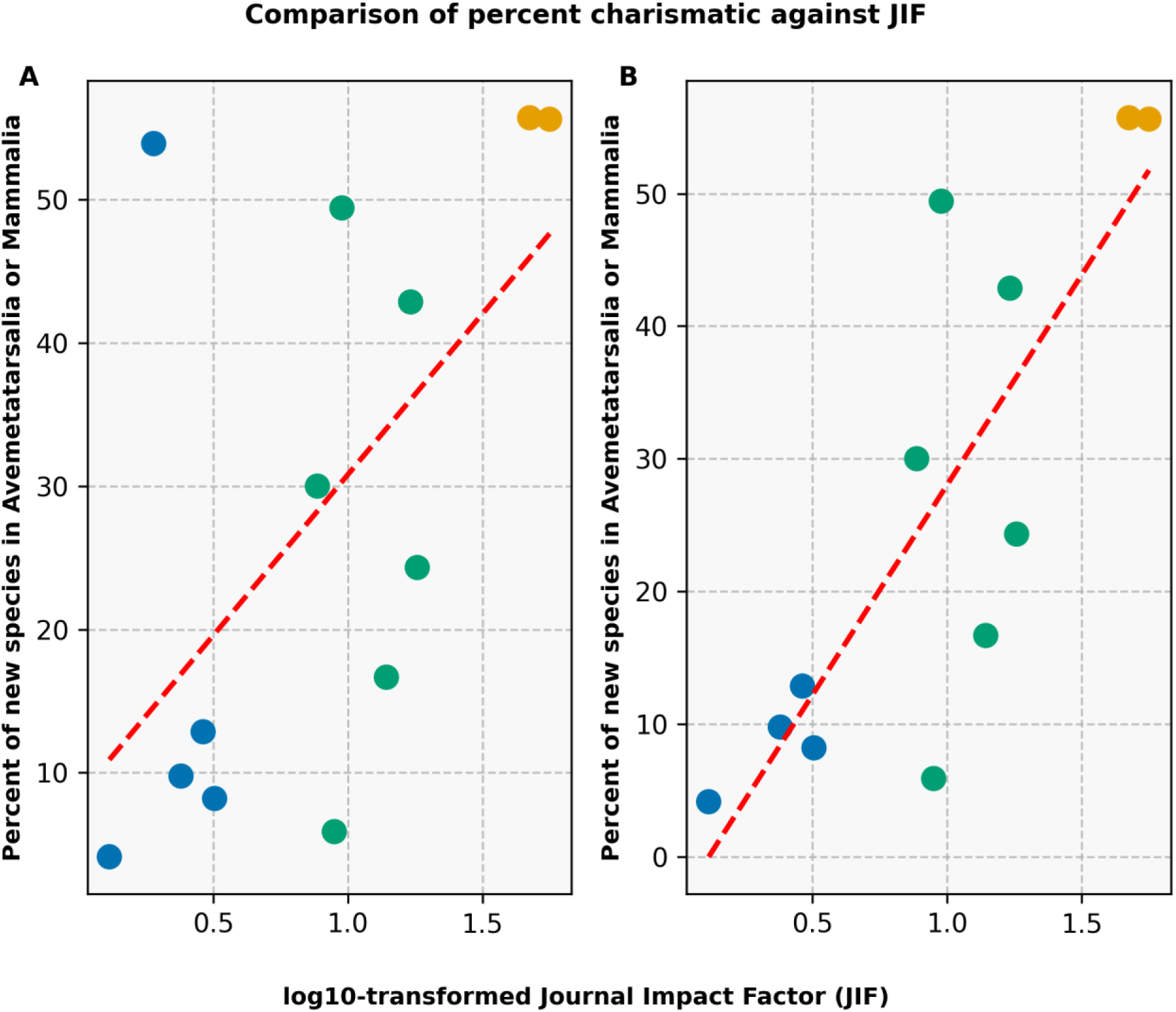
Comparison of log-10-transformed Journal Impact Factor and the percent of new species classified under Avemetatarsalia or Mammaliamorpha between journals. (A) all journals; (B) excluding the *Journal of Vertebrate Paleontology*. The dashed red lines represent a linear regression. Colors correspond to those in Figures 23–24.

### Taxonomic biases are also present in databases

As mentioned above, about 11.5% of species in this dataset (nearly 1,000) belong to genera that were not found through the PBDB API and that appear to be genuinely absent after manual examination. Although this is a clear minority of the sampled genera, it could alternatively be framed as a rather grim observation that ∼1 in 9 genera with a new species published in the last quarter-century are missing from a database that aspires to comprehensively sample extinct organisms. Considering the pervasive use of PBDB data, anthropological gaps are concerning as they compound existing preservation and sampling biases. These gaps are also somewhat surprising given that all of the sampled journals are well-known and widely indexed, as compared with journals in which new species may be published but not be as discoverable (e.g., new journals, non-English journals, journals that do not use DOIs). Although some delay in adding new data to the PBDB is to be expected, the vast majority of these species (*n* = 897 [92.95%]) were published prior to 2026, which as of the time of API retrieval, means that the names had existed for at least eight months; more than 40% (*n* = 416) were published prior to 2016, or more than a decade ago. Additionally, some of the genera not found in the PBDB are decades if not centuries old (e.g., Zmudzinski [2020] named two new species of the mite *Neophyllobius* Berlese, 1886).

Examination of the patterns of missing data among the 20 categories created for this study reveals similar patterns to those observed for the broader inter-journal comparisons (Fig. 31). In general, non-vertebrates have higher missing taxa rates than vertebrates (though non-primate mammaliamorphs also had a fairly high proportion of missing genera). Charismatic categories such as avians, non-avian avemetatarsalians, and primates had very low rates of missing genera. The catch-all ‘Other’ category was particularly high as well due to representation of taxa such as acritarchs and various Proterozoic microfossils that are more likely to lack precise taxonomic placement and that are certainly not regarded as charismatic. An important consideration is that differences between clades may not only result from general charisma or lack thereof but also from the taxonomic specialization of different contributors to the PBDB; this might account for why some invertebrate groups have relatively low missing genera rates and why non-primate mammaliamorphs have relatively high missing genera rates.

**Figure 31.**
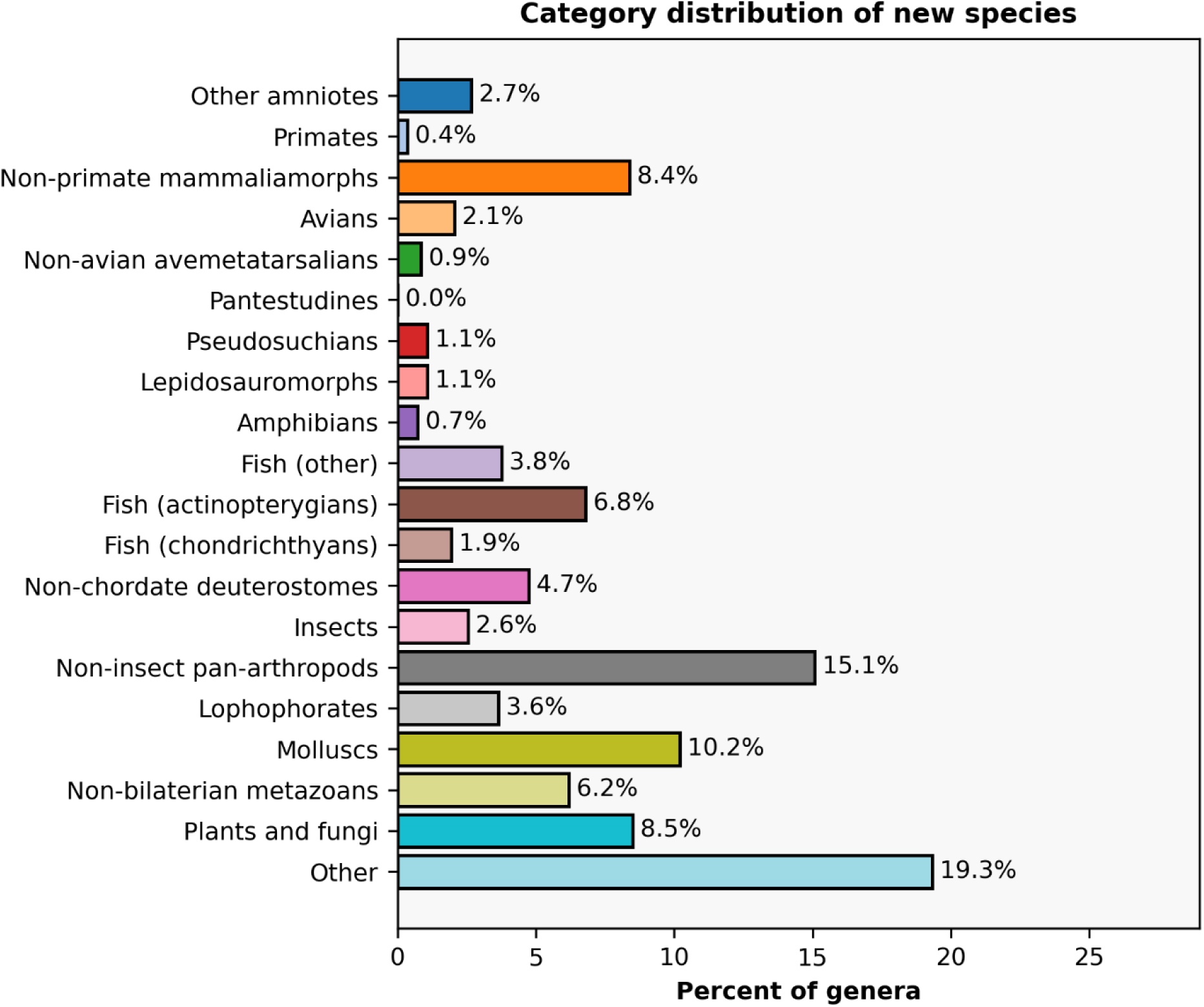
Distribution of taxonomic categories for unique genera with at least one new species that was published in any one of the journals sampled here and for which the genus was not found in the Paleobiology Database (PBDB) as of September 8, 2026. This excludes genera that failed to be retrieved programmatically but that were found manually and those that were retrieved the PBDB but with little to no rank information (regardless of whether that rank information needed to be augmented further). Genera listed in the PBDB but as subgenera (less than 15) are included. Multi-species genera are represented only once. Categories are ordered roughly by phylogenetic divergence, with plants, fungi, and invertebrates towards the bottom, fish in the middle, and amniotes at the top.

Comparison of the difference in proportions that a category represents of all unique genera in the dataset and those genera missing from the PBDB further supports taxonomic bias in missing data (Fig. 32). Several clades have differences in proportions exceeding 2%, including the grouped plants and fungi, actinopterygian fish, and the catch-all ‘Other.’ Conversely, the more charismatic clades like non-avian avemetatarsalians and non-primate mammaliamorphs are relatively underrepresented (so too are non-insect pan-arthropods and lophophorates).

Disparate attention is also evinced when comparing what proportion of unique articles that have at least one species without a genus listing in the PBDB are comprised by each journal (Fig. 33). The control journals comprise the vast majority (>88%) of such articles, with the *Journal of Paleontology* representing more than a third of all such articles, although it bears noting that their relatively large sample size also increases the likelihood for novel taxa to not be in the PBDB.

**Figure 32.**
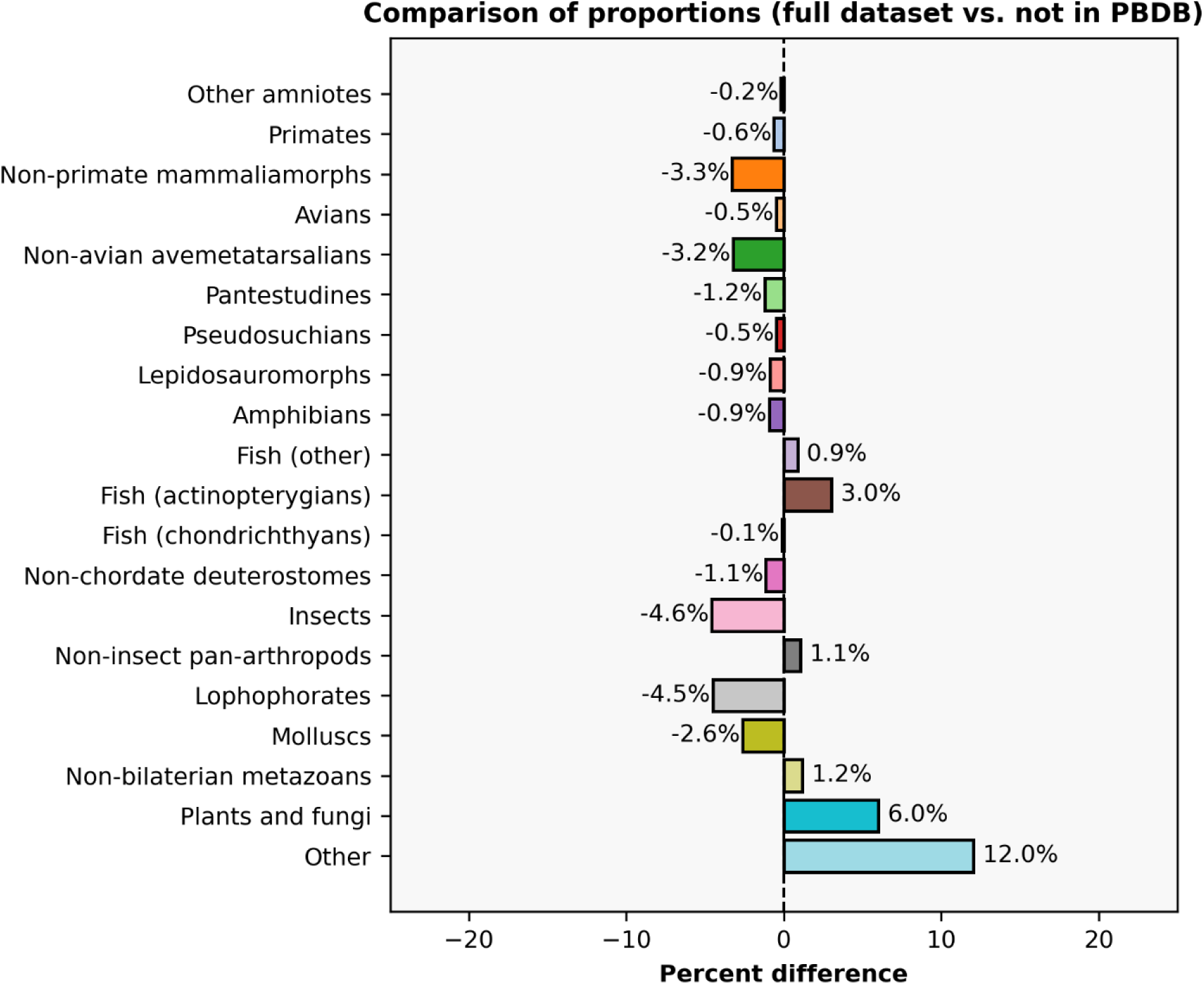
Comparison of proportional representation of taxonomic categories between all sampled journals and the subset of unique genera missing from the PBDB. The values represent the difference between proportion of all unique genera in the dataset that a category comprises (after manual augmentation for missing values) and the proportion of unique genera missing from the PBDB that a category comprises. Positive values (to the right) indicate that a given category comprises a higher proportion of the missing PBDB entries (i.e. proportional overrepresentation, as with actinopterygians); negative values (to the left) indicate the inverse (i.e. proportional underrepresentation, as with insects).

**Figure 33.**
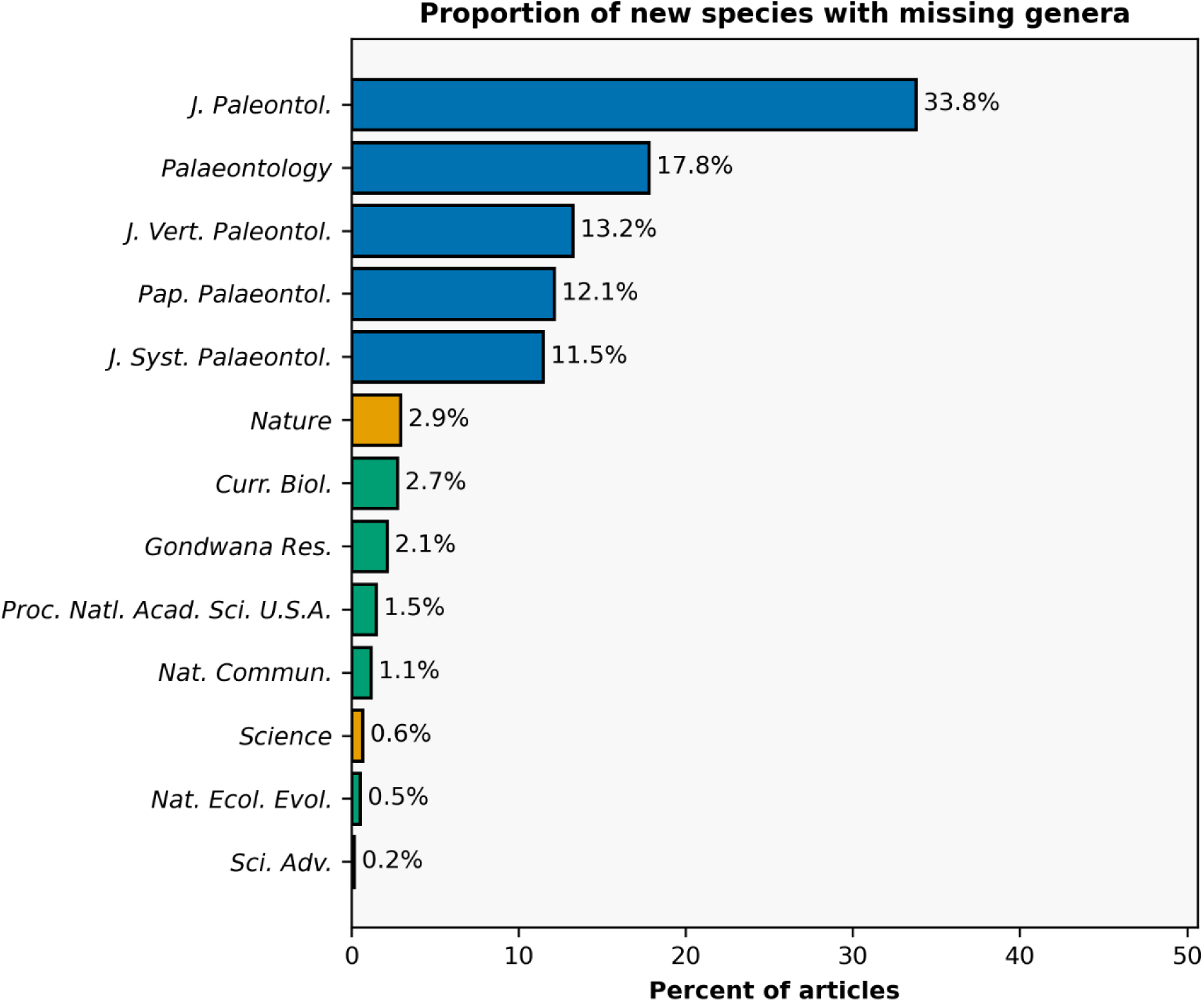
Comparison of proportion of unique articles with at least one new species for which the genus is missing in the PBDB. The proportion visualized represents each journal’s proportion out of the total subset of articles (e.g., 33.8% of all articles with at least one missing genus were published in the *Journal of Paleontology*). Unique articles are the metric used here rather than unique genera or species to avoid overweighting single papers that might contain many new species (with or without missing genera). Colors correspond to the tripartite grouping of journals used in this study (yellow = high-JIF, green = medium-JIF, blue = low-JIF).

Likewise, intra-journal comparisons (what percent of each journal’s articles contain at least such genus) reveals a similar (though less consistent) pattern (Fig. 34). On balance, control journals tend to have higher proportions of articles for which some genera are missing from the PBDB; higher-JIF journals like *Current Biology* and *Gondwana Research* with high proportions could be linked to their relatively invertebrate-skewed samples compared to other high-JIF journals. Interestingly, the *Journal of Vertebrate Paleontology* has a relatively low proportion among control journals that is even lower than that of *Nature*. This could be taken as further evidence for the general charisma of vertebrates relative to non-vertebrates and a correlation between missing data and the ratio of vertebrates to non-vertebrates among sampled naming papers.

**Figure 34.**
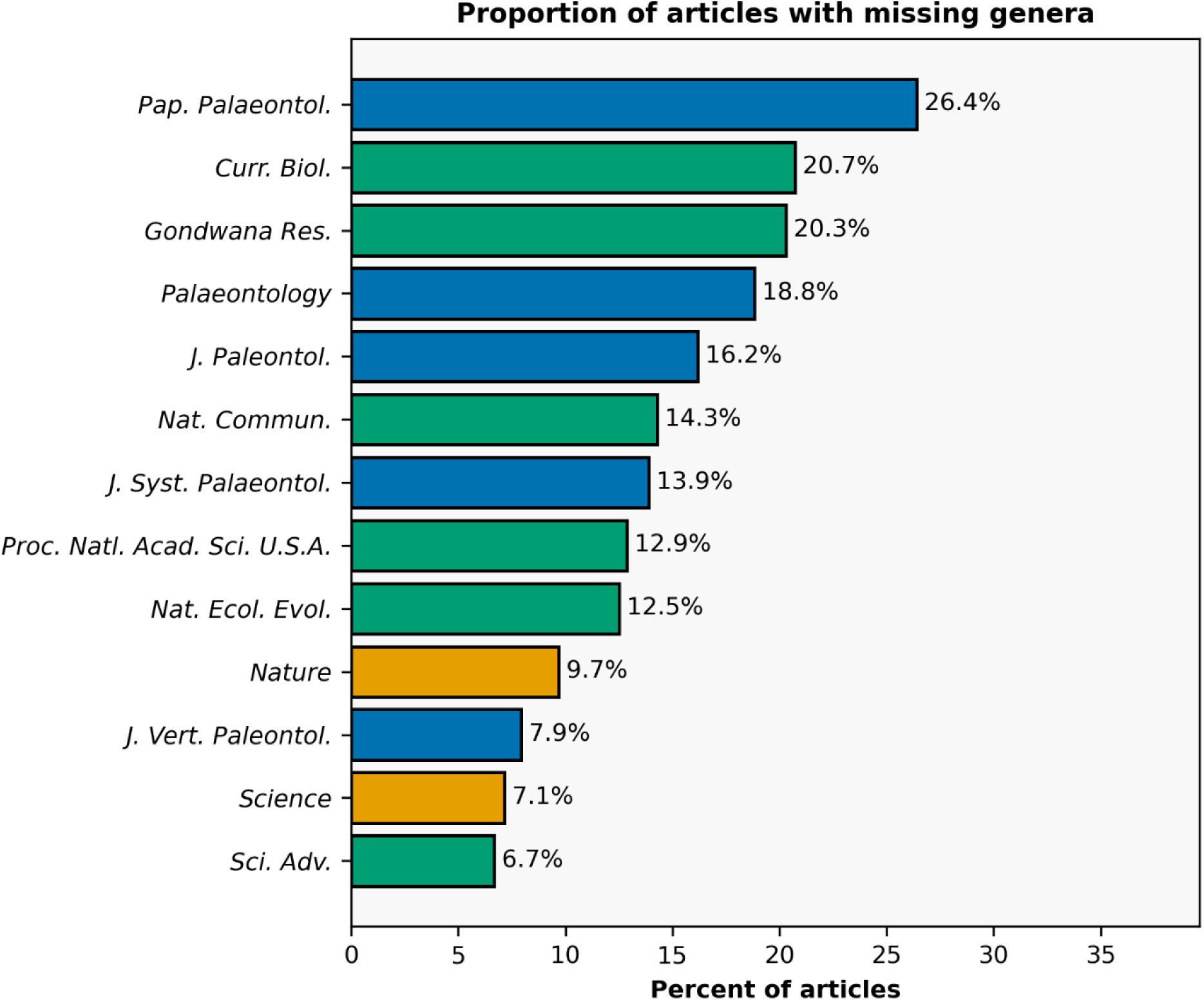
Comparison of proportion of unique articles with at least one new species for which the genus is missing in the PBDB. The proportion visualized represents each journal’s proportion out of its total number of articles in the dataset (e.g., 16.2% of all articles in the *Journal of Paleontology* include a new species placed in a genus that is missing). Unique articles are the metric used here rather than unique genera or species to avoid overweighting single papers that might contain many new species (with or without missing genera). Colors correspond to the tripartite grouping of journals used in this study (yellow = high-JIF, green = medium-JIF, blue = low-JIF).

As this study is not focused on assessing quality of large databases, only a few remarks on these results are presented here. Issues or deficiencies with PBDB data for certain clades have been previously discussed (e.g., Prothero, 2015; Mondal & Harries, 2016; Zizka et al., 2019; Schroeder et al., 2022; Greenwalt et al., 2025; Cui et al., 2026), though whether errors in large datasets in general affect the general results of downstream studies remain disputed (e.g., Benton, 1999; Adrain & Westrop, 2000; Wagner et al., 2007; Allmon et al., 2017; Gardiner et al., 2026). The deficiencies noted here are probably not as bad as they appear, as a range of taxonomic acts subsequent to the original description may have altered the binomial of a recently named species. For example, some genera may have been synonymized with others, so the original genus in which a new species was placed and is listed in the dataset may no longer be valid. However, the PBDB can and should record both the junior synonym and its relationship, which it does for some but hardly all of the genera sampled here, so missingness due to synonymy is not ideal, even if it is an explanator. Because taxonomic identifications and frameworks are hypotheses, not immutable facts, these can change over time, sometimes frequently, and at least some of a database’s utility is tied to its ability to record not just the current but also the past taxonomy of a given taxon (e.g., Markwick & Lupia, 2002).

This inadvertent case study does indicate that for some clades, the assumption that the PBDB faithfully approximates the known fossil record is potentially not true, beyond existing biases related to making data public that are more challenging to resolve, such as ‘dark data’ (e.g., Marshall et al., 2018; Dean & Thompson, 2025). If taxa are not represented in the PBDB, their occurrences, the more common datapoints analyzed by re-users, almost certainly are not either. This point underscores the importance of data entry and curation in large, taxon-agnostic databases by workers with taxonomic, stratigraphic, and spatiotemporal expertise (e.g., Allmon et al., 2018; Guinot et al., 2026) and the need for potential users of databases to critically assess their records of interest and to take action as necessary (e.g., Jones et al., 2025), which includes a determination of whether the database is appropriate for the questions being asked.

### The causal mechanism(s) of taxonomic bias remain unclear

Although this study has demonstrated a clear correlation between a journal’s profile/prestige (as measured by one commonly-used index, the Journal Impact Factor [JIF]), it has not demonstrated a causal mechanism. In other words, the taxonomic bias observed here is not clearly the direct result of an editorial bias against taxa, even if this is a reasonable hypothesis. The primary challenge in testing for causality is that doing so requires detailed information on submissions made to journals, including details on why certain submissions were or were not sent out for review and, for those that were reviewed, why they were or were not accepted for publication. This information is highly unlikely to be made available by journals. Despite some aspects of publishing becoming more transparent, such as open peer review (e.g., Wolfram et al., 2020), which usually includes editorial decision letters, this cannot account for manuscripts that are reviewed but ultimately rejected.

Furthermore, the practice of desk rejecting submissions (declining to send them out for review) plays a significant role at prestigious journals and is likely to become even more utilized given the meteoric rise in scholarly outputs (e.g., Fox et al., 2017; Aczel et al., 2021; Severin & Chataway, 2021; Hanson et al., 2024) and the increased usage of generative AI that has enabled faster manuscript generation (e.g., Maupin et al., 2025; Watson, 2025; Spitzer, 2026; Gartenberg et al., 2026). For example, *Science* only sends out 15% of all submissions for review, and *Nature* says that “most” submissions are desk rejected (American Association for the Advancement of Science, n.d.a; Nature Portfolio, n.d.a). The point at which submissions are being considered for external peer review is likely a critical pinch-point in which bias against taxa might appear, as editors screening manuscripts are less likely to have specialist expertise that reviewers would. In discipline-agnostic journals, paleontology submissions may not even be handled by paleontologist. It is easy to conceive a situation like this where a more charismatic taxon is recognizable to a non-expert, but a less charismatic one is not, and that this can sway an editor’s perception of the broader relevance/interest of a submission. However, desk rejection is even more opaque than standard peer review, and it is considered unlikely that granular data on this practice would ever be shared.

Another possible explanation is that researchers, particularly those who specialize in less charismatic taxa, are self-selecting against submissions to higher-profile, more selective journals based on their own perception of the likelihood of acceptance of their work on the basis of their study system. The causal mechanism here would be researchers’ behavior, not editors’ behavior. If clades are not represented in submissions, they cannot be represented in publications. Self-selection is a compelling hypothesis because decisions about where to submit manuscripts are intrinsically strategic, particularly when considering highly selective venues. The costs associated with reformatting manuscripts for resubmission to different journals are well-documented (e.g., Jiang et al., 2019; Clotworthy et al., 2023), which means that aiming for a very high-JIF journal and progressively “working down” through lower, but still high, JIF journals, can carry significant costs, especially for more career-unstable researchers.

If taxonomic bias arises at least in part from self-selection, it does not mean that researchers are baselessly self-selecting or that journals are not indirectly contributing to this behavior. Most journals are selective to some degree, at minimum with respect to their topical scope, and even relatively low-JIF journals (JIF < 3.0) can and do desk-reject submissions for reasons such as a perceived lack of broader relevancy/interest. Criteria such as this are more subjective than more unambiguous flags such as inappropriate scope, poor formatting and writing, or ethical violations. Researchers’ experience submitting to less selective journals might shape their impressions of where their work is likely to be published, especially if they feel that an editorial decision like a desk rejection was in part related to a lack of (perceived) broader interest in their study system. Both this point and the broader remark about self-selection are made in part based on my own experience publishing (or attempting to publish) as a specialist of a non-charismatic clade (temnospondyls).

### Limitations of this study

There are undoubtedly many attributes of a given submission that influence its likelihood of being accepted, ranging from the methods of the study to how a study interfaces with existing literature to the peer reviewers and their relationship (or lack thereof) with the authors. This study has only focused on one, albeit one that is often informally discussed as a potential influence: the taxonomy of the study system. A wide range of paleontological work is published in high-JIF journals, much of it involving the description of material assigned to already-named species (e.g., Chuliver et al., 2024; Haridy et al., 2025; Maidment et al., 2025, 2026; Mongle et al., 2025; O’Connor et al., 2025; Liu et al., 2026; Lu et al., 2026; Makovicky et al., 2026) and studies that address questions other than taxonomy (e.g., Manafzadeh et al., 2024; Fastovich et al., 2025; Green et al., 2025; Paterson et al., 2025; Wilson et al., 2025; Lei et al., 2026b; Ruan et al., 2026). Likewise, paleontological work beyond new species can be of interest to the broader public, such as description of material with exceptional preservation. The high quality of many specimens from the Jehol (Cretaceous) and Chengjiang (Cambrian) Biotas in China probably contributes to the frequent publishing of this material (beyond naming papers; e.g., Lei et al., 2026a; Liu et al., 2026; Zeng et al., 2026) in *Nature* and *Science*. There are more than a dozen naming papers for each biota in the dataset, which includes representatives of clades not typically regarded as charismatic, such as the plants *Archaefructus* and *Leefructus* (Sun et al., 2002 [*Science*], 2011 [*Nature*]), the lamprey *Mesomyzon* (Chang et al., 2006 [*Nature*]), and the early deuterostomes *Dianchichystis* and *Vetulocystis* (Shu et al., 2004 [*Nature*]).

Another attribute that is only briefly explored here but that warrants further investigation is whether there are patterns in authorship of paleontological studies in high-JIF journals. Attributes of researchers and research groups may also account for biases, not only taxonomic, among paleontological studies (e.g., Warnock et al., 2020; Monarrez et al., 2022; Arens et al., 2024; Valenzuela-Toro et al., 2025; Wang et al., 2025a). As it pertains to publication in high-JIF journals, there are several considerations. Firstly, taxonomic bias in high-JIF journals could, in part, be introduced by repeat publishing by the same researcher(s) or research groups; because the sample size of paleontological studies published in any such journal is relatively small, it is easier to skew the taxonomic representation than in a paleontology-specific journal, for example. The broad temporal sampling of this study was designed in part to mitigate this possibility, but some journals with small sample sizes can still have skewed results in certain contexts (see previous discussion of insect groups in *Nature* and *Science* and of ‘amphibian’ groups in *Nature Ecology & Evolution*).

Secondly, researchers who publish once in a high-JIF journal stand a good chance of being repeat authors. Ostensibly, prior publishing success confers at least an indirect advantage for future submission (e.g., Sekara et al., 2018) through indirect avenues such as higher reputational standing and building the skillset necessary for submissions to high-profile journals, even if prior publication is not a direct criterion in evaluating submissions (although there is some evidence that prior publication does directly benefit repeat submitters, e.g., Petersen & Penner, 2014; Siler et al., 2022). Examination of the *Nature* and *Science* naming papers in this dataset reveals that around 10% of researchers who (co)authored at least one of these papers have (co)authored two or more (Fig. 35). Based on unique ORCIDs, there are several researchers listed on more than 10 naming papers in these journals in this dataset. There is also a skew for more repeat authors to have published multiple times in *Nature* (which has a higher sample size of naming papers; Fig. 36; Table 3). Presumably, prior publication in one high-JIF journal also confers an advantage for subsequent first-time submission to another high-JIF journal (e.g., prior publication in *Nature* probably confers some advantage for a future submission to *Science*).

**Figure 35.**
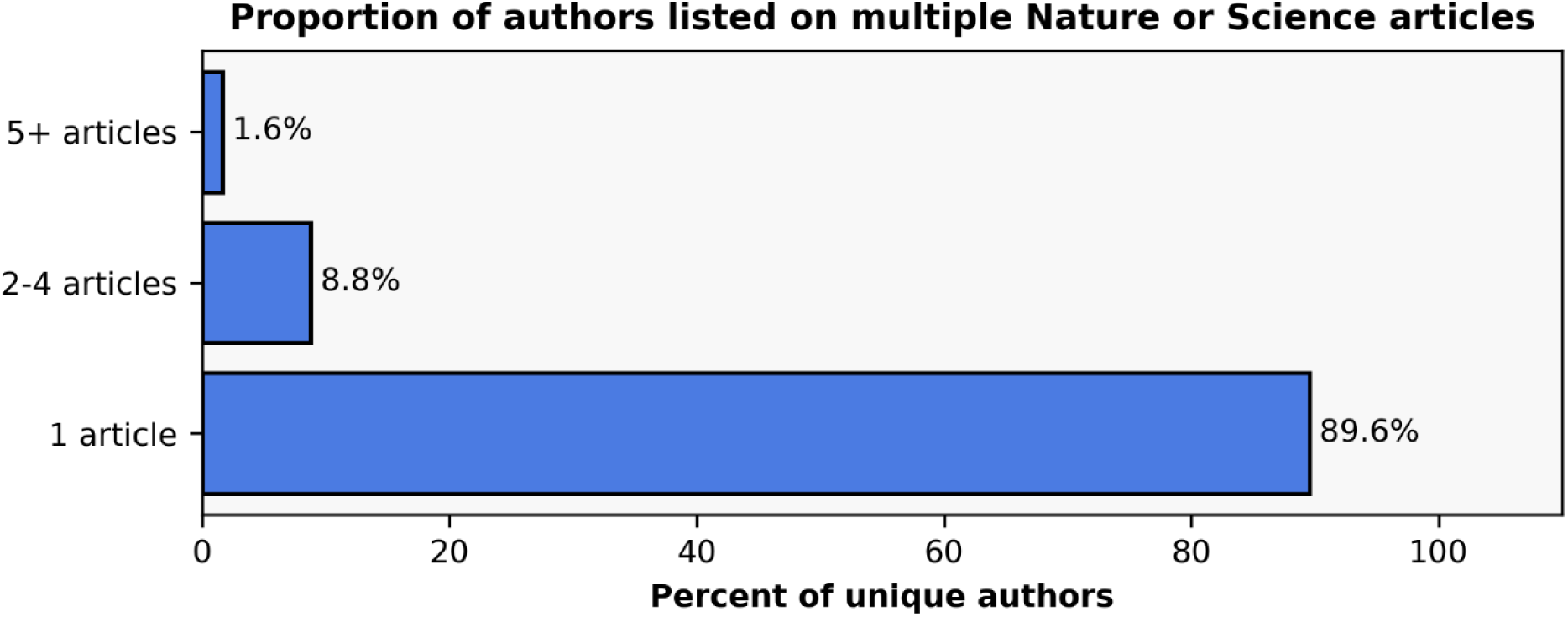
Comparison of the proportion of authors who have multiple naming papers in *Nature* or *Science* in this dataset. This calculation only uses author entries that had linked ORCIDs (*n* = 863 of 1,083 unique names) to minimize issues with shared names, so some author entries are not represented; examination of proportions based on name-only did not indicate substantial differences from those based on ORCID-only. This calculation groups researchers across the two journals, so a researcher with one *Nature* publication and one *Science* publication in this dataset is counted as ‘2-4 articles.’

**Figure 36.**
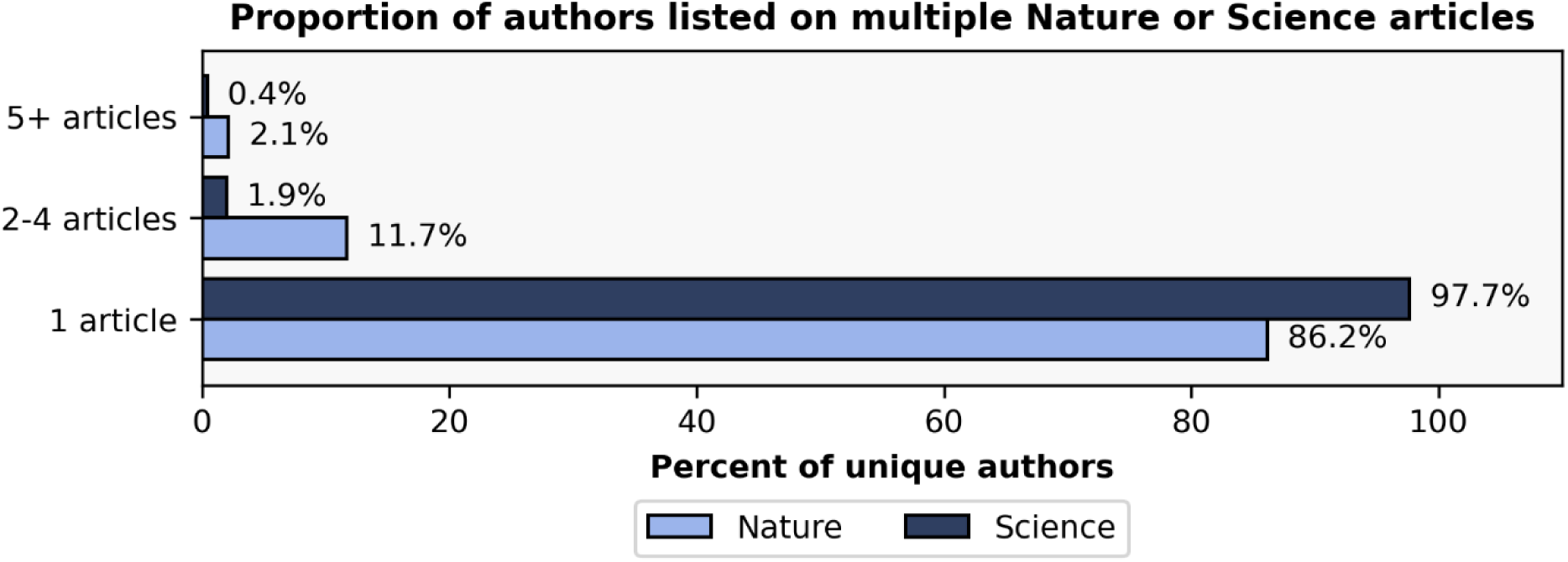
Comparison of the proportion of authors who have multiple naming papers in *Nature* or *Science* in this dataset. This calculation only uses author entries that had linked ORCIDs (*n* = 863 of 1,083 unique names) to minimize issues with shared names, so some author entries are not represented; examination of proportions based on name-only did not indicate substantial differences from those based on ORCID-only. This calculation also parses between journals, so a researcher with one *Nature* publication and one *Science* publication in this dataset is counted as ‘1 article’ for both, not ‘2-4 articles.’

The final consideration is how interpersonal dynamics between specialists of charismatic taxa may lead to biases within specific topics. Studies have shown that researchers of charismatic extant taxa tend to exhibit more problematic behaviors related to monopolization and territoriality around the research topic (e.g., Tensen & Teske, 2025; Valdez et al., 2025). As topic selection by researchers is in part driven by social dynamics and experiences that may lead to non-random exclusion of some demographics from certain topics (e.g., Gould & Valdez, 2022; Xiang et al., 2025), domination of the research space (e.g., of a study system or locality) by certain workers could constrain the space available for other researchers. For example, because authors of articles in high-JIF journals significantly bolster their reputation through this publication, they may also be more likely to serve as reviewers for submissions to such journals and therein to have the potential to act as a gatekeeping mechanism for work on their topic.

Further investigation of authorship patterns of paleontological studies published in high-JIF journals is warranted but likely would benefit from expanded sampling to increase topical coverage (e.g., all studies with a primary focus on descriptive anatomy) and therein to be able to increase journal coverage (e.g., some journals omitted here due to very low naming paper representation, like *Systematic Biology*). Some of the authors of *Nature* and *Science* naming papers, both repeat and non-repeat in this specific context, have additional publications in these journals and other high-profile venues that do not formalize new species.

## Conclusion

This study has demonstrated marked taxonomic disparity among naming papers for extinct taxa, with clades widely regarded as charismatic, such as mammals and avemetatarsalians, being proportionately overrepresented in high-JIF titles such as *Nature* and *Science*. Whether this results from selective journals actively rejecting work on less charismatic clades at a higher rate, from researchers of such taxa submitting less frequently to selective journals, or from other mechanisms remains unclear. However, I believe that there is good reason to suspect that taxonomy plays some role through subjective assessment criteria such as whether a study will attract broad readership and attention because of the well-known biases in interest among the broader public and because the general taxonomic identity of a study system is more easily comprehended than many other attributes of research when submissions are initially being screened by journal editors.

A preference for charismatic taxa by journals, especially those that are managed by for-profit companies and that lack a partnering scholarly society where society members act as editors (e.g., Nature Portfolio), would not be surprising. Over half of all peer-reviewed articles are published by the “big five” for-profit companies (Elsevier, SAGE, Springer Nature, Taylor & Francis, Wiley), who generate 30–40% profit margins that are largely unrivaled in any industry (e.g., Larivière et al., 2015; Butler et al., 2023, 2024). Given their for-profit nature, these publishers certainly have a vested interest in publishing content that attracts not only broad academic readership but also broad non-academic attention (e.g., press coverage of new dinosaur species). Ostensibly, this explains many of the results of this study and why *Nature* (Hutchinson & Garcia, 2002; Erickson et al., 2004; Xu et al., 2004, 2006, 2012; Voris et al., 2025b; Zanno & Napoli, 2025) and *Science* (Schweitzer et al., 2005a, 2005b, 2007; Erickson et al., 2006; Sereno et al., 2009; Benson et al., 2010; Brusatte et al., 2010; Marshall et al., 2021; Griffin et al., 2025) have published more papers focusing exclusively on tyrannosauroids (a majority of which specifically involve *Tyrannosaurus* Osborn, 1905) in the past quarter-century than they have ever published on most higher clades; this does not include papers that include tyrannosauroids as one of several focal clades (e.g., Organ et al., 2008 [*Science*]).

Taxonomic bias in publishing, regardless of the causal mechanism(s), is more than just an epistemologically interesting phenomenon. The value of a paleontology as a primarily “basic”/”foundational” discipline has long been a challenge for workers seeking a career in the field (e.g., Plotnick et al., 2025). Across the biological sciences, the progressive de-valuation of taxonomic work and taxonomic expertise (“taxonomic impediment”) is well-trodden ground (e.g., Buyck, 1999; Drew, 2011; Wägele et al., 2011; Engel et al., 2021; Löbl et al., 2023), but most of this work has only addressed the issue at a large scale, not with respect to specific lower clades (e.g., Green, 1998; Coleman, 2015) or with respect to specific professional activities such as publication in high-JIF journals.

In light of the documented disparity between and within extant and extinct clades of organisms in this study and for extant systems, there is good reason to suspect that career threats to paleontologists with taxonomic focuses are not uniform across clades of specialization. Due to the pervasiveness of metrics-based research assessment, which includes journals’ branding, reputation, and JIF, it is easy to conceive of a scenario in which researchers who specialize in charismatic taxa have an advantage in “selling” their work to high-impact journals and, in turn, researchers who publish in such journals, a small minority of the field and specializing in a small part of the Tree of Life, would acquire significant cumulative advantage (the ‘Matthew Effect’; e.g., Merton, 1968, 1988; Petersen et al., 2010; Ebadi & Schiffauerova, 2016; Petersen & Penner, 2014; Bol et al., 2018; Katchanov et al., 2023) towards being awarded significant funding, being hired into prestigious positions, and being successful in future submissions to a high-profile journal. At the same time, this would build a cumulative disadvantage for workers of less charismatic taxa. There are no clear benefits to such disparity, but the potential consolidation of academic resources, positions, and focus toward work on a select few clades and by a relatively select few workers inhibits our field’s ability to advance a holistic understanding of the history of all life on this planet and to convey that to the broader public.

## Data and code availability

The dataset associated with this version of the preprint is deposited on Zenodo (Gee, 2026b), the scripts associated with this version of the preprint are deposited on Zenodo (Gee, 2026a), and the supplemental appendices are deposited on Zenodo (Gee, 2026c). The actively developed codebase is also available on GitHub (https://github.com/bryanmgee/new-taxa-in-fancy-journals).

## Acknowledgements

Thanks to Emily Troyer, whose Bluesky post (Troyer, 2026) inspired the undertaking of this project. Chris Griffin, Russell Engelman, and Ben Kligman also provided constructive discussion and encouragement to expand this project beyond a one-week undertaking. Molly Hirst helped with finding some papers and in providing feedback on some sections of the draft. Crossref, OpenAlex, and the Paleobiology Database are thanked for making their APIs publicly and freely accessible. No funding was received in support of this work.

